# Chloroplast nucleoids are highly dynamic in ploidy, number, and structure during angiosperm leaf development

**DOI:** 10.1101/632240

**Authors:** Stephan Greiner, Hieronim Golczyk, Irina Malinova, Tommaso Pellizzer, Ralph Bock, Thomas Börner, Reinhold G. Herrmann

## Abstract

Chloroplast nucleoids are large, compact nucleoprotein structures containing multiple copies of the plastid genome. Studies on structural and quantitative changes of plastid DNA (ptDNA) during leaf development are scarce and have produced controversial data. We have systematically investigated nucleoid dynamics and ptDNA quantities in mesophyll of *Arabidopsis*, tobacco, sugar beet, and maize from the early post-meristematic stage until necrosis. DNA of individual nucleoids was quantified by DAPI-based supersensitive epifluorescence microscopy. Nucleoids occurred in scattered, stacked or ring-shaped arrangements and in recurring patterns during leaf development remarkably similar between the species studied. Nucleoids per organelle varied from few in meristematic plastids to >30 in mature chloroplasts (corresponding to about 20-750 nucleoids per cell). Nucleoid ploidies ranged from haploid to >20-fold even within individual organelles, with average values between 2.6- and 6.7-fold and little changes during leaf development. DNA quantities per organelle increased gradually from about a dozen plastome copies in tiny plastids of apex cells to 70-130 copies in chloroplasts of about 7 μm diameter in mature mesophyll tissue, and from about 80 plastome copies in meristematic cells to 2,600-3,300 copies in mature diploid mesophyll cells without conspicuous decline during leaf development. Pulsed-field electrophoresis, restriction of high-molecular weight DNA from chloroplasts and gerontoplasts, and CsCl equilibrium centrifugation of single- and double-stranded ptDNA revealed no noticeable fragmentation of the organelle DNA during leaf development, implying that plastid genomes in mesophyll tissues are remarkably stable until senescence.

**Significance Statement:** Plastid DNA is organized in nucleoids that are highly dynamic in organization, structure and amount during leaf development. The present investigation fully resolves now this dynamic and is a precise cytogenetic characterization of nucleoids DNA spanning the entire life cycle of the leaf.

## Introduction

The plastid genome (plastome; Renner, 1934) represents one of three spatially separated cellular subgenomes constituting the genetic system of plants. The compartmentalized eukaryotic genomes operate as a functional unit, forming an integrated co-evolving genetic system, in which the expression of the dispersed genetic information is tightly adjusted in time, space, and quantitatively (Herrmann, 1997, Bock, 2007, Greiner *et al*., 2011). Basic cellular functions that are indispensable for growth, development and reproduction, including gene expression, photosynthesis, various other metabolic pathways and cell division, depend on the interplay of the genetic compartments (Bock, 2007).

An intriguing characteristic distinguishing the plastome from the nuclear genome is its high copy number per organelle and cell. This redundancy explains much of the non-Mendelian pattern of plastid inheritance, including somatic segregation and transmission of plastid-encoded traits to the next generation. By contrast, the functional significance and persistence of the high plastome copy numbers throughout leaf and plant development are not fully understood. Already from early work, it became evident that both the degree of the plastome reiteration and the ratio of nuclear to organellar genomes, the cellular subgenome homeostasis, are highly variable, can change with development, tissue and nuclear ploidy, and appear to be relatively stringently adjusted by at least two counteracting processes that operate to change or maintain genome-plastome ratios (Butterfass, 1979, Herrmann and Possingham, 1980, Rauwolf *et al*., 2010, Liere and Börner, 2013). The multiple copies of the plastid genome are condensed in nucleoids that reside in the stroma and exhibit prokaryotic properties, consistent with the cyanobacterial ancestry of the plastid (reviewed in Herrmann and Possingham, 1980, Sakai *et al*., 2004, Powikrowska *et al*., 2014). In the leaf mesophyll, the development of chloroplasts from undifferentiated proplastids present in meristems is accompanied by an increase of plastids in both size and number per cell (cf. Butterfass, 1979). This includes a substantial increase in nucleoid number and plastome copies per cell, while nuclear DNA amounts remain constant (e.g., Herrmann and Kowallik, 1970, Selldén and Leech, 1981, Boffey and Leech, 1982, Hashimoto, 1985, Miyamura *et al*., 1986, Baumgartner *et al*., 1989, Miyamura *et al*., 1990, Fujie *et al*., 1994, Rauwolf *et al*., 2010, Golczyk *et al*., 2014, Ma and Li, 2015).

Although numerous studies have suggested that the spatial organization of DNA in chloroplasts of mature leaf tissue is comparable for quite a wide range of seed plants (e.g., James and Jope, 1978, Kuroiwa *et al*., 1981, Golczyk *et al*., 2014), our knowledge about the localization, structural organization and quantity of plastid DNA (ptDNA) is rather fragmentary. The available information is restricted to a limited number of species and relatively few (often barely comparable) developmental stages, tissues or conditions. Developmental patterns in shape and arrangement of nucleoids have not been systematically studied. Furthermore, reports on fundamental aspects such as DNA quantities per organelle or cell, their dynamic changes, and the maintenance or degradation of ptDNA during tissue maturation are highly controversial, thus adding to the confusion. Reliable quantitative data are almost entirely lacking. DNA amounts reported for fully developed chloroplasts span almost three orders of magnitude, from less than half a dozen (Pascoe and Ingle, 1978) to 1,000 or more copies (e.g.,Boffey and Leech, 1982, for further references see Rauwolf *et al*., 2010, Liere and Börner, 2013). While there are consistent data about the degradation of ptDNA in senecescing tissues (Sodmergen *et al*., 1989, Inada *et al*., 1998, Golczyk *et al*., 2014) under participation of the exonuclease DPD1 (Takami *et al*., 2018), the question whether ptDNA in leaves is indeed lost or substantially reduced already relatively early during leaf development (e.g., Rowan *et al*., 2004, Oldenburg *et al*., 2006, Shaver *et al*., 2006, Rowan *et al*., 2009), or is rather kept at high copy numbers and remains functionally active until senescence, needs to be resolved (e.g., discussion in Golczyk *et al*., 2014, Oldenburg *et al*., 2014, Sakamoto and Takami, 2018, Takami *et al*., 2018). Comparably, it needs to be clarified whether or not plastid genes and genomes are inactivated by mutations and degraded to non-functional fragments in mature, photosynthetically active mesophyll cells (Kumar *et al*., 2014, Oldenburg *et al*., 2014, Kumar *et al*., 2015) or remain intact (e.g., Ma and Li, 2015). Obviously, the intense debate about loss, inactivation or retention of ptDNA during leaf development or under certain conditions has precluded deducing a meaningful view of the cellular basis of the plastome during development.

To resolve this controversy, and to provide complete datasets about the fate and amounts of the ptDNA including the dynamics of plastid nucleoids during the entire leaf development, we set out to comprehensively investigate ptDNA in mesophyll cells from early post-meristematic tissue until late senescence. Since the contentious findings reported in the literature were obtained with comparable material, often from the same species, it is evident that they reflect deficits in the methodology and/or experimental artifacts. Therefore we have studied quantity, arrangement and integrity of DNA in plastids of mesophyll cells over the entire developmental cycle of leaves in four higher plant species (*Arabidopsis thaliana, Beta vulgaris, Nicotiana tabacum* and *Zea mays*), employing an array of independent techniques, including advanced supersensitive DAPI-based epifluorescence microscopy, real-time quantitative PCR with total DNA from leaves as well as from mesophyll protoplasts, pulsed-field gel electrophoresis, restriction of high-molecular weight DNA from chloroplasts and gerontoplasts, and ultracentrifigation of single- and double-stranded ptDNA in analytical CsCl equilibrium gradients. Especial care was taken determining ptDNA amounts. Fluorescence emissions of individual nucleoids, for instance, were quantified relative to that of T4 phage particles (that served as a haploid standard) in high-resolution images obtained by integrating (3D) records systematically taken within seconds at consecutive vertical focal levels along the z-axis across entire organelles into 2D projections. Compared to conventional approaches this technique avoids the problem of pattern variation with changes of focal plane (see e.g., James and Jope, 1978, Hashimoto, 1985, Golczyk *et al*., 2014), results in superior optical resolution and image sharpness, and allows both more precise localization and accurate quantification of ptDNA. In a previous study, we analyzed mesophyll tissue from nearly mature to necrotic leaves (Golczyk *et al*., 2014). In this work, we have focused predominantly on early leaf development, covering the transition from the meristematic and early post-meristematic stages to maturity. During this developmental process, leaves convert from sink to source organs and their plastids undergo profound changes. We have addressed quantitative and morphological aspects of ptDNA organization in mesophyll cells over the entire developmental cycle and discuss our findings in the light of the controversies about stability and integrity of the chloroplast DNA in leaf development.

## Results

### Nucleoid patterns in plastids during early leaf development

Figures 1 and 2 show representative photomicrographs of a developmental series of DAPI-stained mesophyll cells from sugar beet, *Arabidopsis*, tobacco and maize ranging from meristematic/post-meristematic to post-mature leaf tissue. A more comprehensive developmental record is presented in Data S1 - S4 (panels 1 - 128 for sugar beet, panels 129 - 271 for *Arabidopsis*, panels 272 - 330 for tobacco, and panels 331 - 384 for maize). As mentioned above the photomicrographs shown represent projections of combined 3D records across entire individual organelles, visualizing the nucleoids from the different focal planes of an organelle in a single image (see Discussion). Explants, leaflets and leaves from which samples were taken are described in Material and Methods, some examples are photographically documented in Golczyk *et al*. (2014), and for sugar beet, also in Rauwolf *et al*. (2010).

**Figure 1.**
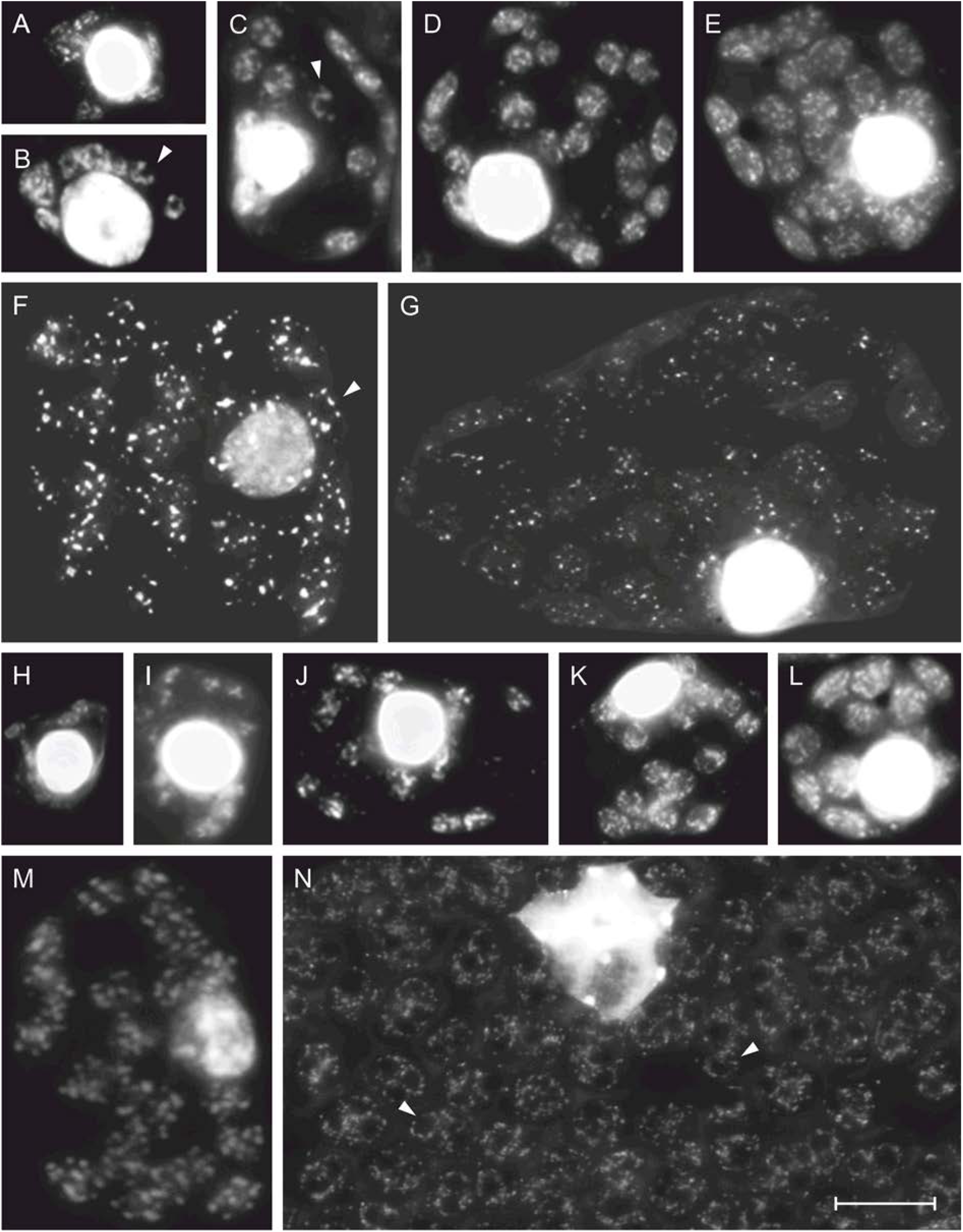
DAPI-DNA fluorescence photomicrographs of mesophyll cells from *Beta vulgaris* (a-g) and *Arabidopsis thaliana* (h-n). [(a), (b), (h), (i)]: representative DAPI-stained cells of squashed meristematic leaflet tissue, [(c-e), (j-l)]: juvenile mesophyll cells, and [(f), (m)]: cells of premature mesophyll. (g): diploid cell of mature mesophyll, (n) nuclear region in a (probably polyploid) postmature mesophyll cell; the entire cell is shown in panel 271 of Data S2. Arrowheads in (b, c, f and n) mark circular nucleoid arrangements. Bar = 10 μm.

**Figure 2.**
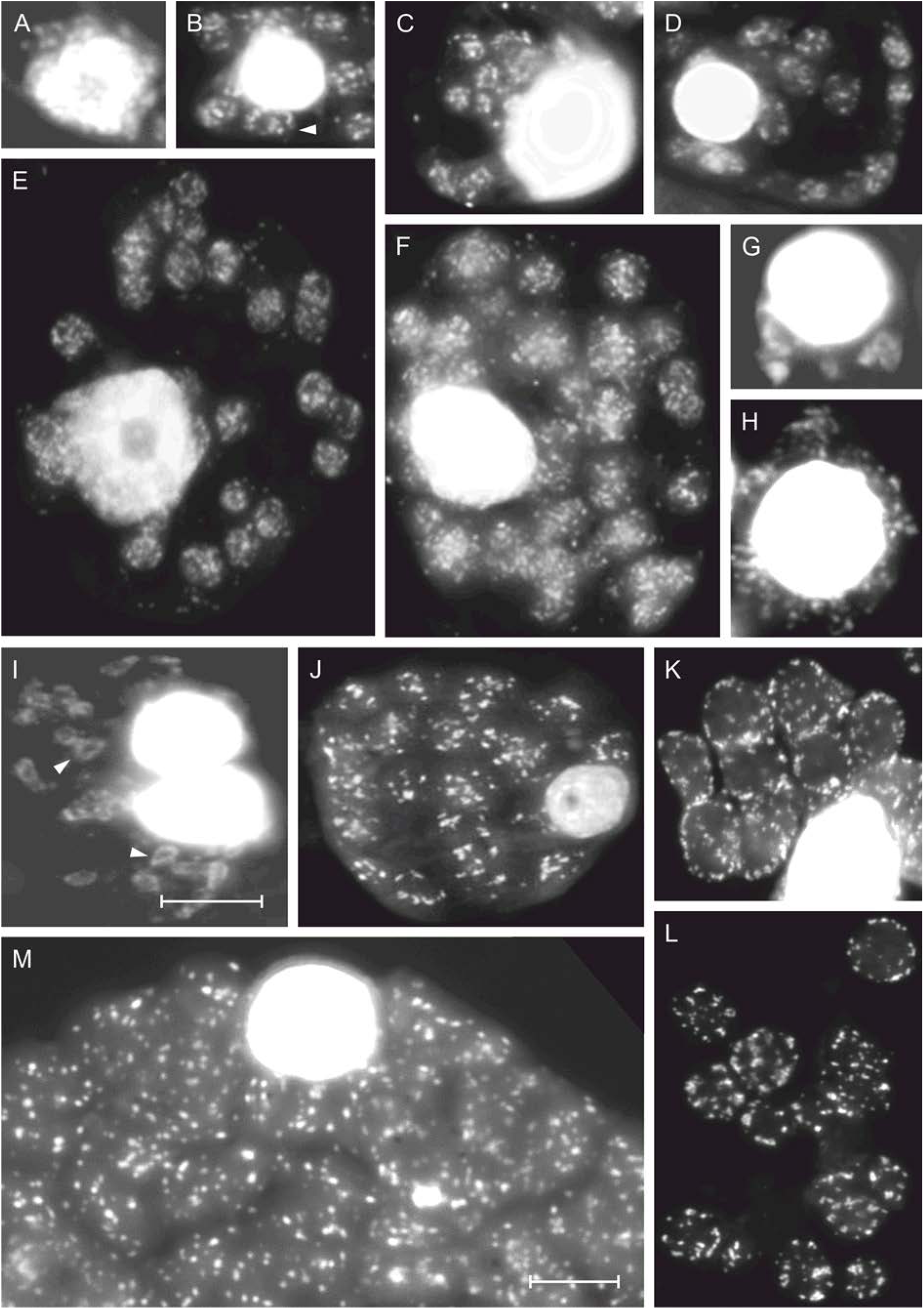
DAPI-DNA fluorescence photomicrographs of mesophyll cells from *Nicotiana tabacum* (a-f) and *Zea mays* (g-m). [(a), (g), (h)]: representative DAPI-stained cells of squashed meristematic leaflet tissue, (b-e) of juvenile mesophyll tissue, (f, j) of premature and (m) of diploid mature mesophyll. Arrowheads in (b) and (i) mark circular nucleoid arrangements. (k and l): transition stages from ring-shaped [k] and intermediate ring-shaped to disperse (l) nucleoid arrangements in premature tissue. Entire cells illustrating this nucleoid pattern are presented in panels 374 - 380 of Data S4. Bars (a-i) and (j-m) = 10 μm.

Significant DNA fluorescence in plastids could be discerned during all stages of leaf development. Nucleoids were clearly visible within the organelles as distinct fluorescing spots that were scattered virtually randomly in almost all matrix areas. An individual spot may traverse several planes, either as individual or stacked nucleoids (cf. Herrmann and Kowallik, 1970), and there was substantial nucleoid heterogeneity in and between individual organelles (see below). In young leaf material, fluorescence occasionally appears somewhat diffuse, presumably due to the 2D projection of the spatial records of densely packed nucleoids. Analysis of meristematic and early post-meristematic cells was sometimes difficult, because the cytoplasm adhered tightly to the strongly stained nucleus. The intensity of nuclear staining was locally so high that it outshined plastid fluorescence, thus preventing adequate photographical documentation of nucleoids at normal exposure times. Leaf development was accompanied by spatial changes of nucleoid patterns, which exhibited remarkable similarity among the species studied. The overall findings for the early stages of leaf development are based on the analysis of about 1,300 cells and 3,760 chloroplasts. They are briefly summarized below, documented in the Figures and Supplementary Datasets mentioned above, and summarized in Table 1. Complementary information is presented in Appendix S1. Figure 3 presents schematically the major changes in nucleoid morphology and distribution patterns in mesophyll plastids during leaf development, as detected by fluorescence microscopy.

**Figure 3.**
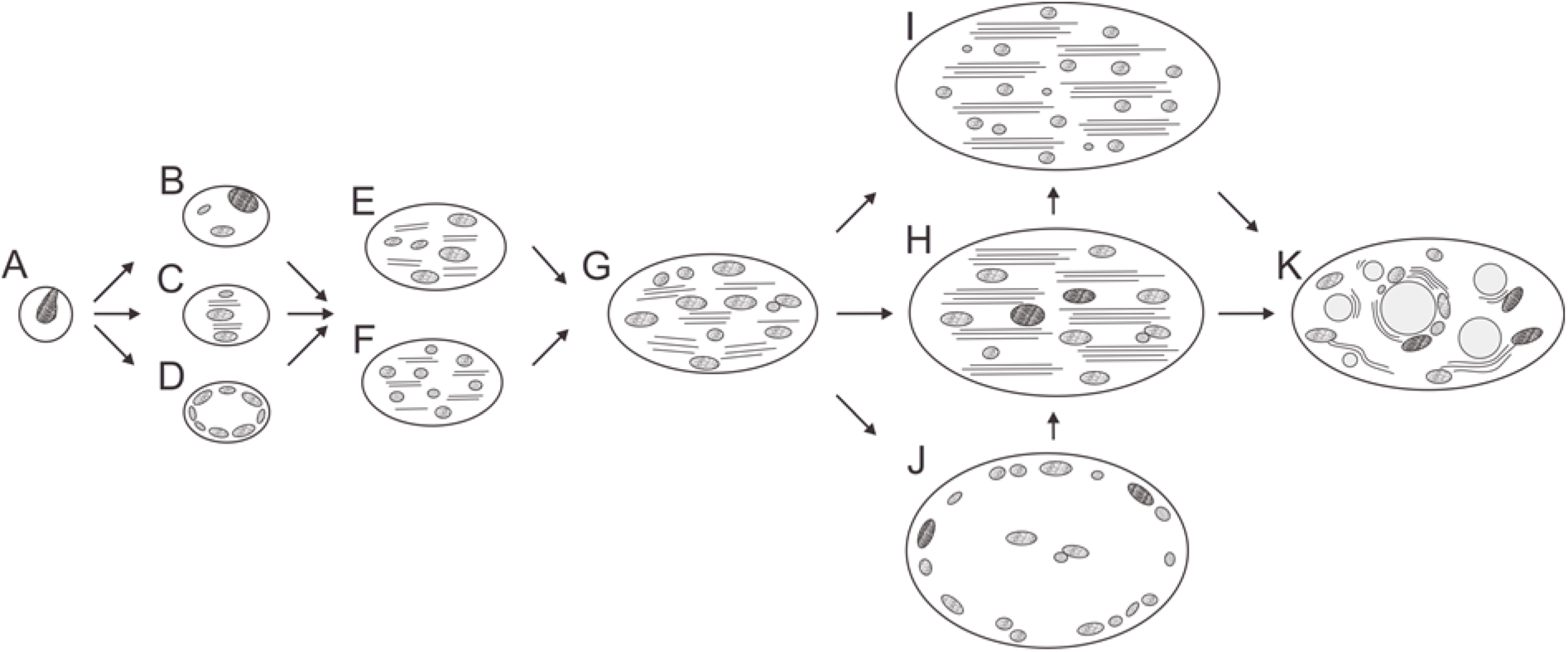
Model illustrating the major spatial changes and the patterns of nucleoids occurring in plastids during development from embryonic to senescent leaf mesophyll tissue, as deduced from DAPI-based fluorescence microscopy and three-dimensional reconstructions from ultrastructural serial sections through stroma-depleted organelles (Herrmann and Kowallik, 1970, Kowallik and Herrmann, 1972). (a) The tiny plastids present in meristems are proplastids or leucoplasts of ≤1 μm in diameter that harbor a single nucleoid. (b-d) Plastids with a few nucleoids that either lie side-by-side (b), become stacked and are interspersed by emerging thylakoid layers (c), or form ring-shaped arrangements along the organelle periphery, with a necklace-like knotty or stringy appearance (d). (e, f) Developing chloroplasts from juvenile material harbor either a limited number of brightly fluorescing DNA spots (e) or a larger number of smaller, densely packed DNA spots (f). (g) Mature chloroplasts usually contain approximately 30 discrete DNA spots. The organelles may enlarge without significant increase in DNA content either without (h) or with (i) accompanying nucleoid division, or display transitory circular nucleoid arrangements (j). (k) Gerontoplasts with plastoglobule are morphologically diverse and can also harbor circular nucleoid arrangements (see also Golczyk *et al*., 2014). Note that variation in nucleoid ploidy is not fully represented in this drawing. Organelles, nucleoids or membranes are not drawn to scale. The model likely applies to many vascular plant species.

**Table 1.**
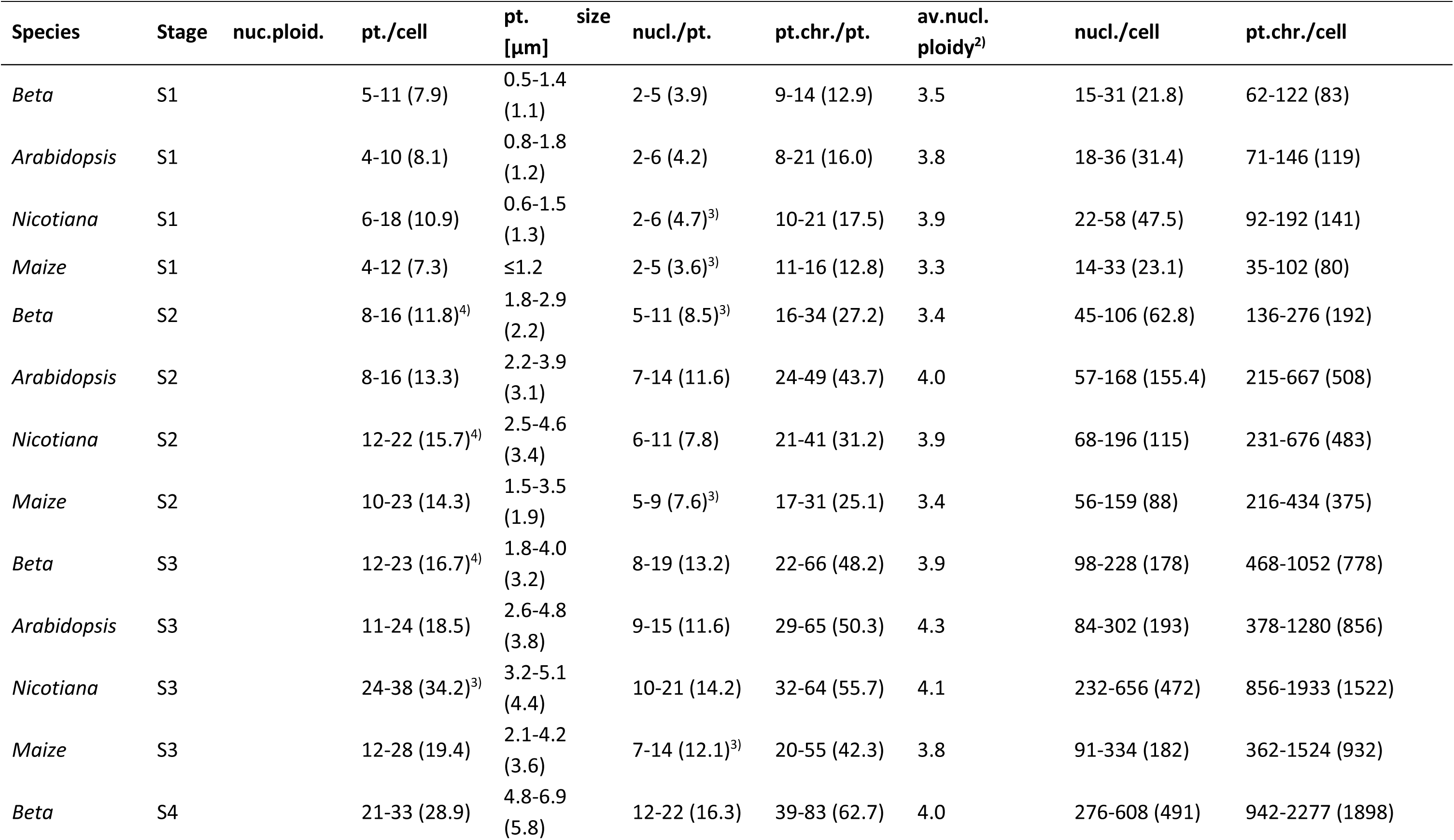
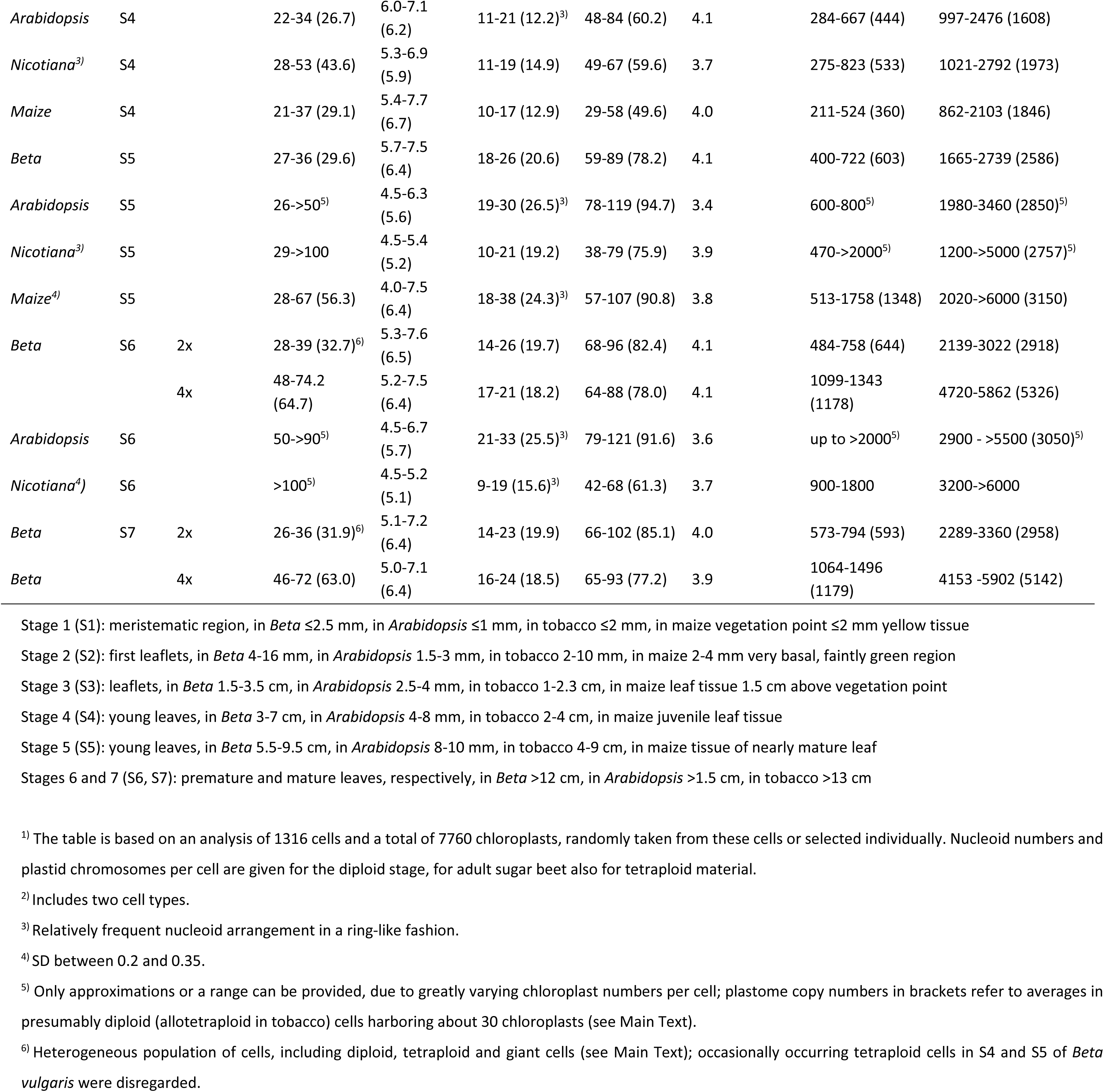
Structural and Quantitative Fluorescence Microscopic Analysis of ptDNA in the Mesophyll of Four Higher Plant Species^1)^

During early mesophyll development from leaf stages 1 - 5 (cf. Material and Methods), cell size, number and size of plastids as well as nucleoid number per organelle increased continuously, as expected. The round-shaped cells enlarged and elongated, the diameters of the organelles expanded from about 1 μm in meristematic/postmeristematic tissue to about 7 μm in premature/mature leaves, corresponding to an about 60-fold increase in plastid volume. Within this time frame, plastid numbers per cell increased from 4 - 8 to 30 - 35 in mature (diploid) cells, and nucleoid numbers rose from 2 - 4 to approximately 25 - 35 per organelle.

*Stage 1:* In meristematic and early post-meristematic leaf tissue, the DNA of the nucleoids replicates, nucleoids divide and segregate into a few spherical, ovoid or oblong DNA-containing bodies that lie side-by-side, are stacked, or are arranged peripherally in a circular fashion (Figure 3a,d, Figure 1a,b,h, and i, Figure 2a,g, and h, Data S1 - S4, panels 1 - 52, 129 - 162, 272 - 283, 331 - 348). Organelles with only a single nucleoid were rare.

*Stages 2-3:* In juvenile tissue of sugar beet and maize, the organelles usually remain relatively small (2 - 3 μm in diameter) and contain a limited number (typically 7 to 14) of scattered DNA spots (Figure 3e, Figure 1c,d, and e, Figure 2b,c, and i, e.g. Data S1 and S4, panels 53ff and 349 for sugar beet and maize, respectively, see also Golczyk *et al*., 2014). In *Arabidopsis* (Data S2, panels 163 - 228) and accompanying incipient cell elongation in tobacco (Data S3, panels 293 - 309), cells and chloroplasts became larger, organelles changed position and accumulated at the cell surface (thus replacing the more or less scattered distribution seen before), and synthesized relatively large amounts of DNA, as judged from high numbers of densely packed DNA spots, which were often impossible to resolve and to count individually (e.g., Figure 1k,l, and m, but see j, Figure 2e,f, Figure 3f, Data S2 and S3, e.g., panels 178ff, 300ff). The heterogeneity of the cells and organelle populations observed indicates intense developmental activity during these and the subsequent stages. In general, the dispersed spotty pattern of nucleoids still prevailed, but ring-like, occasionally asymmetric or elongated half-moon-like arrangements occurred quite often (e.g., Figure 3d-f, Figure 1b,c Figure 2i, Data S1 - S4, e.g., panels 21, 68, 71, 85 - 87, 89, 166, 197, 212, 220, 227, 268, 271, 299, 302, 312, 317, 358, 362. 363, 365, 370).

*Stages 3 - 4:* In elongated cells, chloroplasts were usually tightly packed side-by-side at the cell surface. They contained numerous nucleoids (15 -> 20; e.g., Fig. 1f, Fig. 2f and j, Data S1 and S2, e.g., panels 107ff, 251ff, see also Golczyk *et al*., 2014), but were still not fully expanded (Figure 3g). Organelles bearing fewer nucleoids (8 - 15) were observed, notably again in sugar beet and maize (e.g., Figure 3e,h, Figure 1f,j). In those instances, nucleoid fluorescence emission was generally brighter.

*Stages 4 - 5:* In pre-mature leaves with lamina extensions up to about 9.5 cm in sugar beet and tobacco, and 4 - ≥8 mm in *Arabidopsis*, cell sizes (40 - 50 µm), plastid numbers and sizes in mesophyll tissue approach the means found in mature diploid leaves. In sugar beet, *Arabidopsis,* tobacco and, to some extent, in maize plastid numbers per cell were typically in the range of 25 - 35 (but occasionally ≥45). Chloroplasts were 5 - 7.5 µm in diameter and harbored 14 to >30 usually dispersed nucleoids (the average being approximately 23; e.g., Figure 3h, Figure 2m). Circular nucleoid arrangements were noted again, especially in maize, but were also quite abundant in *Arabidopsis* and tobacco (Figure 3j, Figure 1n, Figure 2k and l, Figure 3j, Data S1 - S4, e.g., panels 270, 271, 328, 329, 374 - 380; in “giant” cells: Data S5, panels c and e).

Relatively large cells (60 - 80 µm) with higher, approximately doubled chloroplast numbers (60 - 70) and larger nuclei appeared as the leaf reached maturity, and probably reflect somatic endopolyploidization (rather than the G2 cell cycel phase; Butterfass, 1979 e.g., Data S1, e.g., panels 128, 271, Data S8, panels a, d, f, g, and j). Chloroplast sizes and nucleoid patterns in diploid and tetraploid cells were indistinguishable, indicating regulation independent of the ploidy level at this stage (see Discussion). In *Beta*, for instance, bimodal size distributions of mesophyll cells were observed at this stage, and the fraction of tetraploid cells increased with leaf age (Butterfass, 1979). An intriguing observation was that chloroplasts in premature to early postmature leaf mesophyll multiply relatively rapidly, without noticeable size changes (and in the absence of cell division). Giant cells with very high and greatly variable organelle numbers were detected in *Arabidopsis,* sugar beet and tobacco, with up to about 150 chloroplasts per cell in *Arabidopsis*, and several hundred in tobacco (Data S5, Data S2, panel 271). Panel (d) in Data S5 illustrates that these cells are clustered and thus do not represent idioblasts. Astoundingly, the chloroplasts displayed rather normal nucleoid patterns, implying significantly elevated ptDNA levels per cell, without much increase in nuclear volume (see Discussion).

Patterns, numbers, shapes and fluorescence emission intensities of nucleoids were not substantially different in chloroplasts of premature, mature or ageing leaves, or in cells differing in ploidy, consistent with previous work (Rauwolf *et al*., 2010, Golczyk *et al*., 2014). This observation indicates that DNA synthesis in plastids largely stops before cessation of cell proliferation, and ptDNA contents per organelle and per cell increase until that stage, but not later (irrespective of endopolyploidization). However, with leaf ageing, chloroplasts (and cells) may expand further, and their DNA can be divided among higher numbers (≥35) of small spots (nucleoids) that are widely scattered throughout the organelle interior (e.g., Data S1 and S2, panels 125, 126, 269; Fig. 1G, Fig. 3I, Golczyk *et al*., 2014). Their significantly lower fluorescence is indicative of nucleoid division without substantial DNA synthesis.

### Quantification of ptDNA per organelle and cell - variation in nucleoid ploidy

To follow the quantitative changes in plastid genome content during leaf development, two strategies were employed determining the amounts of ptDNA: an advanced high-resolution fluorescence densitometry and real-time qPCR. The two approaches are technically independent and thus complement each other. While microfluorimetry allows quantification of ptDNA at the level of individual nucleoids, organelles and cells, qPCR provides approximations of average cellular ptDNA amounts that can be used to calculate mean DNA amounts per nucleoid and plastid.

### Nucleoid diversity

Quantifications based on fluorescence techniques have to take into account the remarkable structural diversity of plastid nucleoids. High-resolution images of DAPI-stained plastids obtained by rapid integration of high-resolution vertical records from different focal planes across an organelle (see Discussion) reveal this variability as well as differences in nucleoid numbers per plastid and a surprising similarity of patterns among the four plant species studied (Figure 4 and Data S6 and S7). The DNA spots were irregular in shape, oblong or spherical, and ranged from approximately 3 μm in length down to the limit of resolution. This variability likely reflects the unequal distribution of the nucleic acid within the organelle stroma and implies substantial ploidy differences between spots. Our estimates suggested that the local DNA concentration can vary by more than an order of magnitude. Mere counts of nucleoids per organelle miss this important feature of ptDNA dynamics during development.

**Figure 4.**
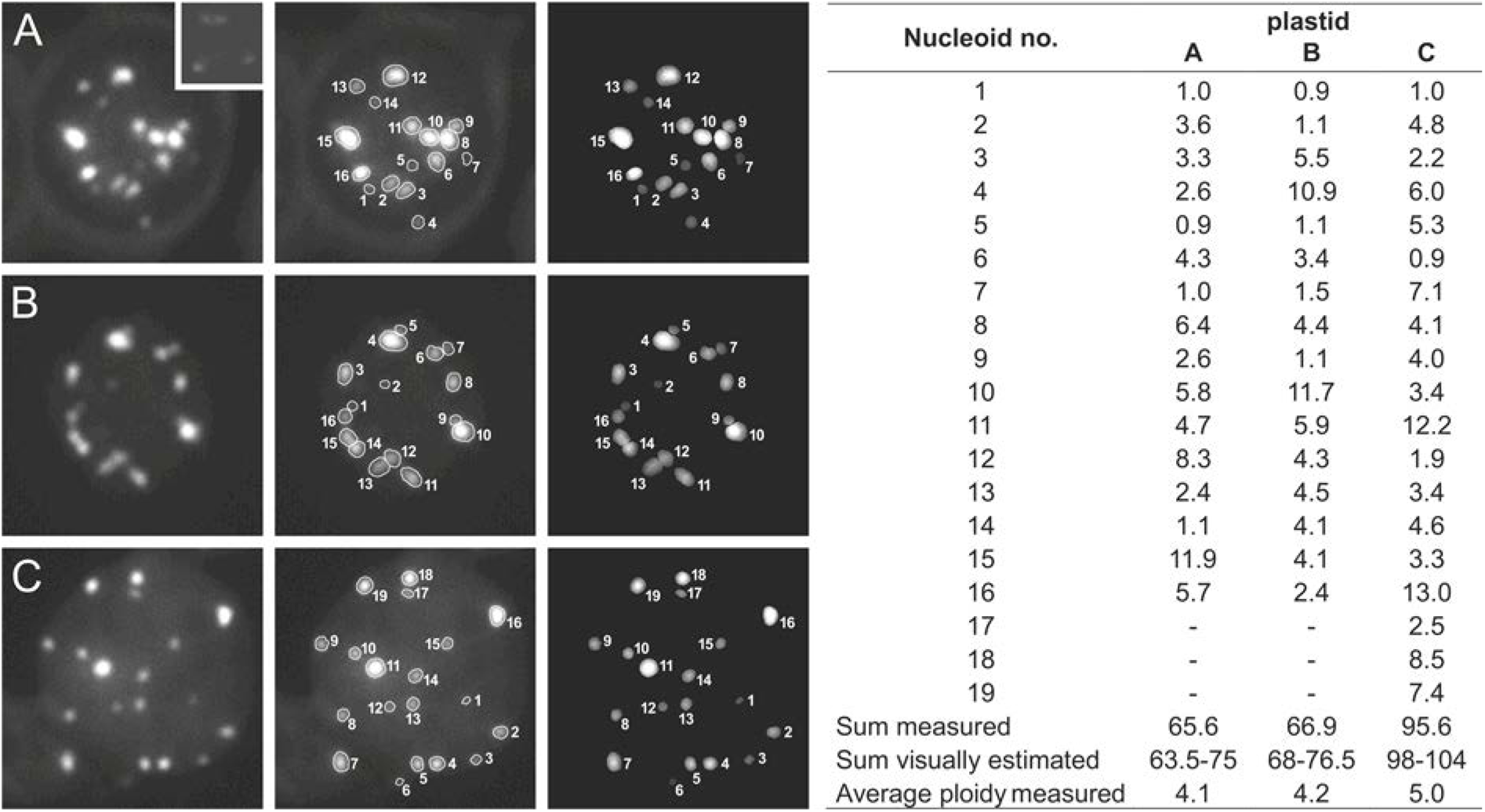
Quantitative high-resolution epifluorescence microphotometry of DAPI-stained individual mesophyll chloroplasts of maize. Organelles from cells of expanding (a), premature (b) and mature (c) leaves with diameters of 3.6, 4.5 and 6.5 µm, respectively, are shown as examples (for additional data see Data S6 and S7). Fluorescence emissions were determined individually for all nucleoids from magnified records (three independent measurement cycles were carried out for each organelle), and normalized to DAPI-stained T4 phage heads (inset in a) that served as a haploid reference corresponding to a single plastome equivalent. Note the considerable fluorescence variation between ptDNA spots. The intensity of the lowest detectable signals in organelles corresponded to that of T4 particles. Data were also analysed visually with a magnifying glass by three independent investigators. To this end, nucleoid fluorescence was compared to a graded series of fluorescence spots of increasing emission intensity quantified *in silico*. The Table lists the determined ploidy values for the individual nucleoids. Nucleoid ploidies ranged from haploid (nucleoids nos. 1, 5, 7 and 14) to about 12-fold (nucleoid no. 15) in the chloroplast shown in (a) (with 16 nucleoids), from haploid (nos. 1, 2, 5 and 9) to about 11-fold (nos. 4 and 10) in the chloroplast in B (with 16 nucleoids), and from haploid (nos. 1 and 6) to about 13-fold (no. 16) in the chloroplast in C (with 19 nucleoids), with average nucleoid ploidy levels being between tetra- and pentaploid. The estimated total number of plastid genome copies per organelle were between >60 and 100. Note that spectrometrically and visually determined values are in good agreement. In approximately 40% of the 42 individual chloroplasts investigated in this experiment, the datasets differed between 7 and 12%, in 45% of the chloroplasts between 13 and 20%, and in the remaining cases up to 27%. As expected, major differences resulted from intensely fluorescing spots that were beyond signal saturation (see Discussion). For further details, see Material and Methods and Main Text.

#### ptDNA quantification based on DAPI-DNA fluorescence

The cytological findings were substantiated by microdensitometric analyses of well separated fluorescing spots in magnified individual plastids and by visual comparison with scales of dots of increasing emission intensity determined *in silico*. To this end, the fluorescence of individual nucleoids in photomicrographs was normalized to DAPI-stained T4 phage particles after background correction (Figure 4 and Data S6). The phage fluorescence corresponded to that of spots with the lowest detectable emission intensity in chloroplasts. Given that the size of the phage genome (168,903 bp; Miller *et al*., 2003) is similar to that of the plastid genome, it is reasonable to assume that these spots are haploid in first approximation, that is, they contain only a single copy of the plastid genome. Consequently, larger and/or brighter fluorescing dots reflect multiple copies of the ptDNA. Measurements were performed individually on all nucleoids of an organelle. Figure 4 and Data S6 show representative examples of quantified nucleoid profiles for individual chloroplasts from young, developing and mature maize, *Arabidopsis,* sugar beet and tobacco mesophyll, and also provide a comparison of densitometrically and visually obtained data.

Based on 1180 organelles investigated, estimates of nucleoid florescence signals ranged from haploid to >20-fold, with averages between 3.0 and 5.7 genomes per nucleoid (calculated by comparison of nucleoid numbers and plastome copy numbers of individual organelles) implying that nucleoids are, on average, tri- to hexaploid. Plastids in juvenile leaf tissue contained 12 - 20 genome copies, and mature chloroplasts 70 - 130 (Figure 4, Data S6 and Table 1). Fluorescence intensities of nucleoids were comparable in plastids of juvenile leaflets, expanded and ageing leaves, although a trend towards lower values was noted in plastids of meristematic tissue and, to a lesser extent, also in plastids of postmature tissues. Virtually no significant intensity differences were found between DNA-containing regions in organelles of different sizes or in chloroplasts of comparable size that reside in cells that differ in nuclear ploidy. Remarkably, there were also no significant differences among the species studied (see Discussion). An example of the overall distribution of nucleoid ploidies in chloroplasts of nearly mature diploid and tetraploid sugar beet mesophyll cells is shown in Figure 5. Plastome copy numbers among individual plastids of a given cell usually differed only moderately. For example, in six organelles per cell that were randomly chosen from five premature mesophyll cells (each harboring about 20 chloroplasts), numbers ranged between 44 - 62 copies per organelle in maize, and 68 - 79 in sugar beet, with averages between 53.7 and 55.4, and 71.5 and 76.3, respectively. Extrapolation to the copy number per cell (by multiplying the average DNA copies per organelle with the corresponding number of plastids per cell) yielded numbers between 40 and 140 copies for meristematic/post-meristematic cells, and between 2,700 and 3,300 copies for (diploid) cells of mature tissue (Figure 4, Table 1 and Data S6). It is noteworthy that microspectrometric values and values obtained by visual assessment for the same sample were in excellent agreement (i.e., within 20% in about 80% of the cases).

**Figure 5.**
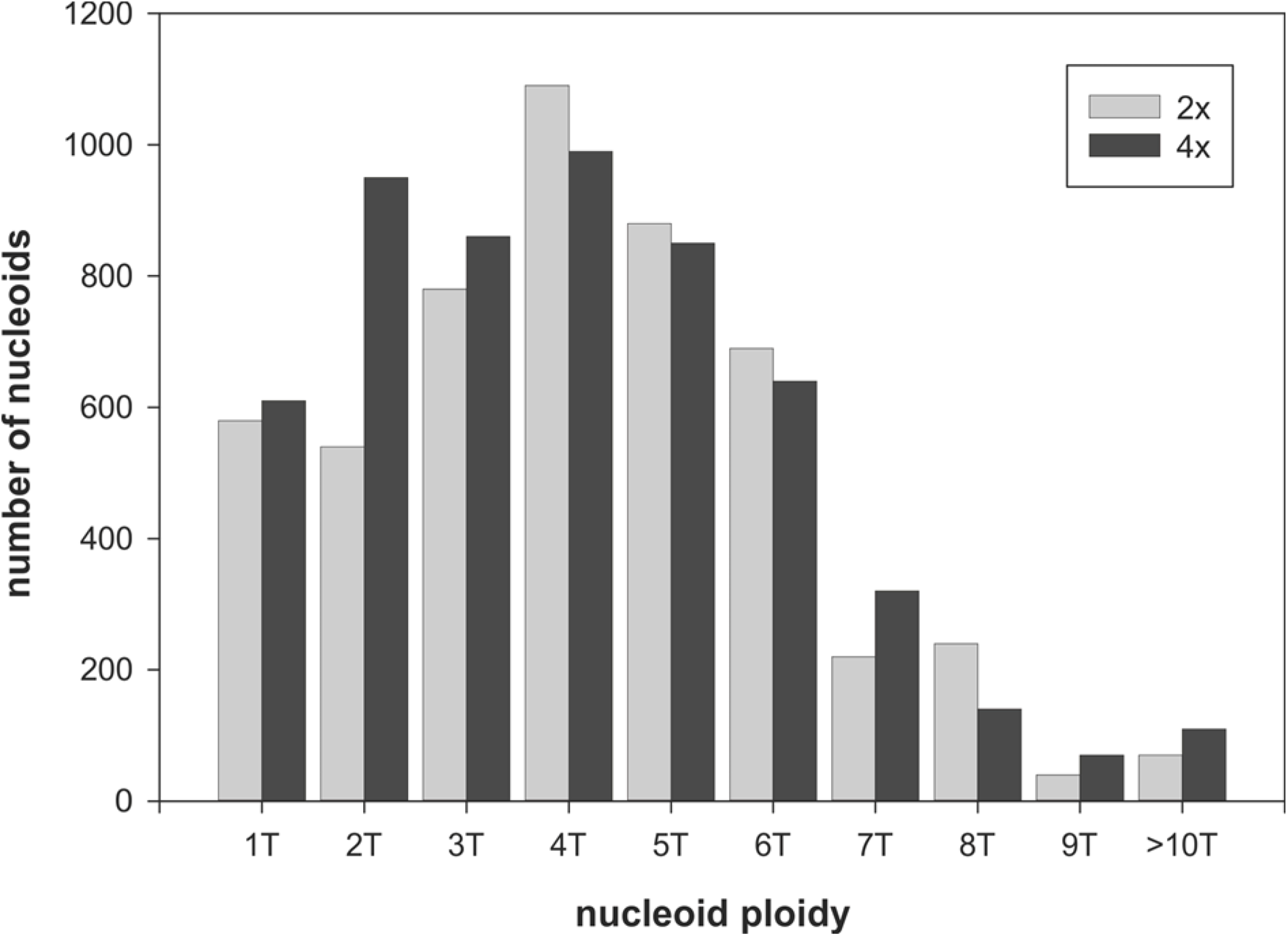
Nucleoid ploidy distribution in mesophyll chloroplasts of premature di- and tetraploid sugar beet leaves determined microspectrometrically and visually from the same staining series. Nucleoid ploidies were normalized to T4 phage head fluorescence and ranged from 1T (haploid) to >10T (>decaploid), the average being 4.3 ± 0.7. The diagram is based on about 450 organelles with an average of 23.2 nucleoids per organelle (corresponding to a total of about 10,400 nucleoids). Data for other developmental stages and plant materials are similar. Somewhat lower ploidies may occasionally be noted for nucleoid spots in organelles of juvenile tissues.

#### Quantitative real-time PCR

qPCR amplified gradually increasing quantities of ptDNA in all species from embryonic to mature stages, which then remained relatively stable in older and advanced senescent tissue (Figure S1, Golczyk *et al*., 2014). It is important to note that the three plastome-specific amplicons selected to be well scattered along the plastid genome yielded comparable results. Although ptDNA values for a given stage may differ somewhat between samples (especially in tissue sampled during the most intense growth period), in all instances, cellular ptDNA levels increased from approximately 100 - 250 plastome copies in meristematic/post-meristematic material to levels in the order of 1,600 - 2,000 copies per diploid cell in mature leaves and subsequent developmental stages.

The PCR-derived values obtained with total leaf DNA were consistently lower than the DAPI-based estimates for mature and ageing tissues, and higher for younger material (see Discussion for possible explanations). In order to assess how non-mesophyll cells and nuclear ploidy influence the estimates, an additional study was conducted with purified mesophyll protoplasts of juvenile, premature and mature leaf tissue from all four species investigated here. Figure 6a-d and Data S8 document the purity of the preparations and confirm that the protoplasts released after pectinase and cellulase treatment were vital (i.e., round-shaped with smooth contours, turgescent and responding osmotically; see Discussion and Appendix S2). Samples from younger tissue contained only low proportions of polyploid cells as judged from the relatively homogenous cell sizes and cellular chloroplast numbers (Butterfass, 1979). As expected, based on the fact that cells in non-green tissues of leaves contain fewer and smaller plastids with less DNA than chloroplasts (reviewed in Liere and Börner, 2013), ptDNA quantities determined per mesophyll protoplast were higher than the corresponding data obtained with total leaf DNA: 1.6 ± 0.1-fold in sugar beet (equivalent to about 2,900 plastome copies per cell), 1.5 ± 0.15-fold in maize and tobacco (about 2,400 to 2,800 copies), and 1.8 ± 0.2-fold in *Arabidopsis* (about 2,750 to 3,100 copies; see Discussion). Mean nucleoid ploidies, calculated as quotients of qPCR values (corrected for non-mesophyll cells and nuclear ploidy) and average nucleoid numbers per organelle, yielded 3.8- to 6-fold higher plastome equivalents than fluorescing spots. These values are in agreement with the copy numbers derived from spectrofluorimetric quantifications (see above) and DNA colorimetry with fractions of isolated weakly fixed plastids from sugar beet (Rauwolf *et al*., 2010).

**Figure 6.**
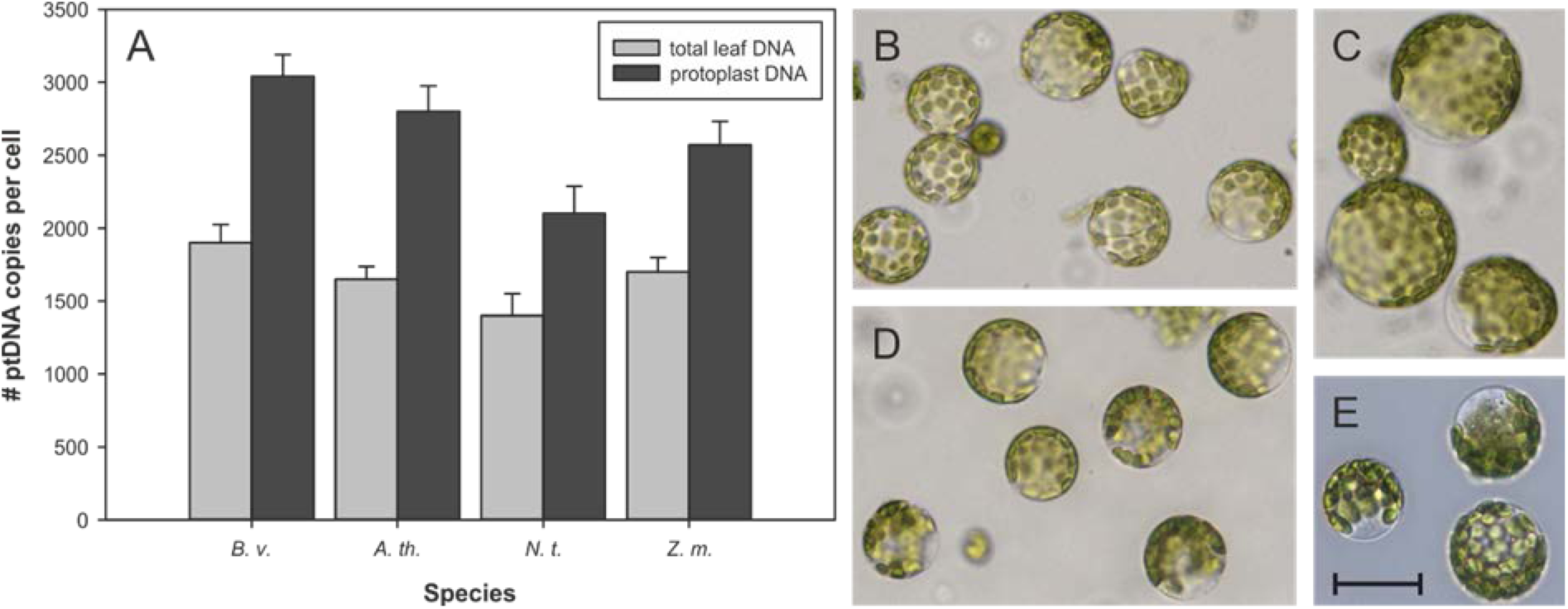
Analysis of plastome copy numbers in mesophyll of sugar beet*, Arabidopsis*, tobacco and maize. (a): Comparison of cellular plastome copy numbers determined by real-time qPCR with total leaf DNA (gray columns) and DNA of protoplasts from premature/mature leaf tissue (black columns; see Methods and Golczyk *et al*., 2014). The graphs show means and standard deviations from four measurements per leaf sample (three replicates per reference gene). For determination of copy numbers and ploidy level adjustments, see Golczyk *et al*. (2014). *B. v.* = *Beta vulgaris*, sugar beet, *A. th.* = *Arabidopsis thaliana*, *N. t.* = *Nicotiana tabacum*, tobacco, and *Z. m.* = *Zea mays*, maize. (b-e): Examples of purified mesophyll protoplasts from premature leaves of sugar beet (b) and *Arabidopsis* (d), and of mature leaves from tobacco (c) and maize (e). All protoplasts exhibit the typical round shape and smooth contours of turgescent, viable cells (see Discussion and Supplemental Text 2). No other leaf cell types contaminated notably the preparation of photosynthetic cells. Note the relatively homogeneous cell sizes in (b) and (d) and two probably polyploid or G2 stage cells in (c) as judged from differences in cell size and chloroplast numbers per cell. Scale bar = 50 μm. For additional information, see Data S8 and Appendix S2.

### Integrity of ptDNA: search for DNA fragmentation during development

Different from previous claims of massive ptDNA loss already in early leaf development (e.g., Rowan *et al*., 2009), Bendich and co-workers more recently postulated that the organellar DNA may not necessarily be completely degraded during leaf development, but functionally inactivated due to mutations induced by reactive oxygen species (ROS) generated in photosynthesis (Kumar *et al*., 2014, Kumar *et al*., 2015). A major argument for this assumption has been the observation that standard quantitative real-time PCR amplifying short DNA segments of less than 200 bp did not reveal a significant loss of ptDNA during chloroplast development in leaves of light-grown maize seedlings, while long-range PCR generating large DNA segments in the order of 11 kb amplified ptDNA to only 0.1% compared to standard PCR from the same material. The large difference in the yield of amplified ptDNA between the two PCR techniques was suggested to result from unrepaired ROS-induced mutations that increase in number during leaf and organelle development, knowing that mutations like single- and double-strand breaks or pyrimidine dimers can hinder DNA amplification by Taq polymerase or prevent it altogether. It was further argued that this massively damaged ptDNA is degraded to non-functional fragments.

In a subsequent study, Ma and Li (2015) amplified comparable amounts of ptDNA by conventional quantitative real-time PCR and long-range PCR using very similar maize leaf material and biochemical reagents. Under optimized conditions for long-range PCR, they observed no significant difference between the results of conventional and long-range PCR, i.e., obtained no evidence for a destruction of ptDNA in maize leaves. Since Bendich and co-workers had generalized their hypotheses about the degradation of ptDNA and extended them to other species (Kumar *et al*., 2014, cf. also Oldenburg and Bendich, 2015) we assessed quality and integrity of ptDNA during leaf development in several higher plant species by three independent methods other than PCR: by visualizing unfractionated high-molecular mass ptDNA released from gently embedded protoplasts by pulsed-field gel electrophoresis (cf. Swiatek *et al*., 2003), by ultracentrifugation of single- and double-stranded ptDNA in analytical CsCl equilibrium gradients, and by restriction of unfractionated DNA prepared from chloroplasts and gerontoplasts purified by combined differential and isopycnic centrifugation (Figure 7d,e, cf. Schmitt and Herrmann, 1977, Herrmann, 1982). The latter approach largely excludes contributions from non-mesophyll cells. The banding pattern of isolated chloroplasts and gerontoplasts from tobacco and spinach leaves in the isopycnic gradients is shown in Figure S2.

**Figure 7.**
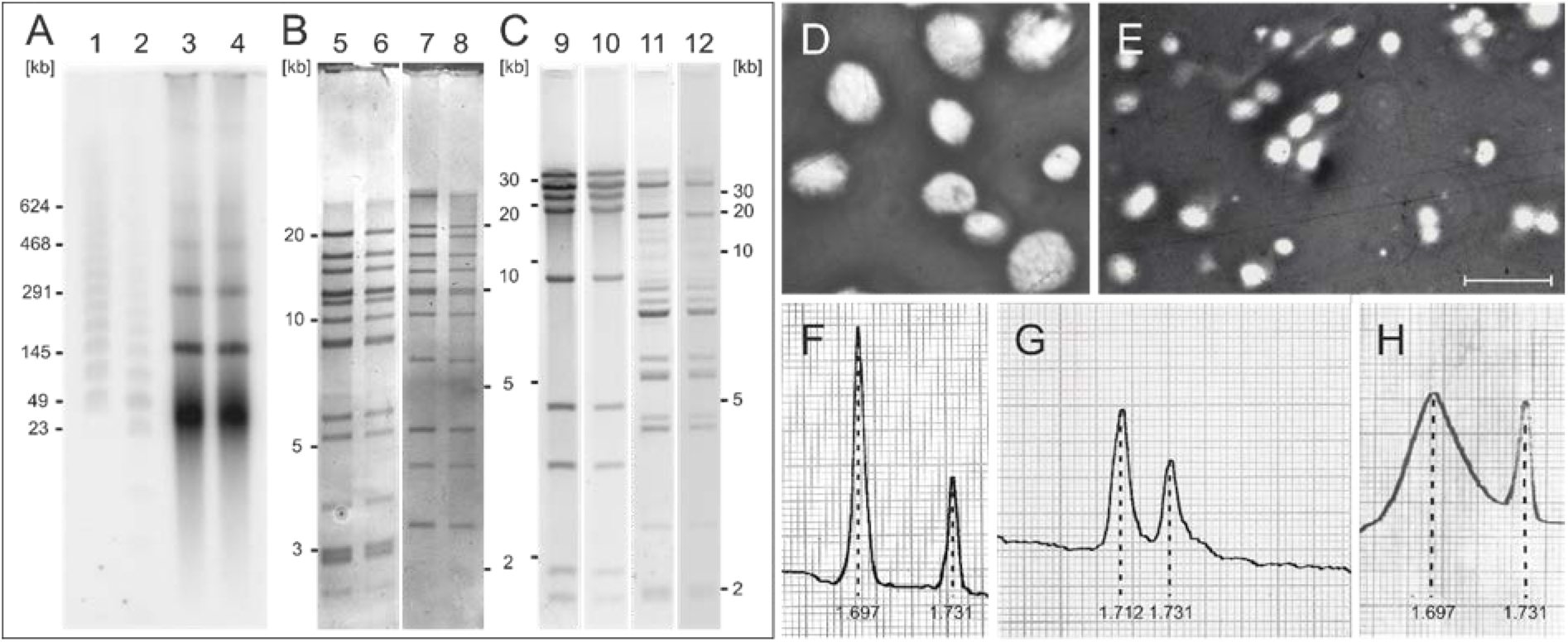
High-molecular mass ptDNA from mature and senescent plastids. (a) Pulsed-field gel electrophoresis of tobacco plastid chromosomes from postmature leaf tissue (lanes 3-4; lanes 1-2 contain size markers). DNA was blotted and hybridized against a radioactively labelled *Nicotiana tabacum* ptDNA library covering the entire plastome (for details, see Material and Methods). The circular ptDNA monomer (156 kbp), dimer (312 kbp), trimer (468 kbp) and tetramer (624 kbp) are clearly visible. The usually seen high molecular weight smear of subgenomic sizes (range 25–45 kbp) presumably represents linear fragments and/or replication intermediates. (b and c) Agarose slab gel electrophoresis of restriction endonuclease-digested DNA isolated from purified mesophyll chloroplasts (lanes 5, 7, 9 and 11) and gerontoplasts (lanes 6, 8, 10 and 12) of mature and yellowish senescent material, respectively, of tobacco (b) and spinach (c). Tobacco: DNA digested with *Xho*I (lanes 5 and 6) or *Pvu*II (lanes 7 and 8), spinach: DNA digested with *Kpn*I (lanes 9 and 10) or *Xma*I (lanes 11 and 12) and separated in 0.5 – 0.7% agarose slab gels; left and right: fragment sizes in kbp (for exact values see Crouse *et al*. 1978, Herrmann *et al*. 1980, Seyer *et al*. 1981). Note the low background in the lanes, especially between the largest fragments, and that sample wells lack significant signals, which is indicative of complete digestion of ptDNA and low nuclear contamination. In each instance, the molecular masses of the detected fragments added up to the size of the circular genome within experimental error. (d and e): Reverse interference contrast micrographs of isolated chloroplasts and gerontoplasts, respectively. The visibility of grana in (d) indicates loss of organelle envelopes due to hypertonic stress in the gradients. The majority of gerontoplasts recovered from gradients were still surrounded by envelopes. Scale bar = 10 μm. Examples of isopycnic sucrose gradients from which plastids were purified are documented in Figure S2. (f - g): Examples of UV scans of banding patterns of native (f) and heat-denatured (g) ptDNA from postmature leaves of spinach centrifuged in neutral analytical CsCl equilibrium gradients at 44,000 rpm for 24 h at 20°C. The gravity field is directed to the right. Peak buoyant densities are indicated. *Micrococcus lysodeikticus* DNA of buoyant density 1.731 g cm^-3^ served as a reference. (h) Molecular weight control, fragmented ptDNA (av. 0.8 x 10^6^ Dalton) of *Antirrhinum majus* (from Herrmann et al. 1975). Note the only moderate increase of the band width in (g) compared to (f).

The tobacco example shown in Figure 7a (lines 3-4) illustrate that comparable amounts of circular monomers and oligomers of plastid chromosomes were present in all leaf samples analyzed. Comparably, restriction analysis of DNA recovered from purified leaf chloroplasts or gerontoplasts with rarely cutting endonucleases verified its high molecular weight and negligible contamination by nuclear DNA. Even the largest fragments in the expected fragment patterns spanning about a quarter or more of the plastid chromosome were present in near-stoichiometric quantities without remarkable background in the gel lanes that would result from broken DNA molecules (Fig. 7b,c, see Discussion). Finally, ptDNA of high molecular weight could also be deduced from narrow banding patterns of native DNA in CsCl sedimentation/diffusion equilibrium gradients, analyzed for seven plant species including maize (e.g., 7f). Those observed with single-strand DNA (7g) excluded increased hidden single-strand breaks, as judged from the DNA size control (h) which expectedly displayed the higher band widths of low molecular mass DNA due to their higher diffusion rates in the sedimentation/diffusion equilibrium gradients. Collectively, our findings verified the presence of a large fraction of essentially intact plastid genomes in all analyzed samples.

## Discussion

### ptDNA is stable during leaf mesophyll development

The present study on the structure, quantity and integrity of ptDNA focused on early stages of mesophyll development and was additionally motivated by the urgent need to critically evaluate and compare methods and techniques that can be used to investigate quantitative aspects of organellar genome dynamics during development (see Introduction). Together with previous work (Li *et al*., 2006, Zoschke *et al*., 2007, Rauwolf *et al*., 2010, Golczyk *et al*., 2014), it provides us with a reasonably complete picture of the fate of the plastome during development from meristematic/post-meristematic to near-necrotic mesophyll in four unrelated vascular plant species and should clarify a number of aspects that have been highly controversial. Using a combination of complementary approaches, we show that substantial amounts of ptDNA are present during all stages of leaf development (Figures 1 and 2, Data S1 - S7). At none of the investigated stages any evidence was obtained for a notable reduction or a significant fragmentation of ptDNA. The high quantum efficiency of DAPI fluorescence and its specificity for double-stranded DNA (Dann *et al*., 1971) permit visualization of organellar DNA uncontaminated by other DNA species directly and unambiguously *in situ.* We have demonstrated that DAPI fluorescence is sensitive enough to detect a single copy of the plastid genome (cf. James and Jope, 1978). The advanced high-resolution epifluorescence microscopy employed in the course of this study allowed us to examine plastids both individually and in the cellular context for structural and quantitative aspects of ptDNA.

The results of our experiments are not compatible with the view that mature chloroplasts contain predominantly highly fragmented and largely non-functional genomes (Oldenburg and Bendich 2015). Pulse-field electrophoresis of total cellular DNA (released upon lysis of immobilized protoplasts) uncovered superhelical molecules, thus verifying the macromolecular integrity of ptDNA. Restriction of ptDNA isolated from gradient-purified chloroplasts or gerontoplasts of late senescent leaf tissue and buoyant density analysis of (heat-denatured) single-stranded ptDNA in analytical CsCl equilibrium gradients (Figure 7) corroborated this finding. The results obtained exclude (i) substantial contamination with nuclear DNA, (ii) the presence of significant amounts of low-molecular mass ptDNA fragments, and (iii) the presence of indigestible high-molecular weight DNA aggregates that remain in the sample wells or in the gel compression zone. The data reveal as well that (iv) the DNA was not damaged by abundant strand breaks and confirmed that organelles from non-mesophyll cells did not contribute substantially to the investigated ptDNA fractions. Thus, our results imply that the plastome copy numbers determined represent predominantly genome-size molecules of mesophyll cells. Moreover plastids in all cells investigated displayed strong and comparable nucleoid fluorescence emission patterns (e.g., Data S2 and S1, panels 220 with more than 30 cells, 221, 217, 218 of *Arabidopsis*, and panels 86, 87 and 114 of sugar beet). This means that a large number of organelles analyzed by us and found to exhibit strong DAPI-DNA signals were from tissue that, according to Rowan *et al*. (2009) and Oldenburg and Bendich (2015), should contain no, very little and/or heavily damaged DNA.

Our findings are also consistent with previous observations, e.g., DNA gel blot data, results of quantitative PCR and ultrastructural work that showed tangled DNA fibrils in plastid nucleoids during all stages of leaf development (Li *et al*., 2006, Zoschke *et al*., 2007, Rauwolf *et al*., 2010, Golczyk *et al*., 2014). The reasons for the conflicting results reported by Bendich and co-workers are not entirely clear yet (Golczyk *et al*., 2014). Given that the various laboratories investigated very similar material, the discrepancies are unlikely to be due to the use of different cultivars or growth conditions. Possible reasons for failed DAPI staining and experimental conditions for long-range PCR of ptDNA have been discussed previously (e.g., Selldén and Leech, 1981, Evans *et al*., 2010, Golczyk *et al*., 2014, Ma and Li, 2015). Further technical issues are discussed in Supplemental Appendix S2.

### Structural aspects of plastome organization during mesophyll development

The predominant mode and common denominator of the spatial organization of ptDNA in mesophyll chloroplasts is a multiple spot pattern of nucleoplasms. This pattern was described from leaf tissue of numerous materials (Herrmann and Kowallik, 1970, Kowallik and Herrmann, 1972, James and Jope, 1978, Coleman, 1979, Kuroiwa *et al*., 1981, Selldén and Leech, 1981, Hashimoto, 1985, Miyamura *et al*., 1986, Fujie *et al*., 1994, Rauwolf *et al*., 2010, Golczyk *et al*., 2014). Our study demonstrates that it lasts from meristematic/postmeristematic to necrotic material, though with notable variation, from single nucleoids in tiny plastids, to multiple clustered, scattered or circular spot patterns. In spite of variation in detail, it also suggests an ordered and recurring sequence of pattern changes during leaf development as well as a remarkable similarity of nucleoid arrangements between quite unrelated species (summarized in Table 1 and schematically in Figure 3). We have found the distinct patterns in all materials studied, though with different frequency and duration, or at varying times during leaf development. Apparently, plastomes of vascular plants share basic architectures and possess the capacity of generating those arrangement modifications, which usually do not reflect distinguishing features between species as occasionally proposed (e.g., Kuroiwa *et al*., 1981, Selldén and Leech, 1981). The respective patterns are transitory and appear to be generated in a relatively flexible way, basically by two processes, (i) on different timing of ptDNA synthesis, nucleoid, organelle and cell division which generally do not occur synchronously, may depend on physiological condition or environment, perhaps also on genotype, and (ii) on the biogenesis and topology of the organelle internal membrane system. Thylakoids and inner envelope membranes, to which DNA is generally attached (Herrmann and Kowallik, 1970, Herrmann and Possingham, 1980), may lead to the distinct nucleoid architectures.

Table 1 summarizes the cytological findings on plastids, nucleoids and ptDNA obtained from post-meristematic to senescent leaf tissue. Organelle numbers, sizes and nucleoid numbers per organelle increased expectedly and approached typical figures seen in mature diploid cells, 28 - 40 (average about 32) organelles, with usually between 18 and >30 discrete and scattered DNA regions per organelle; e.g., Figure 1f,g, Figure 2m, Figure 3g, Data S1 and S2, panels 115ff, 270). Circular nucleoid arrangements, occasionally reported from higher plants, notably from monocots (cf. Selldén and Leech, 1981, Hashimoto, 1985, Miyamura *et al*., 1986, Miyamura *et al*., 1990, Rauwolf *et al*., 2010), seem to be more frequent, quite common, not developmentally restricted (Figure 3d and j), and more diverse than supposed. We have found them usually in knotty closely spaced beads-on-a-string structures in all four species studied, practically at all stages of leaf development (e.g., in meristematic: Fig. 1B, juvenile: Fig. 1C, Fig. 2I, mature: Fig. 1N, senescing mesophyll: see Supplemental Datasets 1 - 4, panels marked with arrow heads and Golczyk *et al*., 2014), and in at least two basic versions. Peripheral circular nucleoid arrangements may be prevailing, occur in all organelles of a cell, particularly conspicuous in maize (Figure 2k,l, Data S4, panels 374 - 380), or were observed in only few organelles. They are transitory; individual nucleoids which are not associated with the peripheral band and increasing in number with progressing development, obviously lead to scattered nucleoid distributions (e.g., Figure 2k,l, Data S4, panels 374-382, but see also Data S2 and S3, panels 270, 271, 326, 327). A different kind of ring-like nucleoid arrangement was now observed in the stroma of plastids of aging and senescent material, apparently linked to the reorganization of the thylakoid system during senescence (Golczyk *et al*., 2014, Fig. 3K; e.g., Fig. 1N, Data S2 and S3, panels 270, 271, 326 - 330, Data S5, panels C and E). Tomographic and ultrastructural analyses indicate that swirled thylakoid membranes and residual membrane patches seen in aging chloroplasts and gerontoplasts are associated with and surround plastoglobuli (Austin *et al*., 2006, Golczyk *et al*., 2014) presumably causing that special nucleoid conformation (Fig. 3K, Golczyk *et al*., 2014). Further details of nucleoid arrangements in plastids and differences among species observed are outlined and documented in Appendix S1.

### Quantitative aspects of ptDNA

Despite the remarkable similarity of quantitative data on ptDNA copy numbers obtained from three different experimental approaches (DAPI-DNA flourescence, real-time qPCR, and previously performed colorimetry with weakly fixed, purified plastids; Rauwolf *et al*., 2010), it should be borne in mind that none of the methods currently available can provide accurate absolute values for ptDNA amounts. None is free of pitfalls, and none of them can address all relevant aspects, including nucleoid number, nucleoid ploidy, number and size variation of plastids in cells, cell size, and nuclear ploidy (cf. Introduction).

ptDNA quantification at the level of individual nucleoids, organelles and cells by measurements of the intensity of the DAPI-DNA fluorescence is generally believed to yield more precise information than other methods (e.g., Miyamura *et al*., 1986, Fujie *et al*., 1994, Golczyk *et al*., 2014). However, even advanced techniques yield only approximate values, due to inaccuracies caused by organelle orientation, focal plane differences, dependence of emission intensities on the nucleoid position within the organelle, differences in self-absorption of fluorescence, extrapolation from tissue sections (Fujie *et al*., 1994), and bleaching of the DAPI-DNA complex with excitation time. By moving the focal plane vertically through the organelle, nucleoid patterns may change substantially as DNA spots become successively visible in different planes and in almost all regions of the stroma (cf. also James and Jope, 1978, Hashimoto, 1985), consistent with early electron microscopic work on matrix-depleted plastids (e.g., Kowallik and Herrmann, 1972). Also, in conventional images obtained at only a single focal level, intense non-focal fluorescent halos obscure details and only focal nucleoids are accessible to analysis. The approach used in our work minimizes these problems, and produces an output equivalent to confocal imaging (Golczyk *et al*., 2014). By combining fast vertical records from different focal planes across an organelle or cell into 2D presentations, it provides superior optical resolution, image sharpness and signal quantification compared to conventional techniques. The main source of inaccuracy observed were (rare) spots of exceedingly high emission signals that are outside the linear range between DNA quantity and emission strength.

qPCR with plastome-specific primer pairs determines ptDNA levels as percentage of the total DNA in a tissue or organ. The values obtained can then be used to calculate plastome copies per cell and, provided that organelle numbers per cell are known, per organelle. Average ptDNA quantities and number of fluorescing spots per organelle provide estimates of average ploidy levels of the nucleoids. Real-time qPCR requires correction for cell types and nuclear ploidy. It usually underestimates ptDNA amounts of mesophyll cells when applied to complex leaf tissues, because non-mesophyll cells such as epidermal cells, cells of the vascular tissue and trichomes, which may amount 40 – 50% of the leaf cell population (cf. Rowan *et al*., 2009, Liere and Börner, 2013), typically harbour fewer and smaller plastids and with significantly fewer ptDNA copies per organelle. On the other hand, qPCR on apical meristems or early post-meristematic leaflets may overestimate ptDNA values, since surrounding post-meristematic tissue (with higher ptDNA quantities per cell) can often not be removed completely. Promiscuous DNA (i.e., nuclear copies of ptDNA sequences) claimed to be a cause of overestimated ptDNA copy numbers (Kumar and Bendich, 2011, Zheng *et al*., 2011), was recently shown to not significantly falsify PCR signals from authentic ptDNA (Udy *et al*., 2012, Golczyk *et al*., 2014).

Four points of general interest emerged from the structural and quantitative findings obtained in this study, and from relevant data in previous work (Li *et al*., 2006, Zoschke *et al*., 2007, Rauwolf *et al*., 2010):

i. The wide range of nucleoid fluorescence emission in individual organelles (e.g., Figure 4, Data S6 and S7) confirms that nucleoids are generally polyploid, with remarkable variation from a single to >20 genome copies (T4 units) per spot. Mean ploidy levels estimated for individual organelles were between 2.3 and 7.5; nucleoid ploidy did not change markedly during leaf development, although slightly lower values were obtained for organelles of meristematic, juvenile and post-mature material (e.g., Figure 1g, Data S1-S3, panels 125, 126, 269, 325). The values of the three approaches used including colorimetric methods (Rauwolf *et al*., 2010) are in excellent agreement and consistent with the analysis of supramolecular membrane-associated DNA complexes isolated from chloroplasts (Herrmann and Possingham, 1980).
ii. Individual plastids harbored 8 - 35 plastome copies in 2 - 6 nucleoids per organelle in meristematic material, and up to about 80 - 130 plastome copies in 20 - >30 nucleoids in mature chloroplasts. The data were remarkably similar for the four species studied. Copy numbers, nucleoid numbers and organelle size were usually correlated.
iii. Our quantifications support a continuous rise of ptDNA levels per organelle and cell during development from post-meristematic/juvenile to near-mature mesophyll tissue that correlates with proplastid-to-chloroplast differentiation (Figure S1). The bulk of ptDNA was synthesized relatively early, and maximal levels were usually reached at premature stages (i.e., before a cell-type specific chloroplast number was established, before organelles assumed their final volume, and before cells were fully elongated and leaves fully expanded). Cellular ptDNA levels increased from about 75 - 120 plastid genome copies in early post-meristematic tissue for all four species studied to maximal levels of 2,750 to 3,200 copies per diploid cell in premature sugar beet mesophyll, 2,620 to 3,080 in *Arabidopsis*, 2,320 to 2,800 in tobacco, and 2,550 to 3,150 in maize (Table 1; cf. also Aguettaz *et al*., 1987, Evans *et al*., 2010, Udy *et al*., 2012, Ma and Li, 2015). The developmental changes determined correspond to an approximately 9.3-fold increase in ptDNA per organelle (and 24-fold per cell) from proplastids to chloroplasts for diploid sugar beet mesophyll cells, which is primarily due to plastid growth and multiplication (see also Rauwolf *et al*., 2010). This number (and the similar numbers for the other three species) are well in line with the 7.5-fold increase in ptDNA per organelle (34-fold per leaf cell) reported for hexaploid wheat (Miyamura *et al*., 1986).
iv. Several observations made in the course of our study suggest that the regulation of cellular genome-plastome homoeostasis during leaf development is more complex than previous work suggested. In general, nuclear ploidy and cellular organelle numbers are correlated in that chloroplast number almost doubles upon tetraploidization (e.g., Butterfass, 1979), as also confirmed in this study. Nuclear ploidy changes do not substantially alter cellular genome-to-plastome ratios, since chloroplast size and DAPI patterns in di- and tetraploid cells are virtually indistinguishable (cf. Data S1-S4). Somatic endopolyploidization is usually negligible in juvenile tissue, but increases substantially with leaf age, and needs to be corrected for in ptDNA quantification. However, it is important to note that the mechanisms that maintain constant genome ratios do not operate at all developmental stages. For example, the influence of nuclear ploidy on plastid number and size in sugar beet was evident in mature mesophyll, but barely detectable in juvenile leaf tissue (Rauwolf *et al*., 2010). Similarly, variable chloroplast numbers that do not strictly correlate with the endopolyploidy levels were reported for *Arabidopsis* (Pyke and Leech, 1991, Barow, 2006, Zoschke *et al*., 2007). Also, the intriguing giant cells observed in this study in *Arabidopsis*, tobacco and sugar beet harbor several hundred chloroplasts, but may not exhibit an equivalent increase in nuclear volume, as it is generally seen with polyploidization (Data S5). Whether this reflects unknown regulatory circuits that alter genome-plastome ratios or, alternatively, is due to extensive endopolyploidization without much change in nuclear volume, remains to be investigated.

Taken together, the data described here provides a general picture of the structural organization of plastomes during leaf mesophyll development. They verify the overall stability of the plastid genome and indicate that plants adjust plastome-genome homoeostasis flexibly during development and adaptation and suggest that the adjustment of cellular genome ratios is substantially more complex than presently assumed. Exploring the underlying mechanisms represents an attractive topic for future research.

## Experimental procedures

### Plant material

The plant material used, greenhouse growth of plants, and collection and treatment of defined tissue samples were essentially as described for *Arabidopsis thaliana,* tobacco and maize in Golczyk *et al*. (2014) and for spinach (*Spinacia oleracea*) and sugar beet in Herrmann *et al*. (1975) and Rauwolf *et al*. (2010). The diploid sugar beet cultivar “Felicita” was obtained from KWS Saat AG (Einbeck, Germany). Leaflets, leaves and explants were classified according to developmental stages. “Stage 1” represents meristematic and early post-meristematic explants from the innermost shoot apex (≤1 mm in *Arabidopsis*, ≤2.0 mm in tobacco and maize, ≤2.5 mm in *Beta vulgaris*). “Stage 2” comprises the first leaflets of 1.5 - 3 mm length in *Arabidopsis*, 2 - 10 mm in tobacco, 4 - 16 mm in *Beta vulgaris*, and 2 - 4 mm from the leaf base in maize. “Stage 3” represents leaflets of 2.5 - 4 mm from *Arabidopsis*, 1 - 2.3 cm from tobacco, 1.5 - 3.5 cm from *Beta vulgaris*, and approximately 1.5 cm above the vegetation point in maize. “Stage 4” leaflets are 4 - 8 mm long in *Arabidopsis,* 2 - 5 cm in tobacco, and 3 - 7 cm in *Beta vulgaris.* “Stage 5” represents juvenile leaves of ≥8 mm in *Arabidopsis,* 4 - 9 cm in tobacco, 5.5 - 9.5 cm in *Beta vulgaris*. “Stages 6 - 8” include premature (e.g., 8 - >12 cm in *Beta vulgaris*), mature and early aging leaves (equivalent to stages II, III and IV in Golczyk *et al*., 2014). Shoot apices were excised with scalpel and forceps under a dissecting microscope. Lamina sectors of green young and nearly mature maize leaves were taken as “stage 4” and “stage 5” samples, respectively. Laminas of sugar beet leaflets of “stage 2” were curled, “stage 3” samples contained leaflets with curled as well as expanded laminas (for images, see Rauwolf *et al*., 2010).

### Microscopy and DNA quantification of nucleoids

DAPI (4’,6-diamidino-2-phenylindole) staining and fluorescence microscopy were conducted as described in Golczyk *et al*. (2014). The staining specificity of the trypanocide fluorochrome was verified as reported previously Rauwolf *et al*. (2010) and Golczyk *et al*. (2014). A T4 phage suspension was purchased from the American Type Culture Collection (ATTC), Manassas, VA, USA [T4 bacteriophage (ATCC® 11303B4™)]. For the ptDNA fluorescence densitometry, a small aliquot of phage suspension was dried on a microscope slide, and tissue explants were mounted close-by on the same slide, gently squashed in a drop of PBS buffer (137 mM NaCl, 2.7 mM KCl, 10 mM Na_2_HPO_4_, 1.8 mM KH_2_PO_4_, pH 7.4), frozen in liquid nitrogen, and air dried after removal of the cover slip. The relatively constant phage fluorescence emission, ranging from 0.85 - 1.1 (av. 1.0 ± 0.1) arbitrary units, can be taken as ploidy unit and used for normalization of nucleoid emission intensities, because coding potential (Freifelder, 1970) and GC content resemble that of plastomes. DNA of individual nucleoids in magnified plastids was quantified by microphotometry, through integration of high-resolution records taken rapidly at different focal planes along the z-axis of the organelle. Images were acquired with a Nikon Eclipse Ni-U epifluorescence microscope equipped with a cooled monochrome camera DS-Qi1, as described previously (Rauwolf *et al*., 2010, Golczyk *et al*., 2014), and the ImageJ software (Fiji package, https://imagej.net/Fiji/Downloads) was used for image processing. After downloading the original camera recorded image files (left panels in Figure 4 and Data S6), fluorescing nucleoids were delimited and corrected for background using the Wand Tool and Tolerance Adjustment Regulation (central and right panels, respectively, in Figure 4, right panels in Data S6). Their pixel area and overall pixel density (= integrated density) were calculated using the function “Measure run” from the “Analyze” menu. Data were also analysed visually with a magnifier and a graded series of *in silico* quantified fluorescence spots of increasing emission intensity. Intensities of individual nucleoids were expressed as equal or multiples of that of phage heads. Occasionally, the weakest organelle spots displayed fluorescence emissions up to 25% lower than phage particles. This effect, presumably in part due to different degrees of DNA compaction, was disregarded. Figures of a given picture series are directly comparable, since images of DAPI stained suspensions of T4 phage particles and those employed for cells or tissues were recorded under identical conditions. Comparisons between species are also feasible since base composition and base heterogeneity of plastomes are very similar.

### Protoplast preparation

Protoplasts from mature leaf tissue were prepared according to protocols previously described for sugar beet and tobacco (Huang *et al*., 2002), *Arabidopsis* (Wu *et al*., 2009) and maize (Edwards *et al*., 1979). Polyploid cells were estimated on the basis of cell sizes and chloroplast numbers. The ratio of di- and tetraploid protoplasts in sugar beet was deduced from about 800 individual cells (Fig. 6 and Supplemental Dataset 8; Butterfass, 1979). Gentle agitation of tissue explants during enzymatic protoplast release prevented artificial cell fusions via cell-connecting plasmodesmata (Hecht’s threads) during preparation. No binucleate protoplasts which would result from cell fusion were detected.

### Pulsed-Field Electrophoresis (PFEG)

To avoid possible ptDNA degradation during chloroplast isolation (cf. Discussion in Golczyk *et al*., 2014), full-length plastid genomes were prepared from agarose-embedded protoplasts of mature tobacco leaves. Protoplast suspensions (8 x 10^6^ cells per ml) were gently mixed with three parts of 1.1% low-melting-point agarose. The embedded cells were then lysed and DNA was separated using a CHEF Mapper® XA System (BioRad, Munich, Germany) essentially as previously described (Swiatek *et al*., 2003). The DNA was then blotted by alkaline transfer onto a nitrocellulose membrane and hybridized to a radiolabelled *Sal*I restriction fragment library covering the entire plastid genome of *Nicotiana tabacum* in 11 ptDNA fragments inserted into vector pBR322 (Medgyesy *et al*., 1985). Radiolabelled signals were detected with a phosphoimager screen and acquired with a Typhoon^TM^ TRIO+ scanner (GE Healthcare, Buckinghamshire, UK).

### Quantitative real-time PCR, purification of chloroplasts and gerontoplasts, and analytical ultracentrifugation of DNA

DNA was isolated according to Doyle and Doyle (1987). Quantitative PCR was performed essentially as reported in Zoschke *et al*. (2007). Primer sequences are summarized in Table S1. All amplified regions are unique and occur only as single copy per plastid genome. Homogenization of leaf tissue, treatment of homogenates, purification of chloroplasts and gerontoplasts by differential and isopycnic centrifugation techniques, isolation and restriction of unfractionated high-molecular mass ptDNA, and slab gel electrophoresis of restriction digests were performed as described in Schmitt and Herrmann (1977) and Herrmann (1982). Continuous linear 20 - 60% sucrose gradients were used. Analytical ultracentrifugation of DNA in neutral CsCl solutions was performed as described in Herrmann *et al*. (1975).

## Acknowledgements

The authors thank Liliya Yaneva-Roder for excellent technical assistance. We are grateful to Dr. Loock and Mr. Hauer (KWS Saat AG, Einbeck, Germany) for providing the sugar beet line, and to the MPI-MP Green Team for plant cultivation. This work was supported by the Max Planck Society to R.B. and S.G. The ptDNA DAPI fluorescent patterns were analyzed with microscopy equipment funded by Polish National Science Center - Grant 2015/19/B/NZ2/01692 to H.G.

## Supporting Information

## Supplemental Tables

**Table S1.**
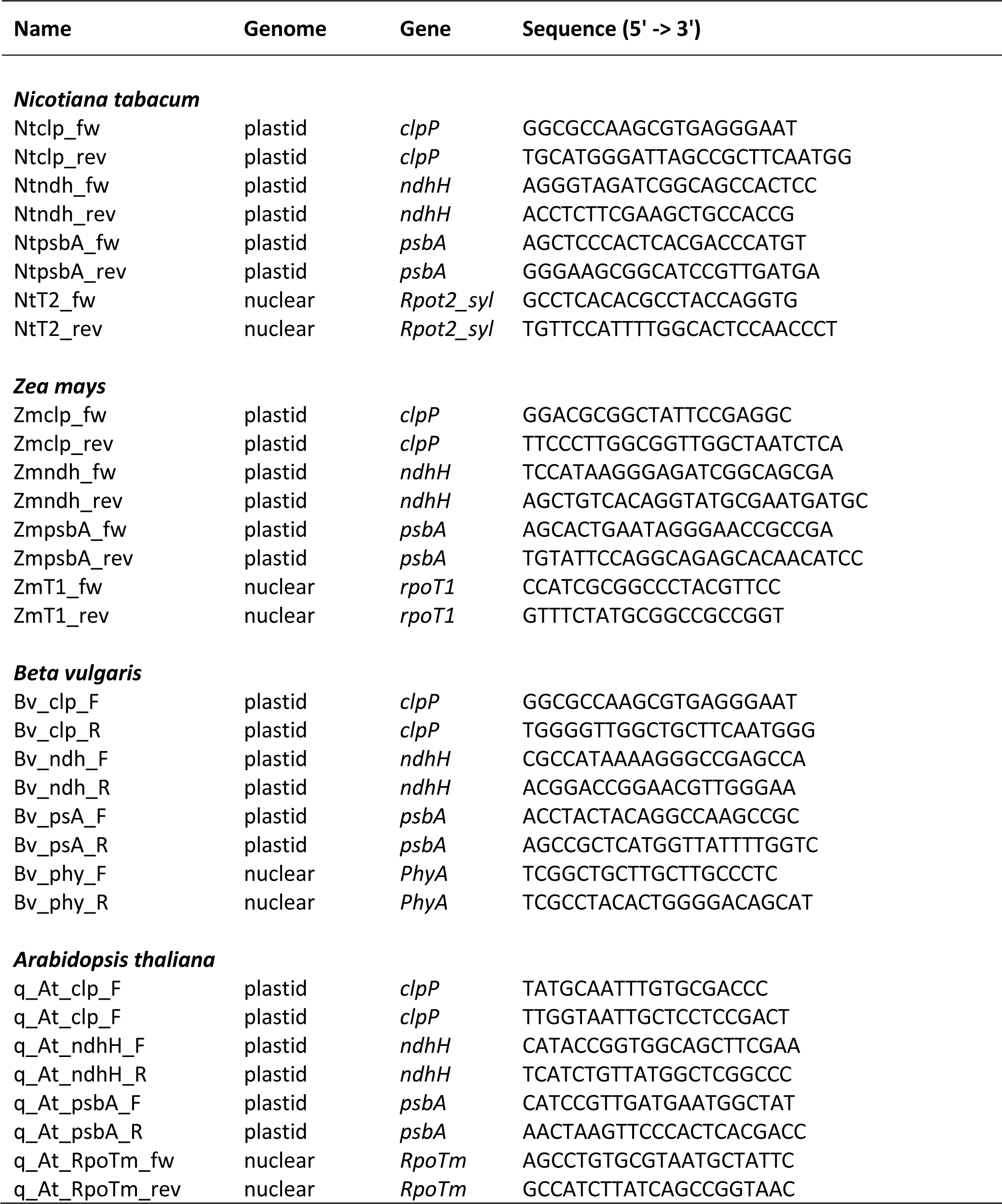
Oligonucleotides used for real-time qPCR.

## Supporting Figures and Legends

**Figure S1.**
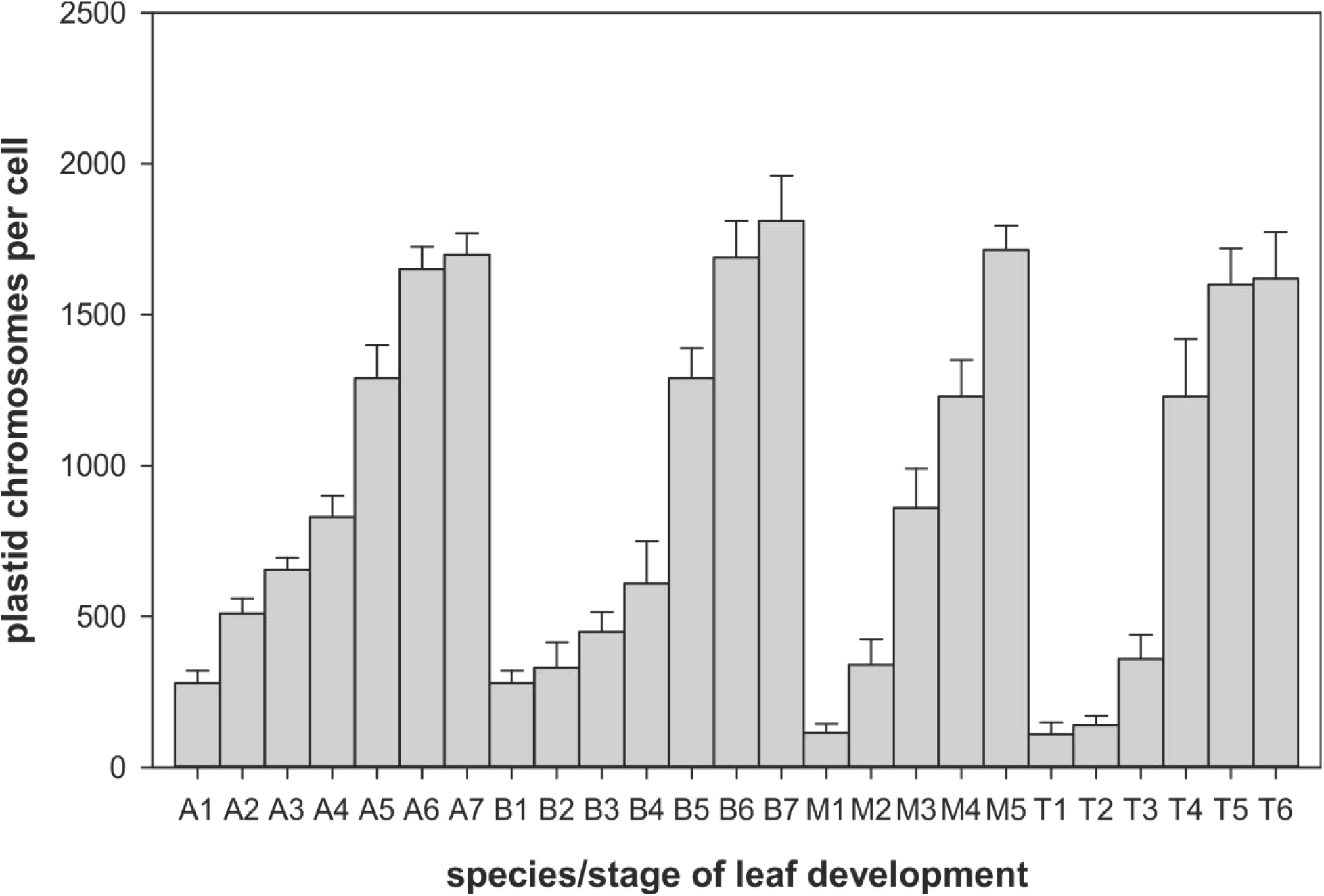
Plastome copy numbers per cell during early leaf development determined by real-time qPCR with total cellular DNA as template against nuclear reference genes [see Material and Methods and Golczyk et al. (2014) for comparable and complementary data]. The graphs show means and standard deviations from four measurements per leaf sample (three replicates per gene). (Stages A1-A6) *Arabidopsis*, (B1-B7) sugar beet, (M1-M5) *Zea mays*, and (T1-T6) tobacco. The data agree well with spectrofluorimetric estimates after correction for cell type and ploidy. Some variation is presumably caused by differences in tissue sampling (see also Discussion). For description of developmental stages, see Material and Methods.

**Figure S2:**
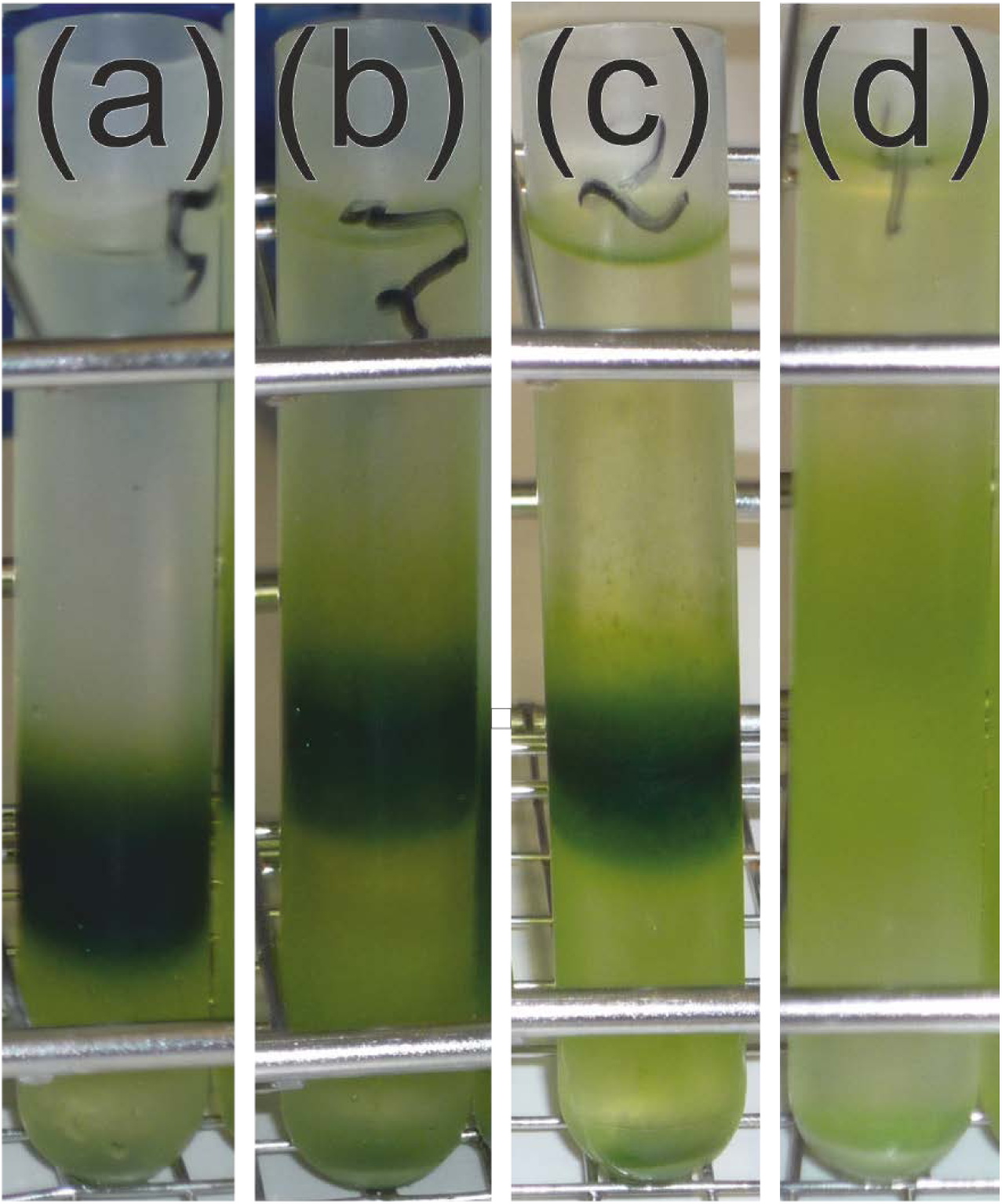
Chloroplast and gerontoplast fractions of tobacco and spinach purified by centrifugation at 30,000 *g* for 30 minutes at 0°C in isopycnic continuous 20 – 60% sucrose gradients. Initial organelle fractions were obtained from filtered homogenates by short-time differential centrifugation as described in Schmitt and Herrmann (1977, p. 182) and Herrmann (1982). Gradients: (a) Chloroplasts of mature leaves and (b) chloroplast/gerontoplast intermediates from aged leaves of tobacco, (c) chloroplasts of mature leaves and (d) gerontoplasts of senescent greenish-yellow spinach leaves. Note that the banding position of plastids in the gradients depends on both, species and leaf developmental stage. Chloroplasts from tobacco band at higher buoyant densities (lower gradient positions) than those of spinach. Chloroplasts of mature tissue, in turn, were found at higher buoyant densities than those of aging leaves, likely indicating a relatively higher lipid content of the latter. Gerontoplasts of yellow leaf sectors were heterogeneous and banded in a wide density range, i.e., over nearly 50% of the gradient volume (see also Figure 7).

## Supporting Data

**Data S1 - S5** illustrate the enormous structural and quantitative variability of plastids and their DNA predominantly during early leaf development. The figures complement corresponding Datasets in Golczyk et al. (2014) dealing with ptDNA from mature to near-necrotic mesophyll.

**Data S1.** DAPI-stained cells in primordial tissue at and around shoot apices and their development to photosynthetic mesophyll cells of early developing leaves (up to about 10 cm) of *Beta vulgaris* (sugar beet), grouped into 4 developmental classes (panels 1 - 128). Note that panels 86 - 88 and 114 display cell clusters in which *all* chloroplasts are well stained. Arrowheads mark examples of circular nucleoid arrangements. Scale bars = 5 μm, in panel 87 also for panels 84 and 86

**Data S2.** DAPI-stained cells from primordial tissue at and around shoot apices and their development into photosynthetic mesophyll cells of early developing leaflets (up to about 1 cm) of *Arabidopsis thaliana,* grouped into 4 developmental classes (panels 129 - 271). Panels 217, 218, 220, and 221 display cell clusters in which nucleoids of all chloroplasts are well stained. Arrowheads mark examples of circular nucleoid arrangements. As judged from nuclear size, cell size and chloroplast numbers, panel 271 shows a polyploid mesophyll cell from postmature leaves with circular nucleoid arrangements in plastids (see also panel 270 and Golczyk et al., 2014). Scale bars = 5 μm, in panel 222 also for panels 217, 218, 220 and 221

**Data S3.** DAPI-stained cells from primordial tissue at and around vegetation points and their development into photosynthetic mesophyll cells of early developing leaves (up to about 9 cm) of *Nicotiana tabacum* (tobacco), grouped into 5 developmental classes (panels 272 – 330). Arrowheads mark examples of ring-like nucleoid arrangements. Note that circular nucleoid arrangements are frequent in panels 327 - 330. Scale bar = 5 μm, in panel 325: 10 μm

**Data S4.** DAPI-stained mesophyll cells of yellow and faintly green primordial tissue at and around leaf vegetation points of early developing, green and dark green lamina samples of *Zea mays* (maize), arranged in 4 developmental groups (panels 331 - 384). Note that circular nucleoid arrangements predominate in stage 4. Bar = 5 μm, in panels 378 - 384: 10 μm

**Data S5.** Giant mesophyll cells with 100 or more chloroplasts in premature to early aging leaves of *Beta vulgaris* (a), tobacco (b-e) and *Arabidopsis* (f). For further *Arabidopsis* cells, see Data S2 online, panel 271, and Golczyk et al. (2014). Note the relatively small nuclei in cells shown in panels (a), (b) and (d), the typical nucleoid pattern in the magnified organelle sector shown in panel (c), and ring-like nucleoid arrangements in (e) and (f) (see also text). Scale bars = 10 μm in (c), (e) and (f), 20 μm in (a) and (d), and 30 μm in (b)

**Data S6.** Quantitative microfluorimetry of nucleoids of randomly selected individual DAPI stained mesophyll chloroplasts from expanding, premature and mature leaves of sugar beet (a-f), tobacco (g-k), *Arabidopsis* (l-s) and maize (t-w), see also Figure 4. The high-resolution microphotographs illustrate the considerable fluorescence variation between DNA spots (left panels). Nucleoid ploidy profiles were normalized either to that of DAPI-stained T4 phage particles (see Figure 4 and tobacco data in this Supplement Dataset for fluorescence in T4 phage suspensions) and/or related to the intensity of the lowest detectable signals in organelles which corresponded to that of T4 particles and served as an additional organelle-internal haploid standard. The organelles shown were selected from different experimental series and may differ somewhat in their magnification; they were analyzed with the respective T4 standard. Three cycles of nucleoid measurements were carried out for each organelle. The results were also compared with corresponding values gathered visually by three independent investigators with the aid of a graded series of nucleoids of determined ploidy. Note that spectrometrically and visually determined values agree well. Of about 55 individual chloroplasts investigated in this experiment, about 30% differed between 7 and 12%, about 50% between 13 and 20%, the remaining cases up to 30%. Major differences resulted from intensely fluorescing spots, as expected (see Discussion). For details see Material and Methods and Main Text.

**Data S7.** Examples of DAPI fluorescence variation among nucleoids in mesophyll chloroplasts. The high-resolution microphotographs from about 100 organelles illustrate the enormous heterogeneity of nucleoid fluorescence emission in chloroplasts of *Nicotiana tabacum* (tobacco), *Zea mays* (maize), *Beta vulgaris* (sugar beet) and *Arabidopsis thaliana*. Organelles with diameters ranging from 1.5 - 7.0 μm were randomly selected from cells of young to postmature leaves. Scale bars = 2 μm, for sugar beet: 1.5 μm

**Data S8.** Examples of purified mesophyll protoplasts from premature and mature leaves of *Arabidopsis thaliana* (a– d), sugar beet (e – h) and tobacco (i – l). Scale bars = 50 μm [(a) as for (b); (g) and (h) as for (f), (i) and (k) as for (l)]. (b, e, h, i and l) show protoplasts from premature, (a, c, d, f, g, j and k) from mature mesophyll. Note examples of rarely present contaminating non-photosynthetic leaf cells in (b) and (f) (arrows). Arrowheads in (a, d, f, g and j) mark cells that are likely polyploid, as judged from larger sizes and higher chloroplast numbers. Samples prepared from premature material display relatively homogeneous cell populations, preparations of mature and postmature material exhibit higher heterogeneity of cell sizes.

## Appendix S1 Nucleoid patterns in plastids during early leaf development

The following data complement information given in the chapters Results and Material and Methods of the Main Text.

*Stage 1*: Cells of 10 - 15 µm in diameter in the 1 - 2.5 mm pale or yellowish region at or around the shoot apex of *Beta* contained 5 - 9 (occasionally up to 12) small plastids (approx. 1 μm in diameter) with low numbers (generally 2 - 5) of nucleoids; organelles with only single nucleoplasms were observed exclusively in the proplastids or leucoplasts of the innermost apical region (cf. also Herrmann and Kowallik, 1970). After division nucleoids assume clustered or scattered positions, or are arranged peripherally in ring-shaped (spot) patterns. Comparable plastid numbers and nucleoid patterns were found in 0.5 - 1 mm meristematic/postmeristematic leaflet explants of *Arabidopsis*, usually in cells of the corresponding yellow or faintly green leaf base of maize, and with somewhat higher numbers in tobacco (6 - 18; Figure 3a-d, Figure 1a, b, h and i; Figure 2a, g and h, Data S1-S4, panels 1-52, 129-162, 272-293, 331-348; see also Herrmann and Kowallik, 1970; Kuroiwa et al., 1981; Hashimoto, 1985; Miyamura et al., 1990). The proportion of plastids with four or more nucleoids was significantly higher in developmentally somewhat advanced tissue, in about 1.5 mm leaflets of *Arabidopsis* and 2 - 5 mm leaf foliage explants of tobacco and *Beta*. Cell sizes, cellular plastid and nucleoid numbers per organelle, but barely organelle sizes, had increased moderately.

*Stages 2 - 3*: With further leaflet development, i.e., to 4 - 16 mm in length of sugar beet, up to about 1.5 cm in tobacco, 1.5 - 3 mm of *Arabidopsis*, and in the (faintly green) leaf base of maize, cells had increased to ≤20 μm. During all early development, in juvenile tissue they appeared more or less round-shaped, leaf laminas were yellow-greenish and still curled in sugar beet, less curled and green in tobacco, and expanded and green in *Arabidopsis*. Corresponding regions close to the leaf base in maize were faintly green. At these stages, remarkable heterogeneity in intracellular organelle arrangement, cell and organelle sizes, nucleoid numbers and arrangement, and nucleoid division became apparent in all species, which presumably reflects the intense leaf growth phase and/or an adaptive flexibility of the system. Dispersed and circular spot patterns could be observed, the latter occasionally with high frequency (Figures 1b and c, 3d-f, 2i, Data S1-S4, e.g., panels 21, 68, 71, 85-87, 89, 166, 197, 212, 220, 227, 268, 270, 271, 299, 302, 317, 358, 362. 363, 365, 370, see Discussion). In sugar beet and maize cells usually contained 8 - 16 (occasionally 12 to about 20) plastids with a limited number (in the range of 6 to 14) of generally scattered nucleoids (Figure 3e, Figure 1c-e, Figure 2j, e.g. Data S1 and S4, panels 53ff and 349ff for sugar beet and maize, respectively; see also Golczyk et al., 2014). Smaller cells with fewer, smaller organelles (2 - 3 μm in diameter) and fewer DNA spots per organelle were still quite frequent. However, at that stage plastids in *Arabdiopsis* (Data S2, panels 183-216) and tobacco (Data S3, panels 301-319) could house relatively high numbers of densely packed, often barely resolvable (e.g., Figure 3f, Figure 1l and m, Figure 2e and f, Data S2 and S3, e.g., panels 181ff, 301ff; Figure 3f) DNA containing areas indicating intense DNA synthesis and nucleoid division without much organelle division. When fewer nucleoids per organelle were present, their fluorescence emission was often brighter (e.g., Figure 3e, g, Figure 1f, Fig 2j and m). At these stages, plastid clustering at cell surfaces began to replace the initially more or less scattered organelle arrangements. Collectively, these findings indicate that ptDNA synthesis may occur with or without notable concomitant organelle or nucleoid division, and that the rates of ptDNA synthesis may more or less be related to or precede the generation of an elaborate internal membrane system (e.g., Data S3, panels 310ff, cf. also Selldén and Leech, 1981).

*Stages 3 - 4*: In 2.5 - 4 mm leaflets of *Arabidopsis,* and 1.5 - 3.5 cm leaflets of sugar beet and tobacco, cells (≤30 µm) usually harbor tightly packed 10 - 22 chloroplasts of 2 - 5 µm diameter with numerous barely resolvable scattered nucleoids (15 -> 20; e.g. Figure 3g, Figure 2f, Data S1 and S2, panels 107ff, 251ff, see also Golczyk et al., 2014). Lower figures (8 - 15), generally with bright fluorescence emission, were observed as well, notably in sugar beet leaflets still with curled lamina, and maize (e.g., Figure 1f). Heterogeneous cell populations observed including relatively small, often still round-shaped cells with varying chloroplast numbers and sizes, smaller chloroplasts in pairs, and conspicuous variation of nucleoid numbers and sizes in and between organelles, again probably reflect developmentally active tissue.

*Stages 4 - 5:* During further leaf development, in pre-mature leaves with lamina extensions up to about 9.5 cm in sugar beet and tobacco, and 4 - ≥8 mm in *Arabidopsis,* cells increase, often by elongation, and may house 14 - 25 organelles that may or may not enlarge simultaneously (e.g., Figure 1f and m, Figure 2e and f). At this stage, cells had reached only about three quarters of their volume (sizes of about 40 - 50 µm) and not established the typical average organelle numbers of mature diploid leaves, with means found in the range of 25 - 35, occasionally ≥45, chloroplasts of 5 - 7.5 µm in diameter and 14 - >30 usually dispersed nucleoids (average around 23); circular nucleoid arrangements were noted as well, especially in *Arabidopsis*, tobacco and maize [Figure 3i-j, Figures 1n, 2k and l, Data S1-S4, e.g., panels 270, 271, 328, 329, 374-380; in “giant” cells: Data S5, panels (c) and (e)]. However, nucleoid arrangements appeared to be more or less terminal and maximal cellular ptDNA amounts were attained already at premature stages, i.e., before a final, relatively stable number of chloroplasts per cell was established and organelles and cells were still enlarging (see also below). The observations are consistent with previous findings that gross DNA replication in plastids appeared to cease before cell proliferation is complete and that ptDNA contents per organelle (and cell) increase generally until that stage, but not notably later. During organelle expansion, chloroplasts shift towards the cell surface.

After cessation of organelle division cells and chloroplasts in mature and post-mature leaves may expand further with continuing leaf ageing. This can happen without significant increase of DNA content (Figure 3h), for distances between individual DNA regions increase, while their fluorescence intensities and numbers remain virtually unchanged. On the other hand, nucleoids may also continue to divide without substantial preceding DNA synthesis reaching numbers in the order of 40 or more spots per plastid, spread throughout the organelle interior, as conceived from significantly lower nucleoid fluorescence (Figure 3i; e.g., Figure 1g, Data S1-S3, panels 125, 126, 269, 325; Golczyk et al. 2014, cf. also Selldén and Leech, 1981; Miyamura et al., 1986). Finally, with organelle division and/or enlargement, ptDNA synthesis may continue to some extent, predominantly due to endopolyploidization (but see Data S5 and Discussion). This process increases in mature leaf tissue and may even prevail depending on plant material (Figure 6a and b, Data S8, Butterfass, 1979). Polyploidization is negligible in juvenile material. Occasionally observed almost doubled plastid numbers in juvenile cells probably reflect G2 cell cycle stages (e.g., Data S1, panel 82, see Butterfass, 1979).

Circular arrangements of nucleoids were first described from plastids of chromophytic algae (Bisalputra and Burton, 1969; Gibbs et al., 1974) in which the organelle DNA is associated with girdle lamellae, a specific thylakoid type that lies inside the organelle rim and forms a loop of nucleoids attached adjacent to one another around the organelle periphery. Plastids of vascular plants obviously possess the capacity of this peculiar arrangement although they seemingly lack that specific membrane type. In fact, ring-like nucleoid organization, occasionally reported from higher plant plastids, notably from monocots (cf. Selldén and Leech, 1981; Hashimoto, 1985; Miyamura et al., 1986; Rauwolf et al., 2010), appears to be more common and more complex than assumed currently. We have found it during leaf development in all four species studied, with remarkable variability, in at least two versions, and, different from the algal case, of transitory nature (Figure 3j, e.g., Figure 2k and l, Data S4, panels 370 - 384, cf. Hashimoto, 1985; see also Main Text). The ring-like arrangements in higher plant plastids resemble the knotty structures seen in algae; occasionally they appear as more or less continuous bands that usually resolve into closely spaced spots at higher magnification, presumably reflecting envelope- or thylakoid-attached individual nucleoids (cf. Herrmann and Kowallik, 1970; Herrmann and Possingham, 1980). Elongated narrow bands represent side views suggesting that the ring conformation lies almost perfectly in one plane around the organelle periphery. Ring circumferences and implicitly nucleoid numbers (and DNA quantities) per ring increase with organelle expansion (size/quantity rule). We observed a seemingly different kind of circular nucleoid arrangement in plastids of aging and senescent leaves in the organelle stroma around plastoglobuli that is probably correlated with the reorganization of the thylakoid system during senescence (Golczyk et al., 2014, Figure 3k; e.g., Figure 1n, Data S2 and S3, panels 270, 271, 326 - 330, Data S5, panels (c) and (e)). The prerequisites for these peculiar nucleoid patterns are not known.

## Appendix S2 Critical aspects of methodology

The analysis of DNA from chloroplasts is complicated by (i) the difficulty to avoid contamination by nucDNA during organelle isolation, and (ii) difficulties with reliably determining the type-purity of ptDNA for a large number of plant species. Assessment of findings and conclusions drawn must, therefore, critically consider the quality of the subcellular fractions used, which depends on isolation buffers and purification conditions. Subcellular fractions have to be clearly defined, non-physiological conditions have to be avoided, and information on controls should be given. The latter is particularly important for the validation of negative results.

### Purity of chloroplast fractions

In several studies, Bendich and co-workers applied two kinds of media for tissue homogenization, the so-called high-salt medium (containing 1.25 M NaCl) and an osmotically balanced, sorbitol-based medium with or without PVP. “High-salt” treatment is supposed to remove contaminating nuclear DNA from the resulting chlorophyll-containing subcellular fraction (Oldenburg et al., 2006; Shaver et al., 2006, p. 75 and 80; Rowan et al., 2007). However, “high salt” can destroy organelle envelopes and yields thylakoid fragments largely depleted of stroma, but no intact chloroplasts (seen in Rowan et al., 2007, p. 11; or Rowan et al., 2009, p. 15). Such fractions are generally contaminated by significant amounts of nucDNA, since exposed thylakoid systems can readily entrap remnants of nuclear chromatin during preparation, which subsequently cannot be removed completely by washing. This problem can be revealed by comparison with conventionally prepared fractions from materials with ptDNA and nucDNA of sufficiently different GC contents to be separable in CsCl equilibrium gradients. Reduction of contaminating nucDNA to ≤5% is possible, but requires special precautions in the preparation of organelles (Herrmann et al., 1975; Schmitt and Herrmann, 1977; Herrmann, 1982). Unclear remains why high salt treated subcellular fractions were resuspended in the osmotically balanced medium (Rowan et al., 2007; Rowan et al., 2009).

### Type-purity of ptDNA

Checking type-purity by centrifugation of isolated native ptDNA in CsCl gradients is not applicable to the majority of vascular plant species studied because their ptDNA and nucDNA possess similar base composition and, hence, similar buoyant density. For these species, the difference in reassociation velocities in denatured DNA mixtures (due to different genomic complexity of the two DNA species) and accompanying buoyant density shifts of single- and double-stranded DNA in CsCl equilibrium gradients has been widely used (e.g., Lamppa and Bendich, 1979; Scott and Possingham, 1983, p. 1757). This *a priori* appealing approach operates with mixtures of the T4 phage/salmon sperm DNA pair that has been vicariously used for ptDNA and nuclear DNA, respectively, as a control model (Herrmann et al., 1974). However, this method cannot be applied to assess cross-contamination of ptDNA and nucDNA, because both DNA species cross-react during reassociation due to DNA promiscuity, thus preventing their stoichiometric segregation (Herrmann et al., 1974).

### Integrity of isolated chloroplasts

The use of suspensions of envelope-bounded chloroplasts prepared in osmotically balanced sorbitol-based media bears the risk of artefact, especially, if fractions are prepared with relatively high gravity fields and/or prolonged centrifugation times. The preparations may be contaminated by various kinds of subcellular particles, including some that possess hydrolytic activity, which may adversely affect the integrity of chloroplasts. The objection of artificial leakiness of envelopes is also valid for envelope-bounded plastids prepared in isotonic sorbitol-based media containing PVP. The relative lipophily and the probable detrimental effect of PVP are evident from its chemical formula. As mentioned previously (Golczyk et al., 2014), chloroplasts prepared in the presence of PVP may appear morphologically intact, but may not be so physiologically, in that their envelopes may be permeable to various kinds of compounds including endogenous nucleases. Whether the medium contains EDTA or Mg^2+^ is not relevant here, because not all potentially interfering hydrolases require the bivalent cation as a co-factor. A straightforward control experiment – isolation of DNA from DNase-treated unbroken chloroplasts that were or were not exposed to PVP – could illustrate its effects on organelle envelopes.

### Protoplast integrity

The integrity of protoplasts should be checked. Globular shapes and smooth outlines are characteristic of viable turgescent protoplasts capable of responding osmotically. The DNA of injured or damaged cells is potentially prone to artifacts which may be caused, for example, by endogenously present (or externally added) nucleases.

