## Supplementary material for "Chloroplast nucleoids are highly dynamic in ploidy, number, and structure during angiosperm leaf development": SI Datasets

Supplemental Dataset 1

### Sugar beet (*Beta vulgaris*)

Examples of DAPI stained cells of apex areas to primordia leaflets up to 2.5 mm

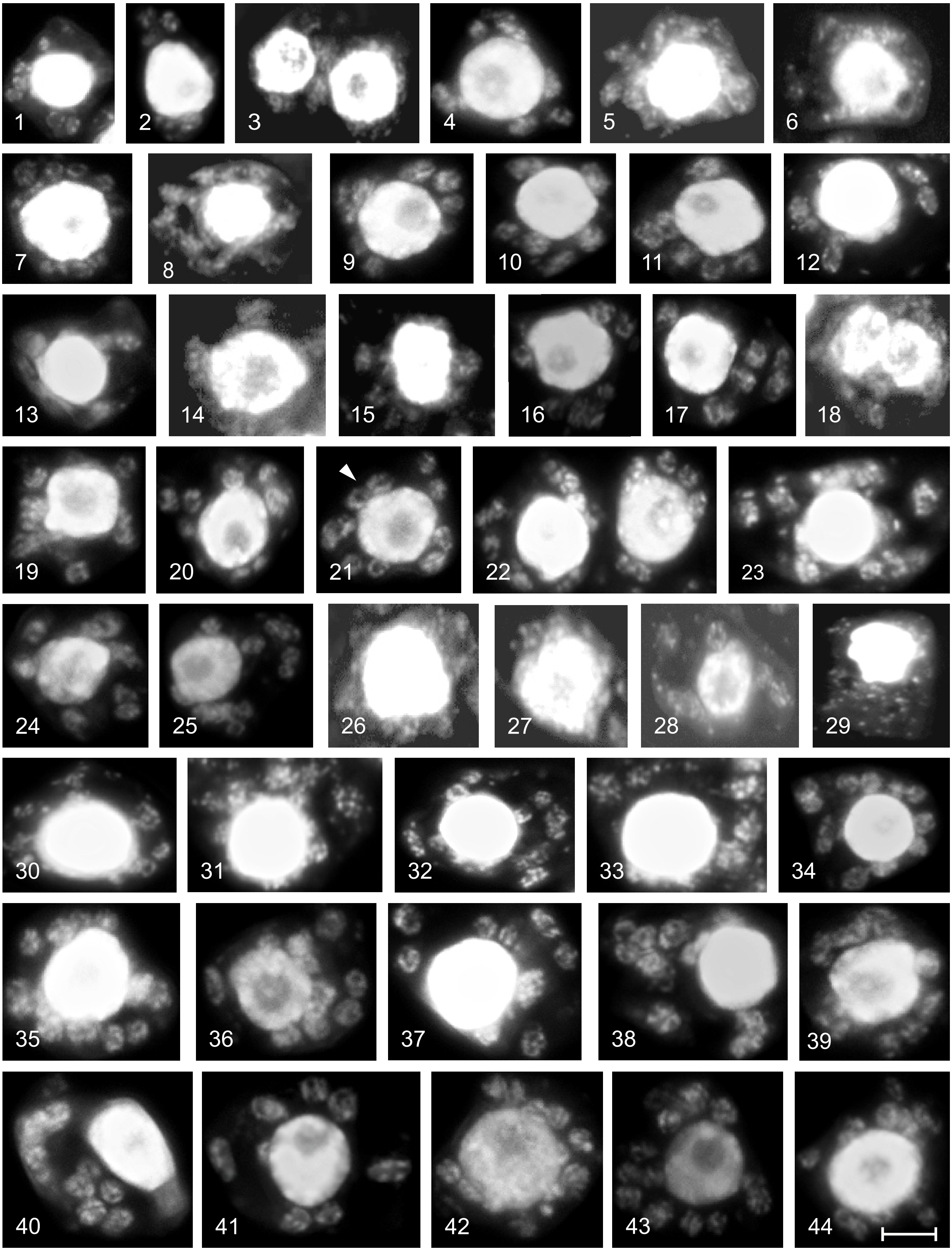

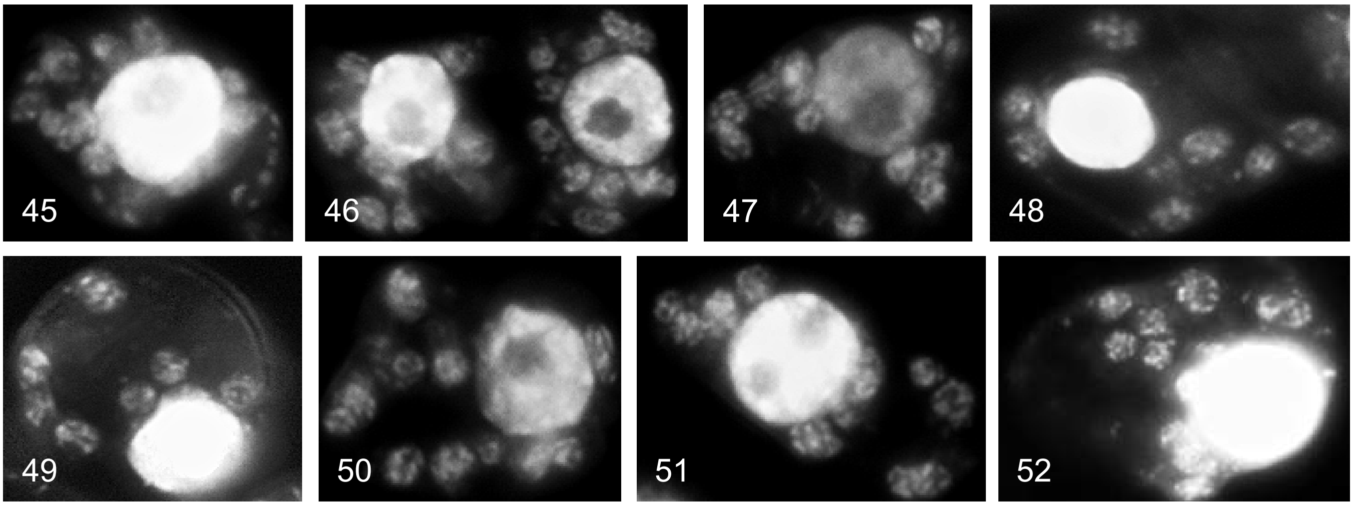

Examples of DAPI stained cells of leaflets between 4 and 16 mm

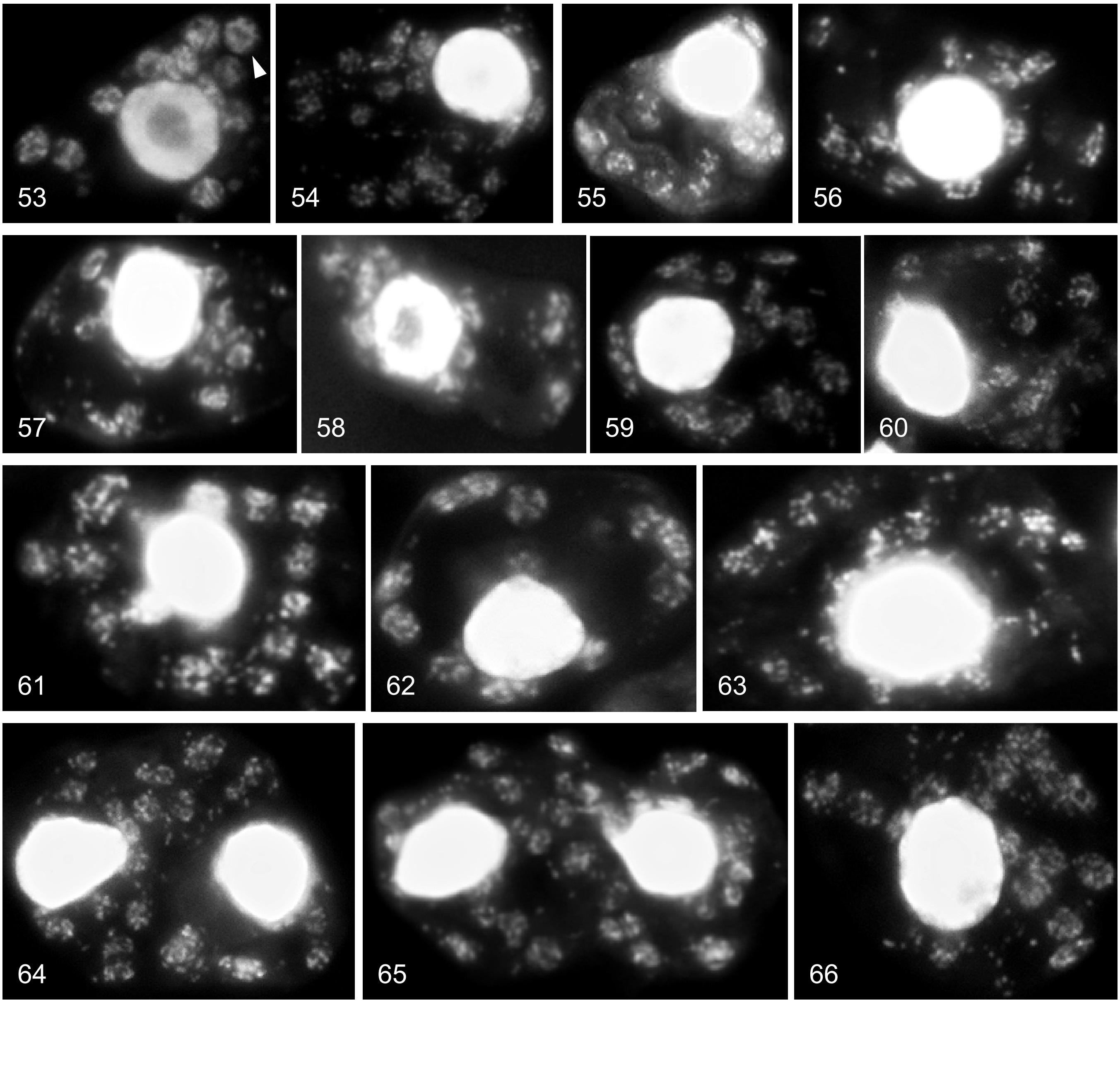

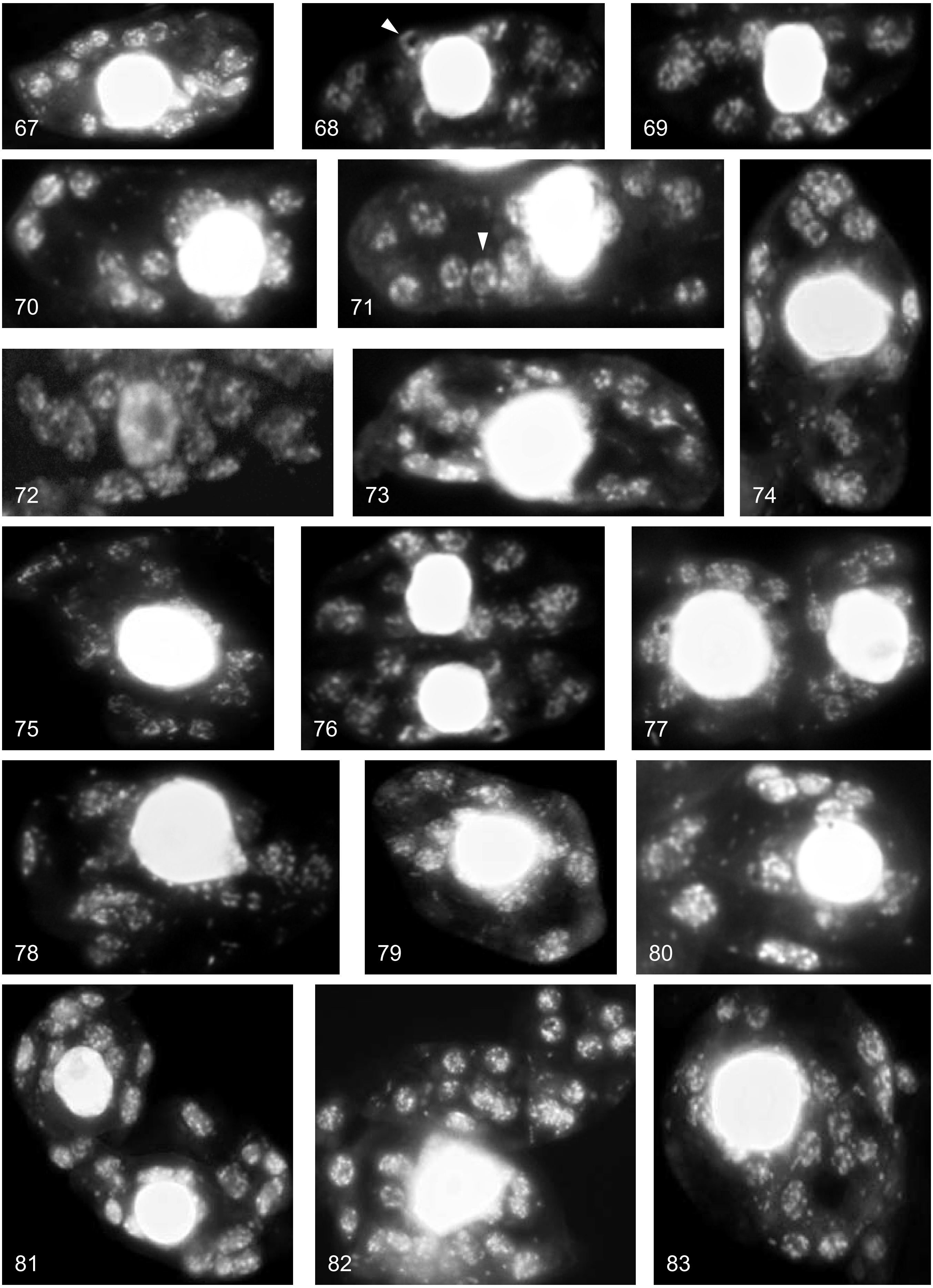

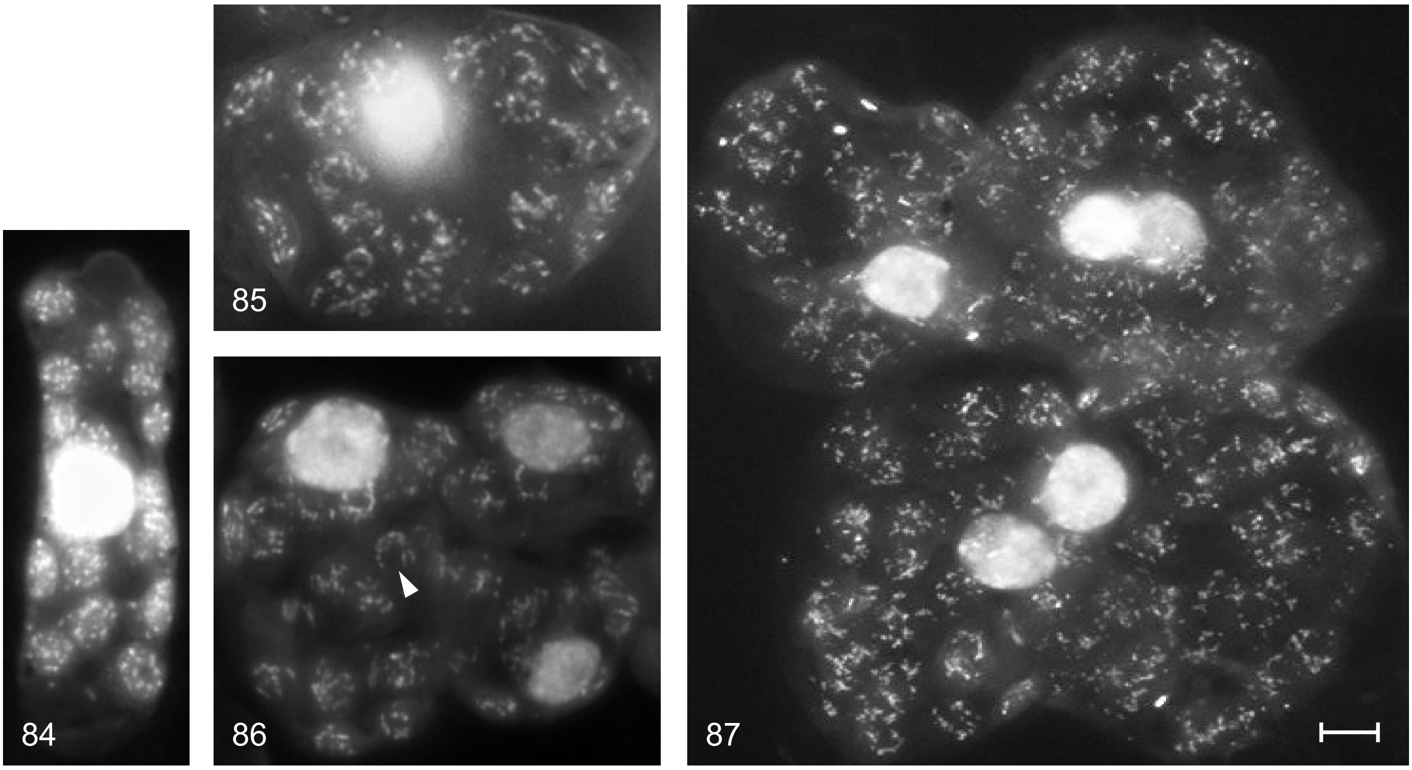

Examples of DAPI stained cells of leaflets between 1.5 and 3 cm

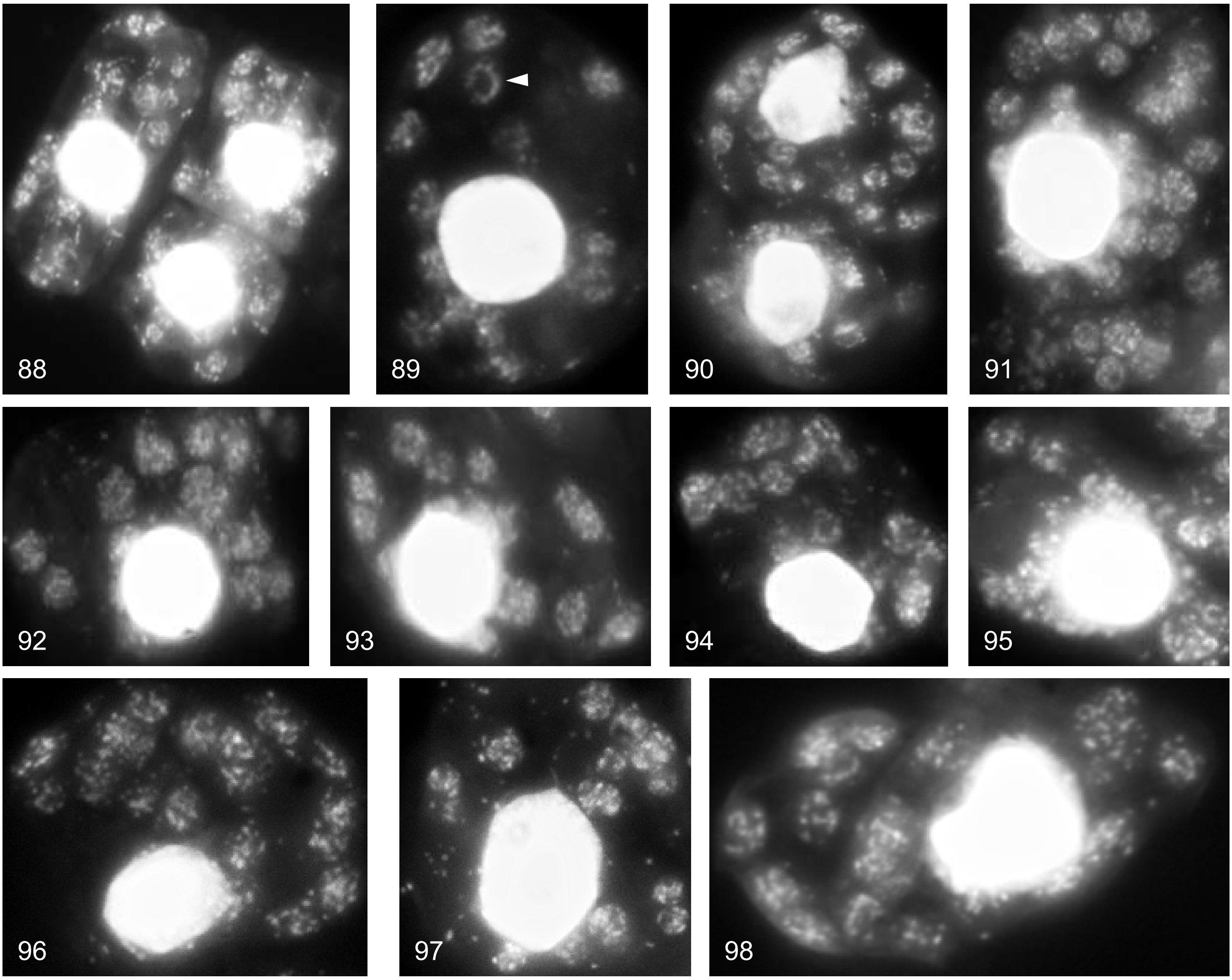

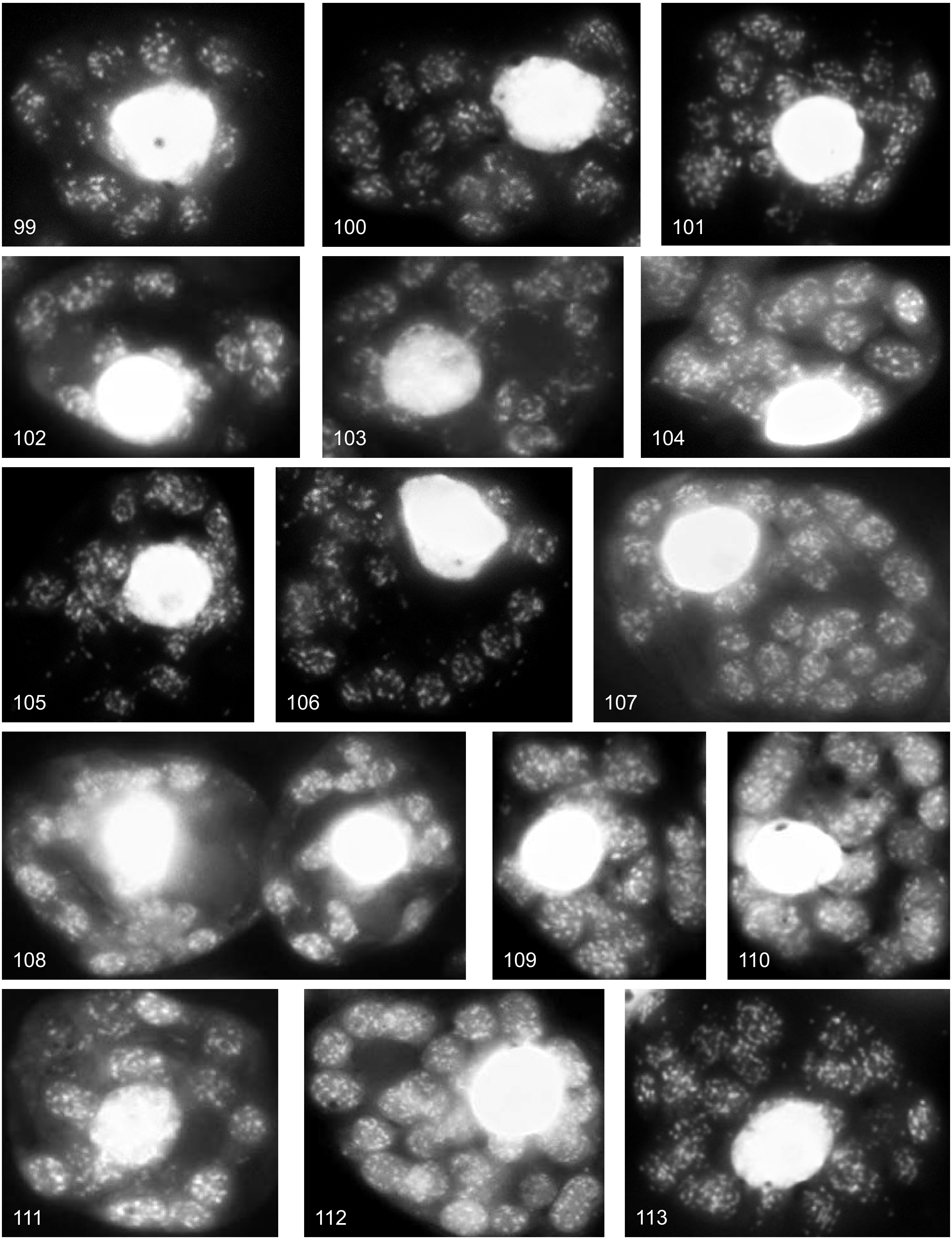

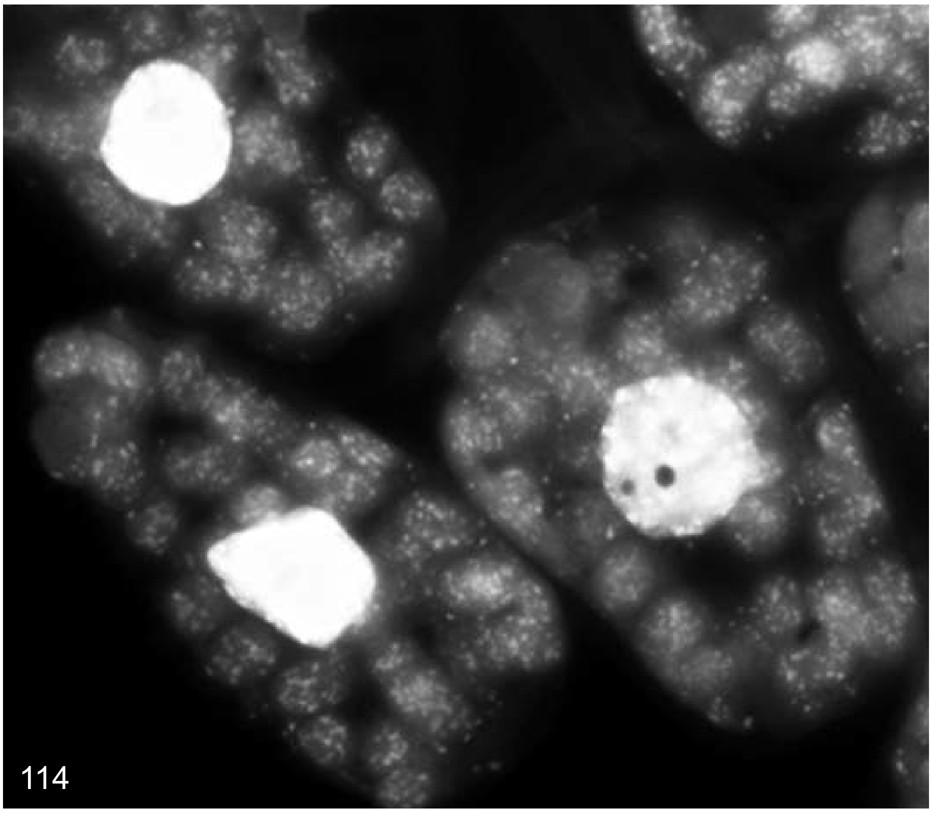

Examples of DAPI stained cells of leaves between 3 and 9 cm, and some of mature and early ageing leaves for comparison with picture series presented in Golczyk et al. (2014)

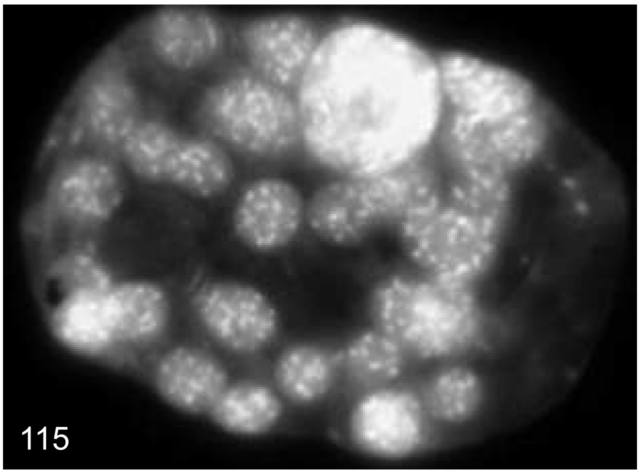

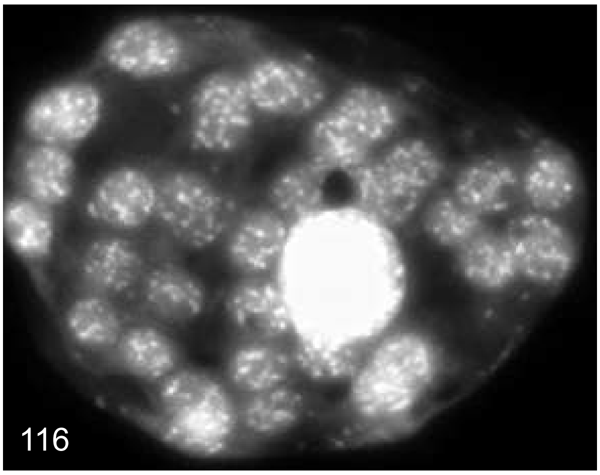

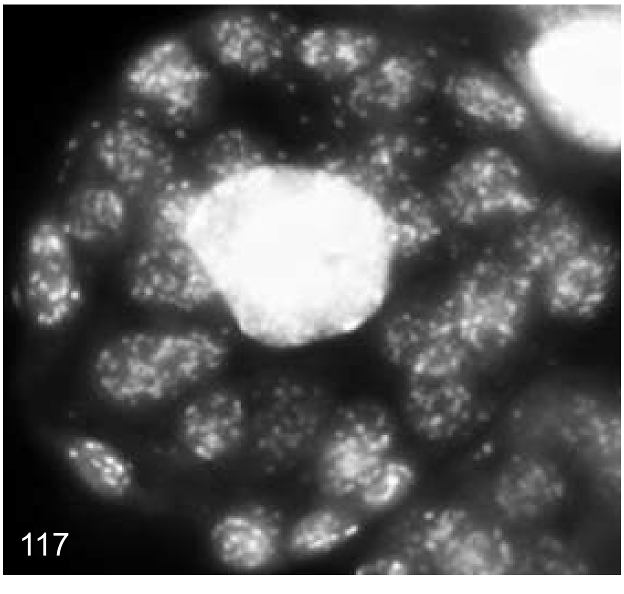

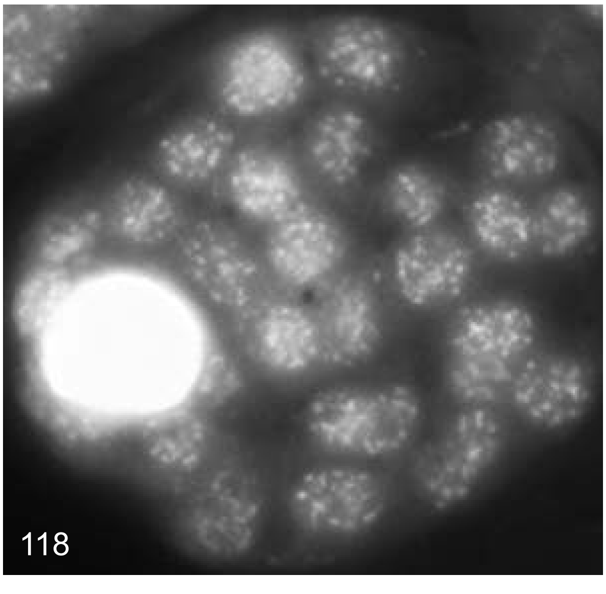

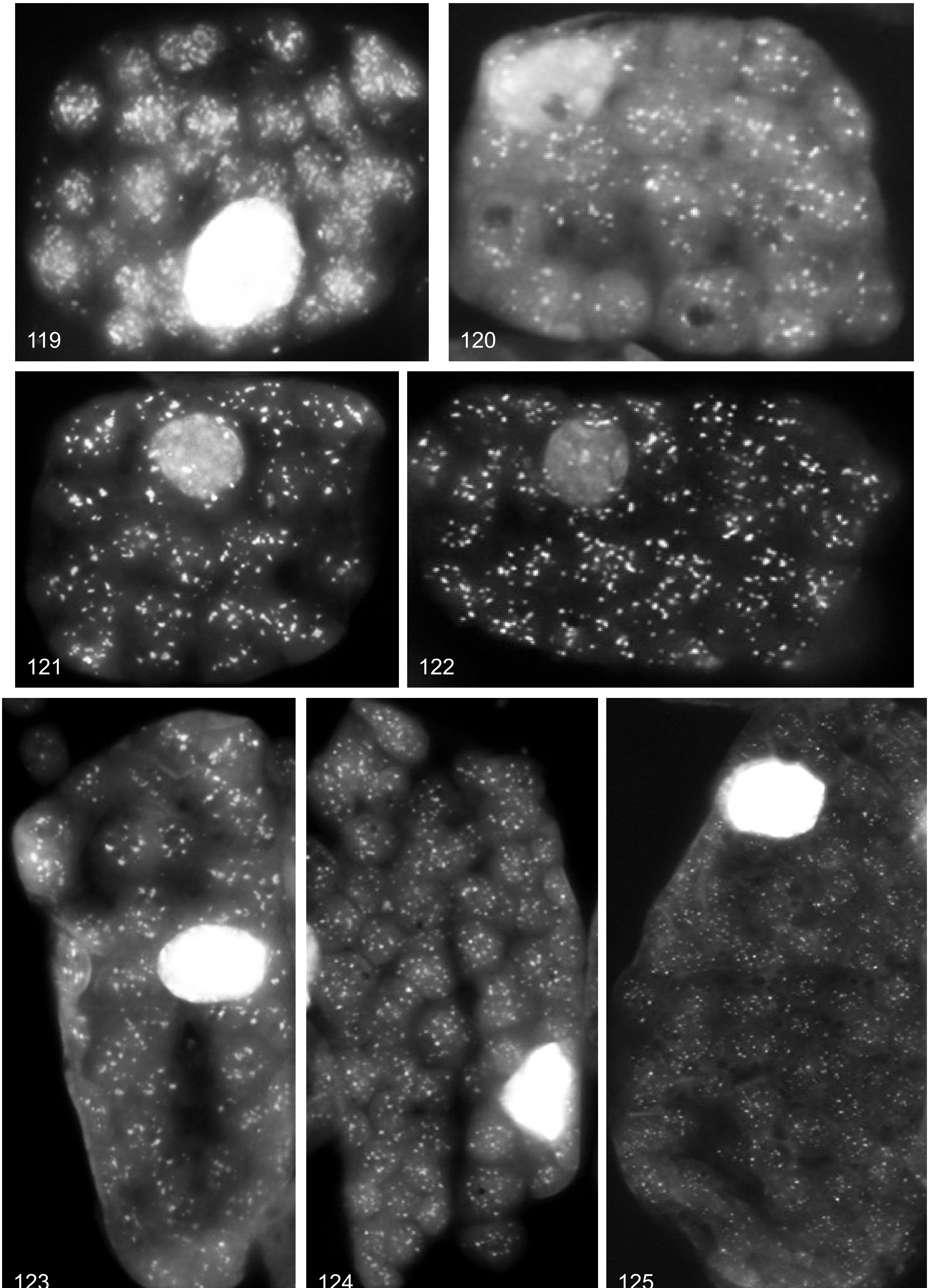
119
120
121
122
123
124
125

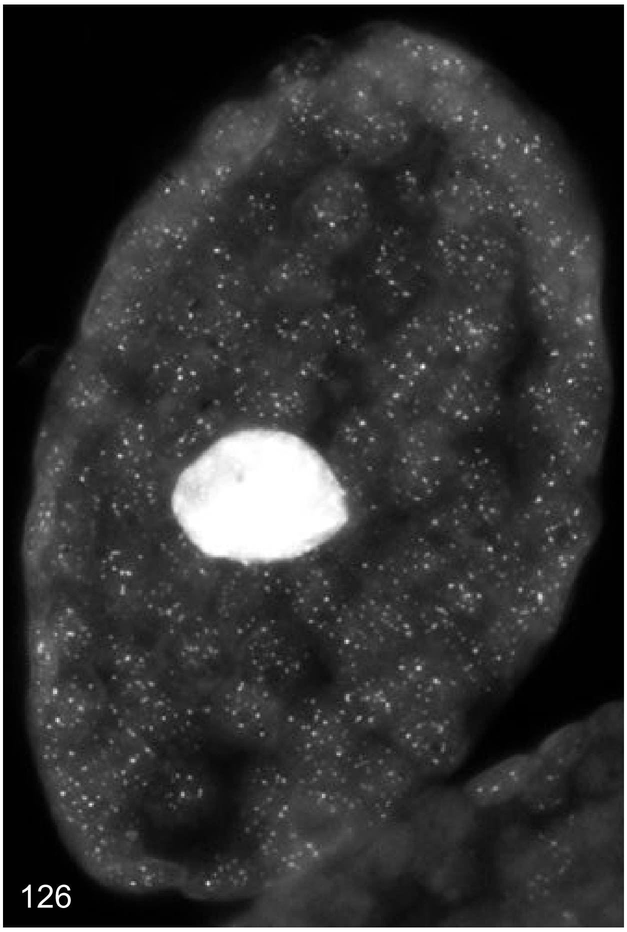

126

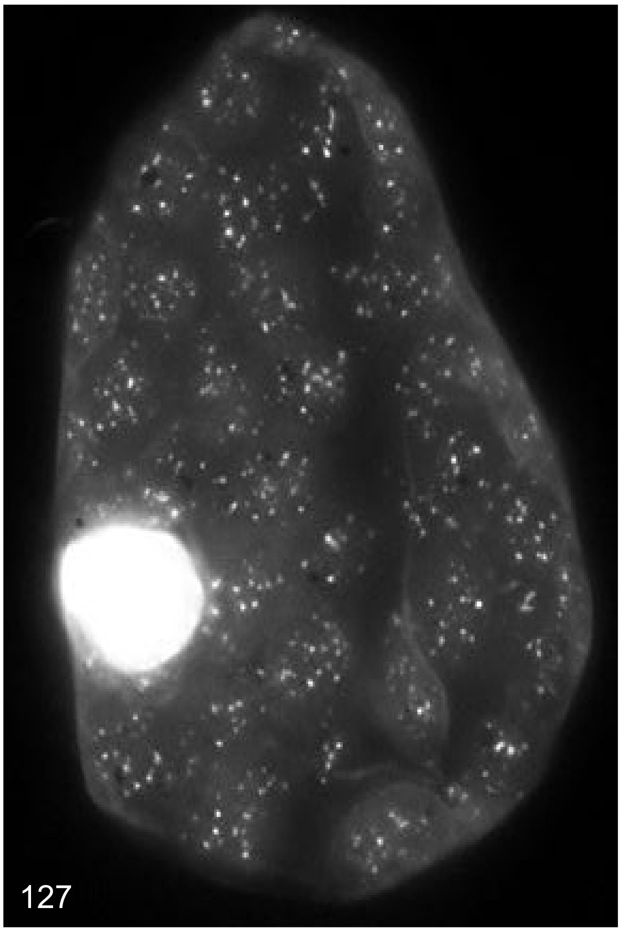

127

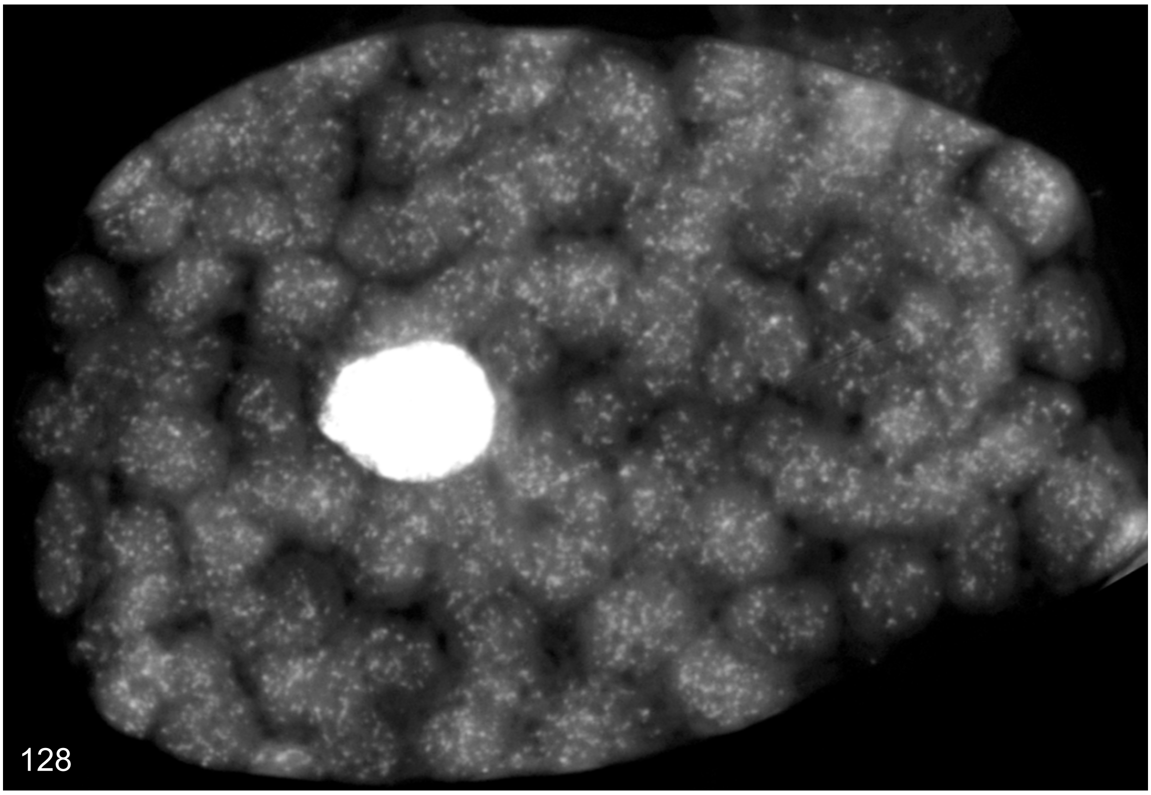

128

Supplemental Dataset 2

### *Arabidopsis thaliana*

Examples of DAPI stained cells of apex areas to primordia leaflets up to 1 mm

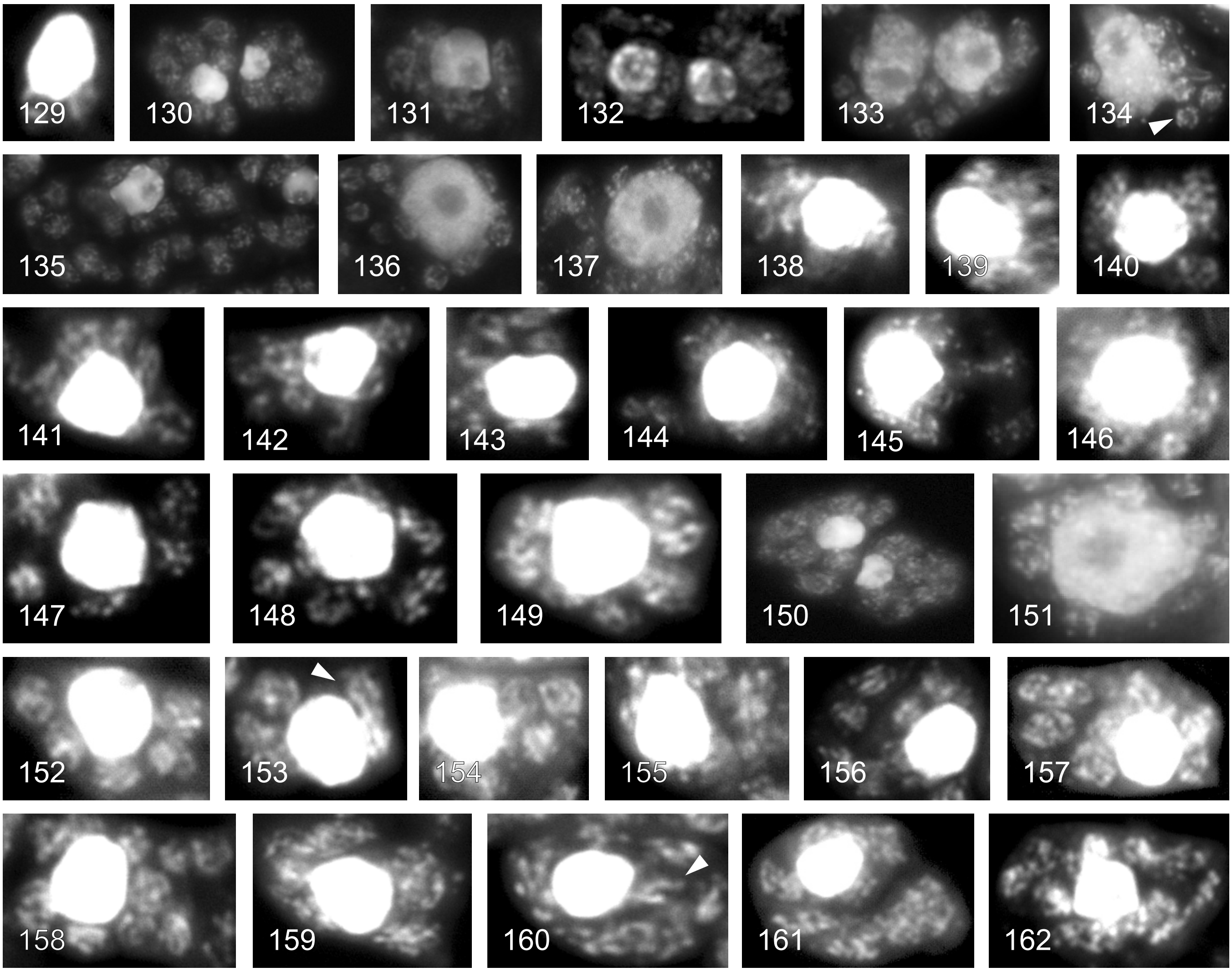

Examples of DAPI stained cells of leaflets between 1.5 and 3 mm

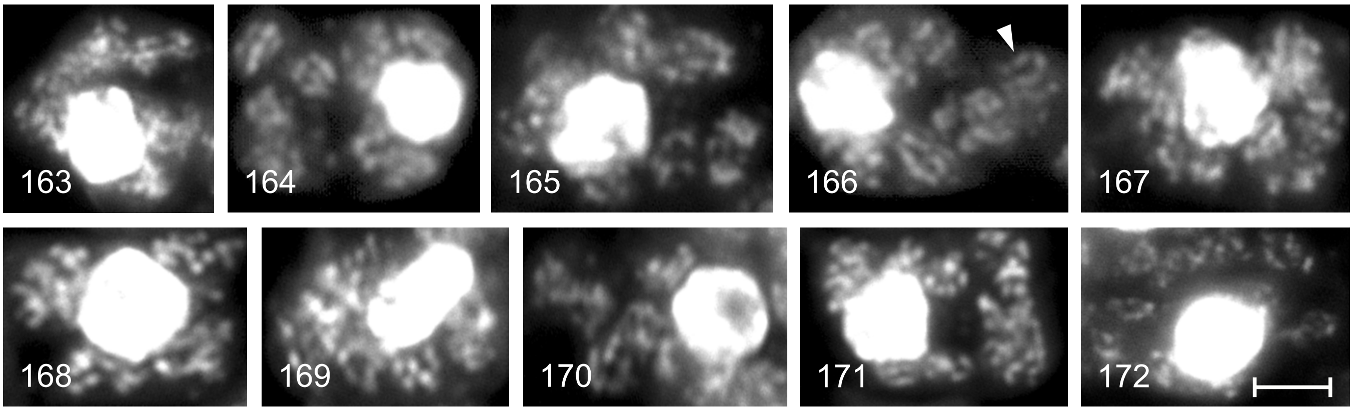

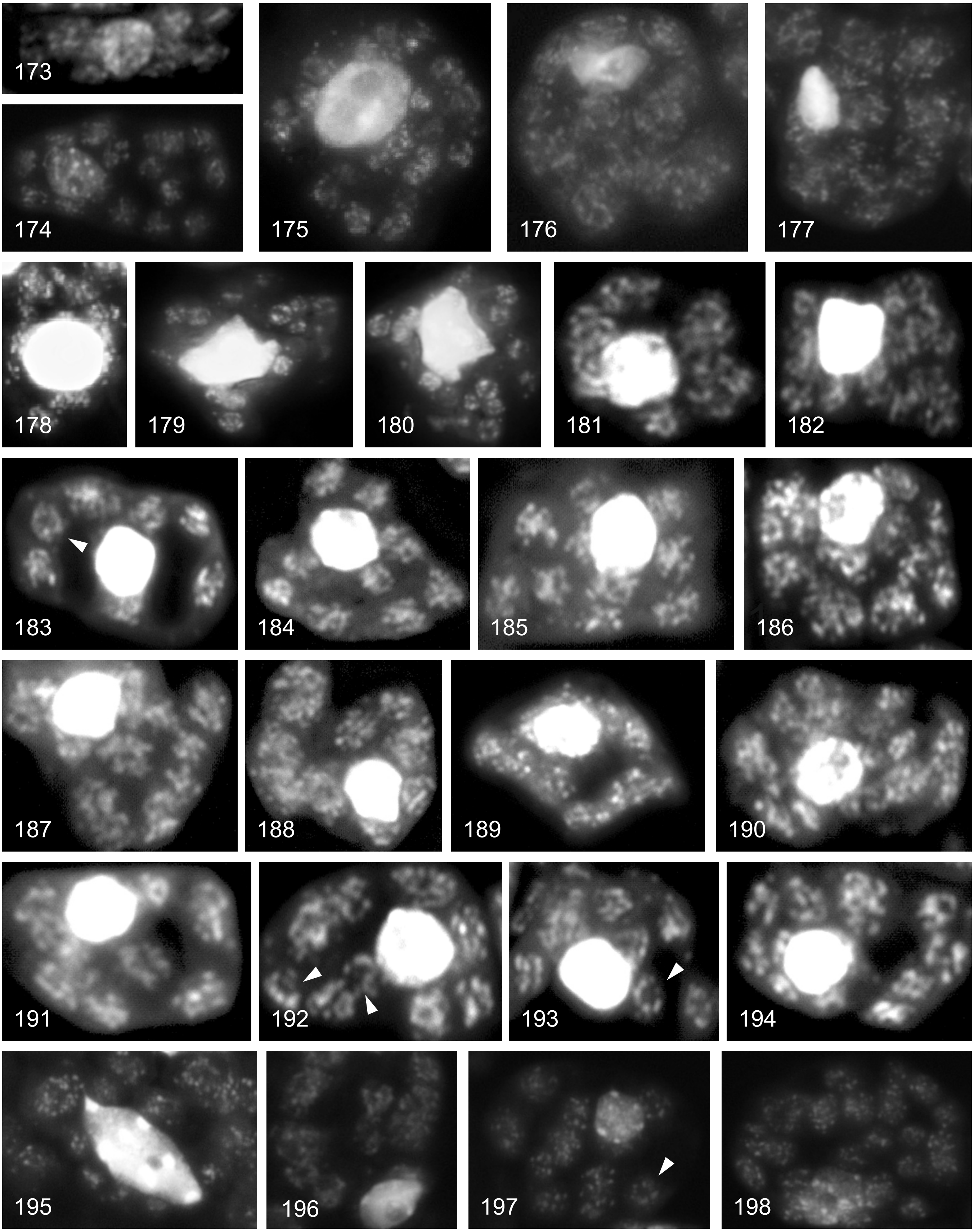

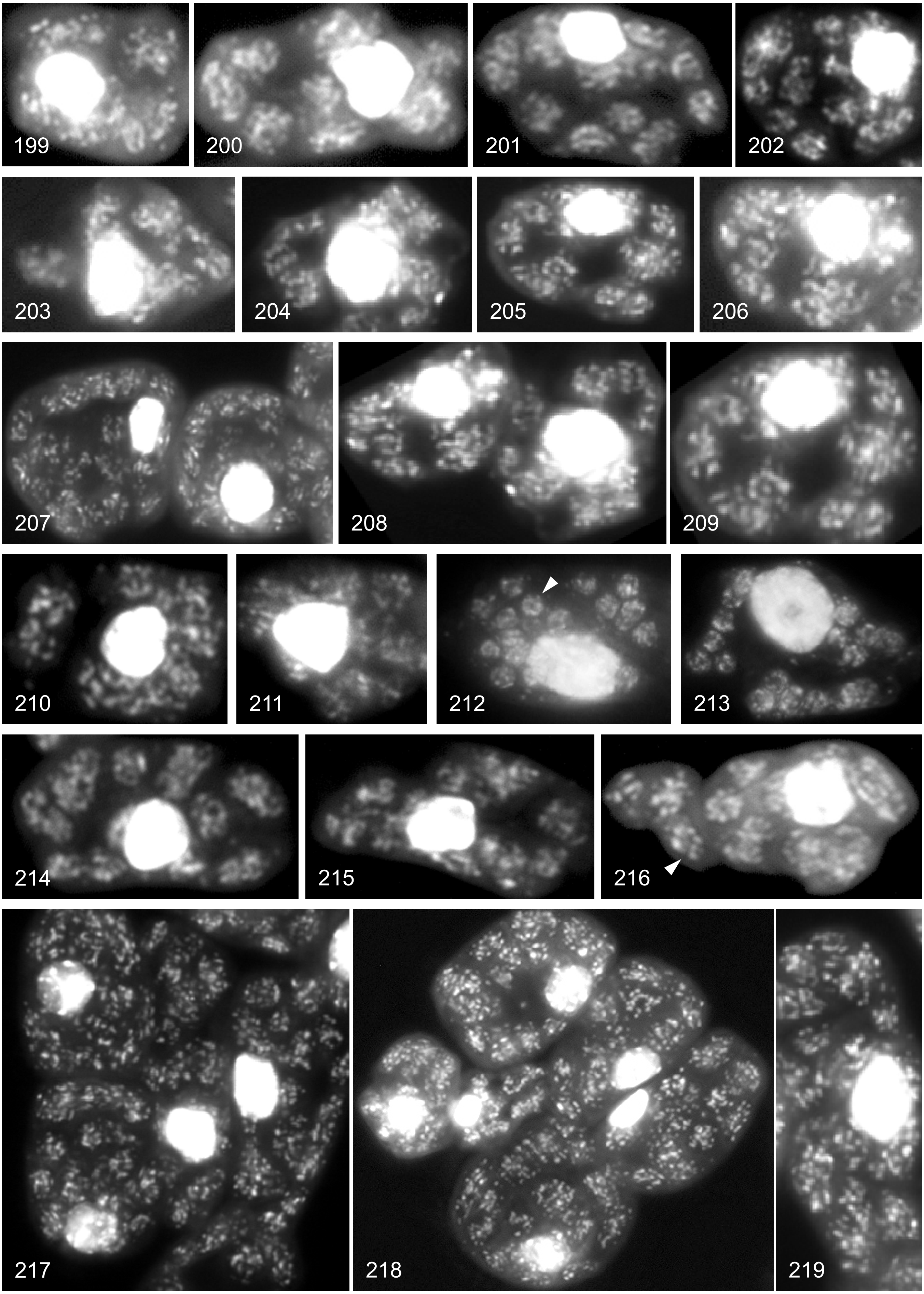

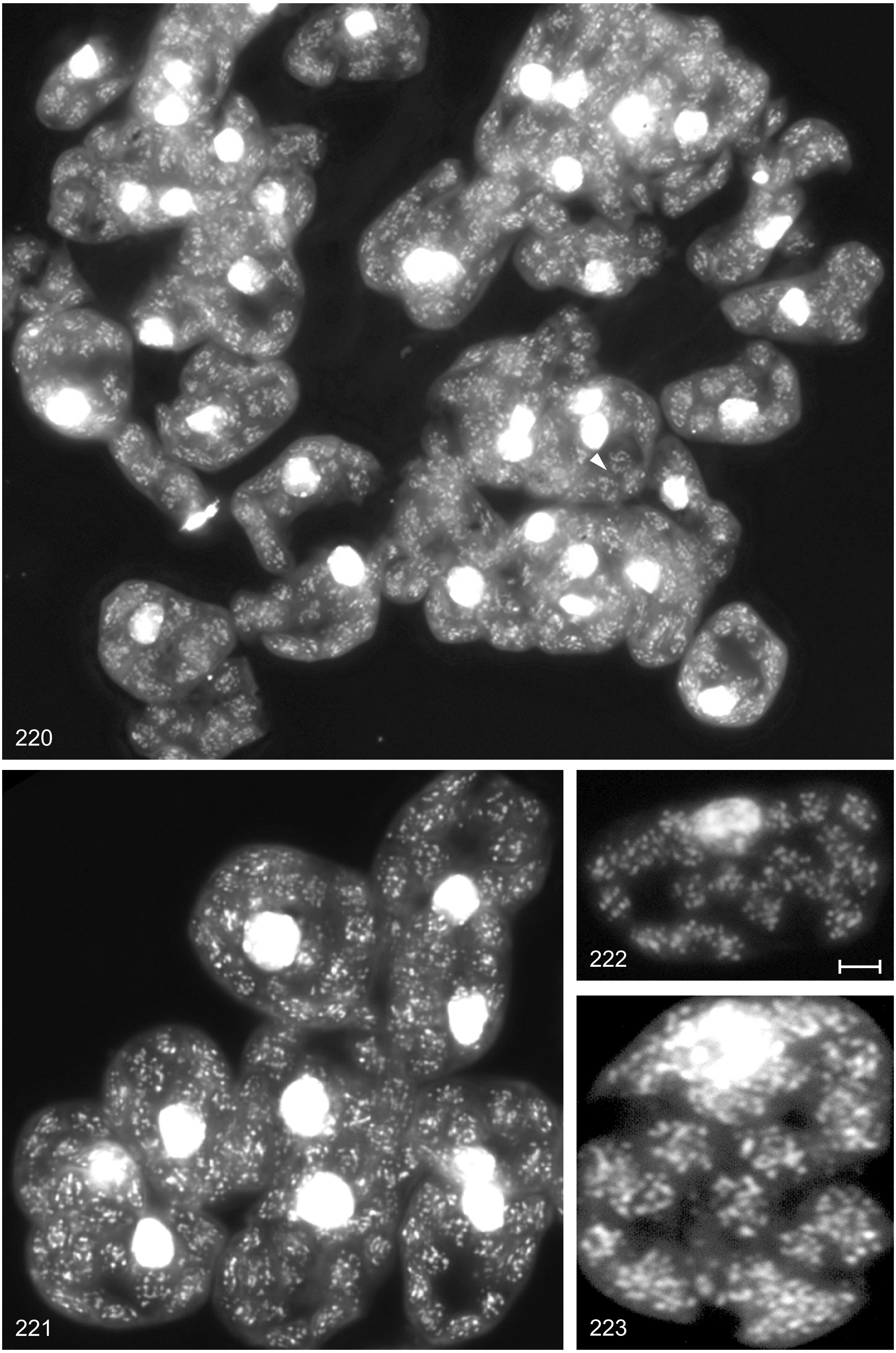
220
221
222
223

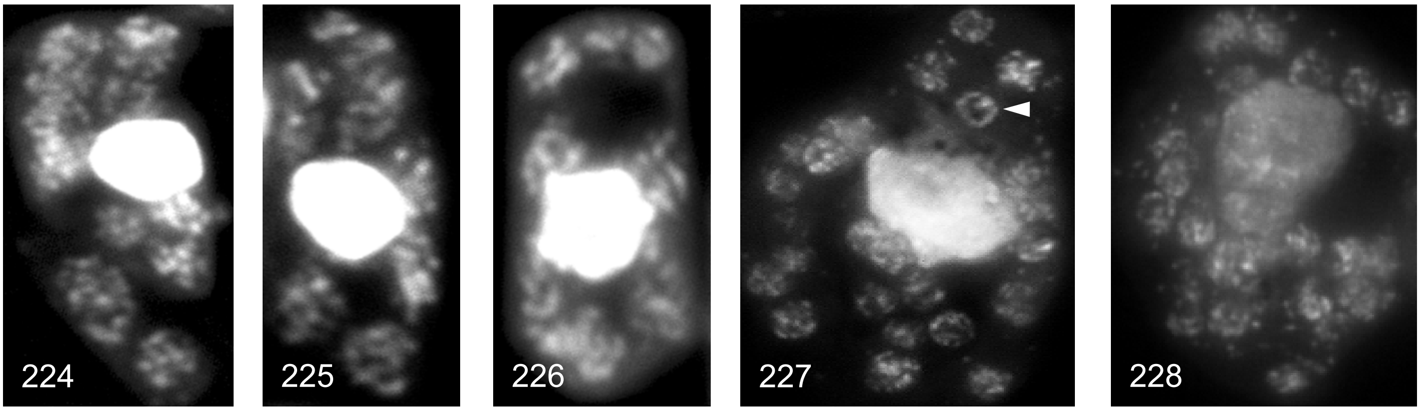

Examples of DAPI stained cells of leaflets between 2.5 and 4 mm

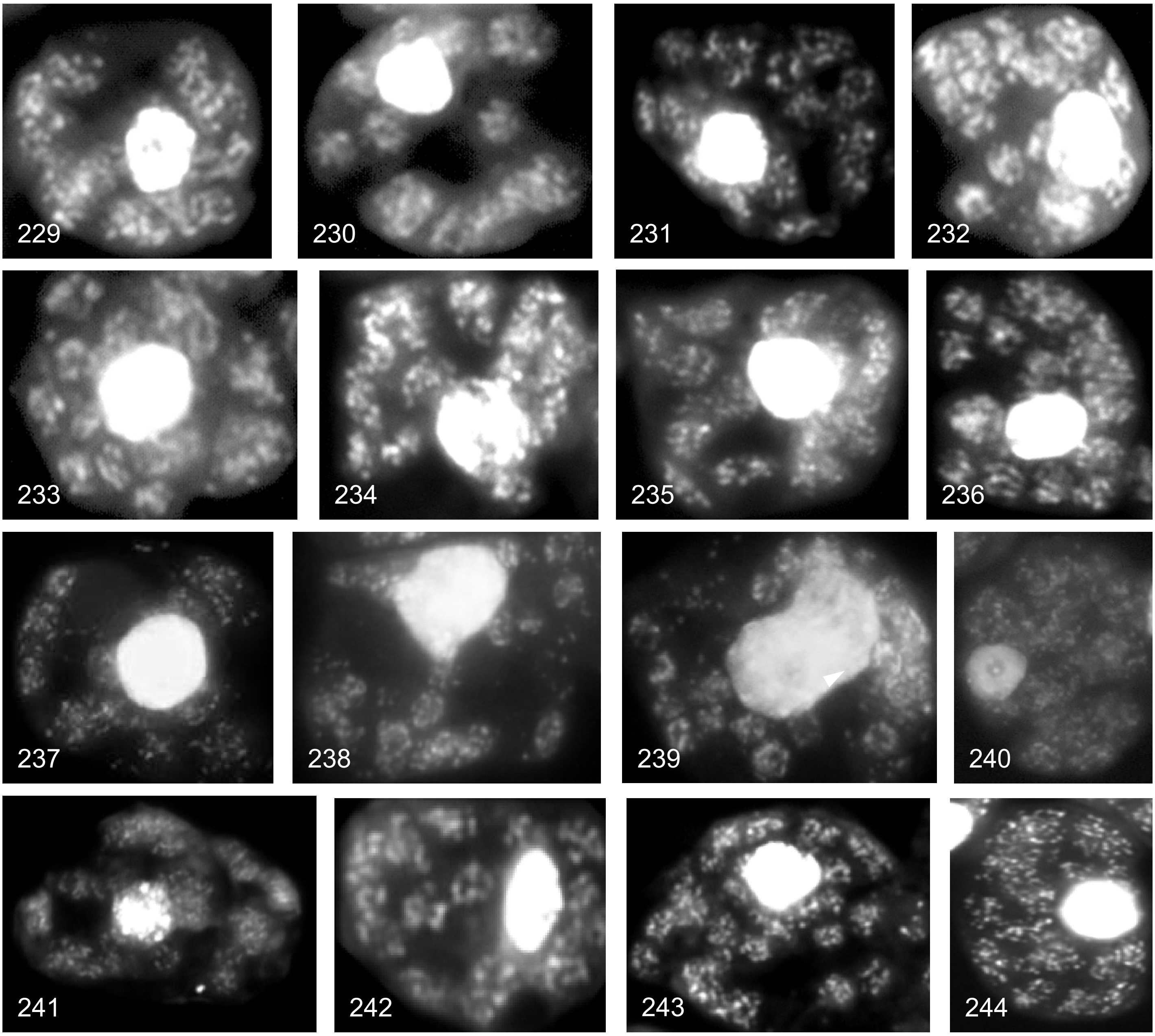

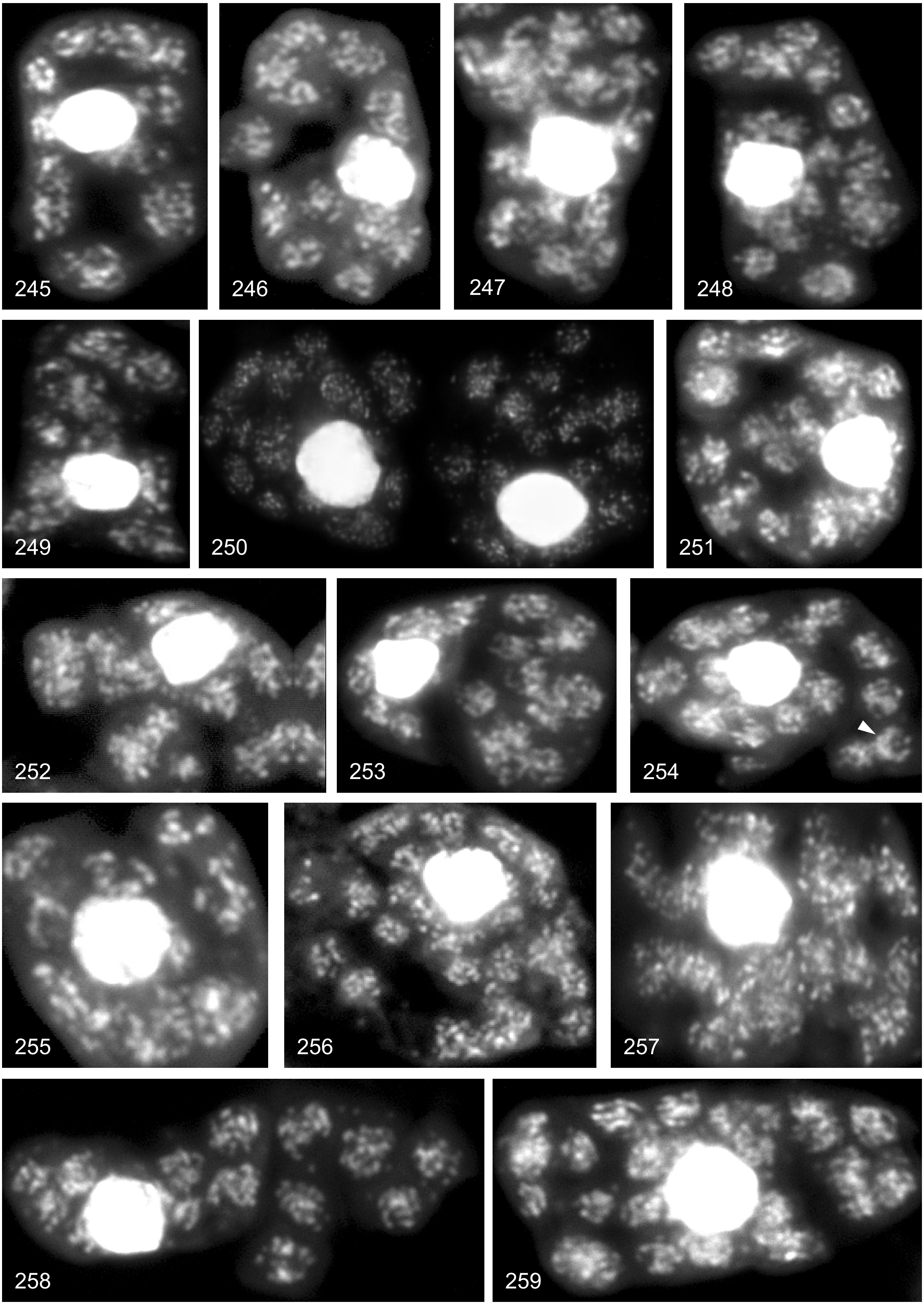

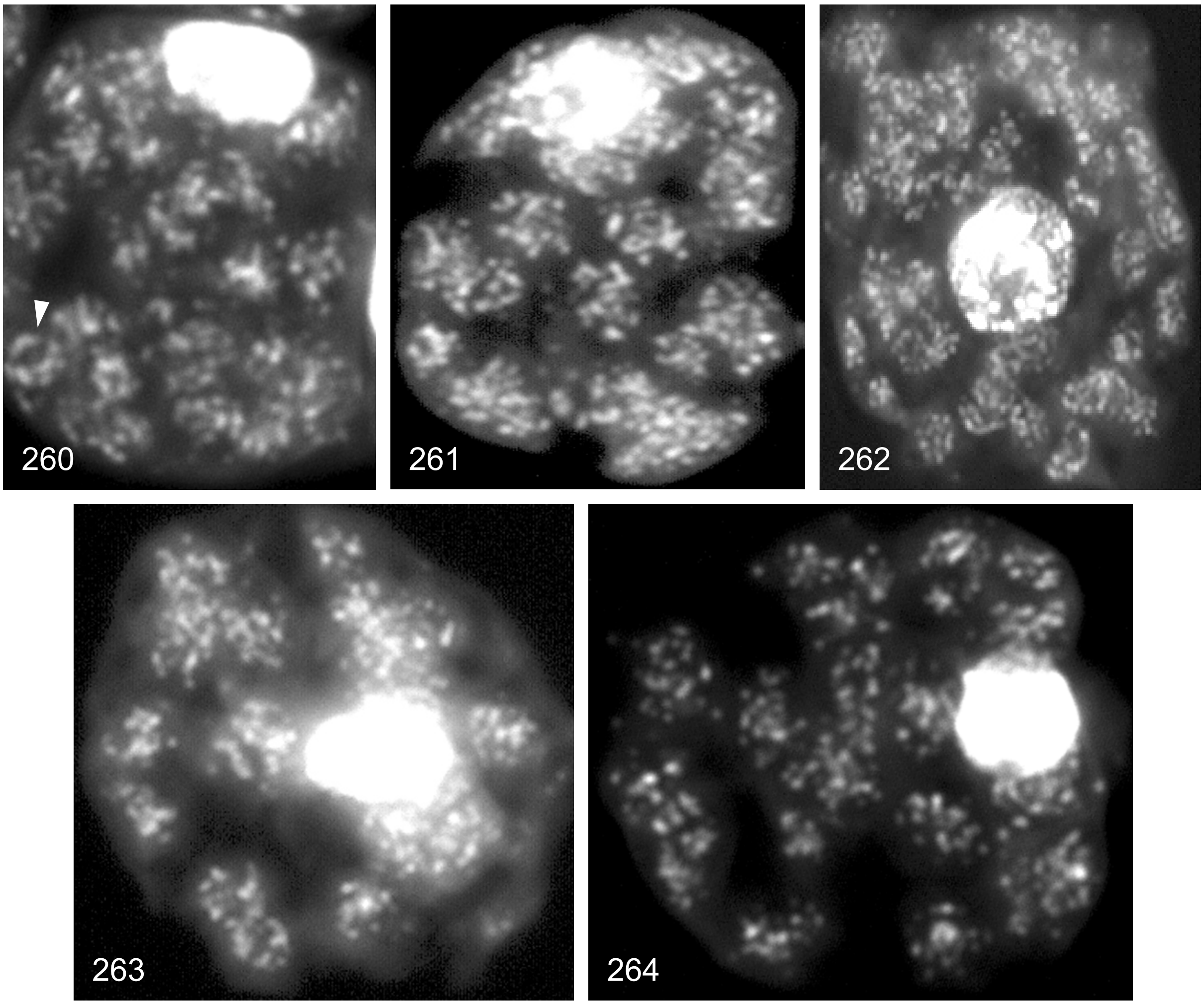

Examples of DAPI stained cells of leaflets between 4 and 8 mm

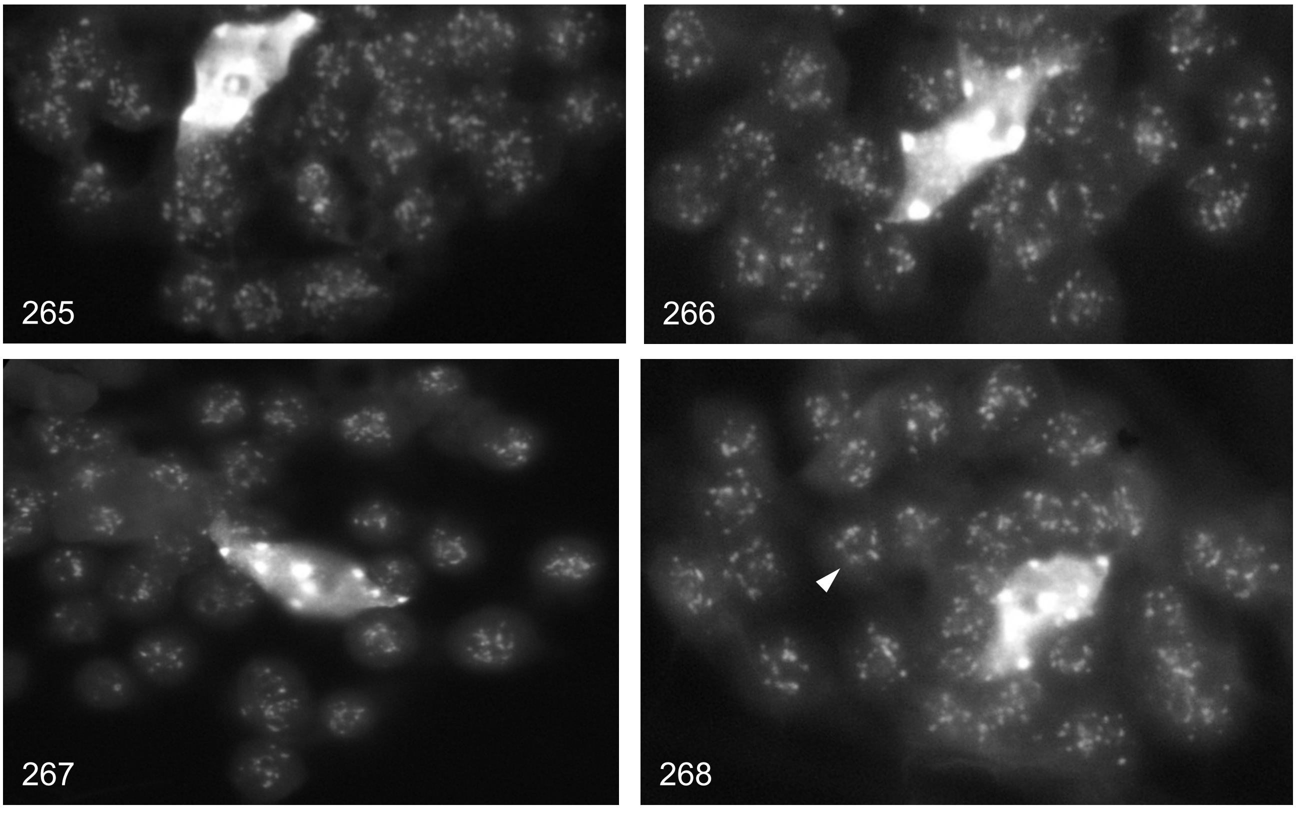

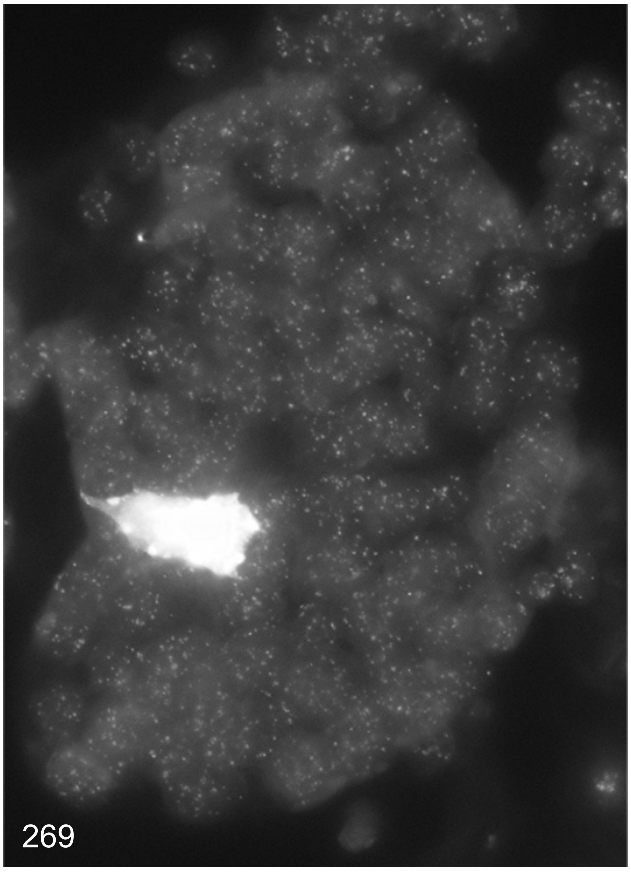
269

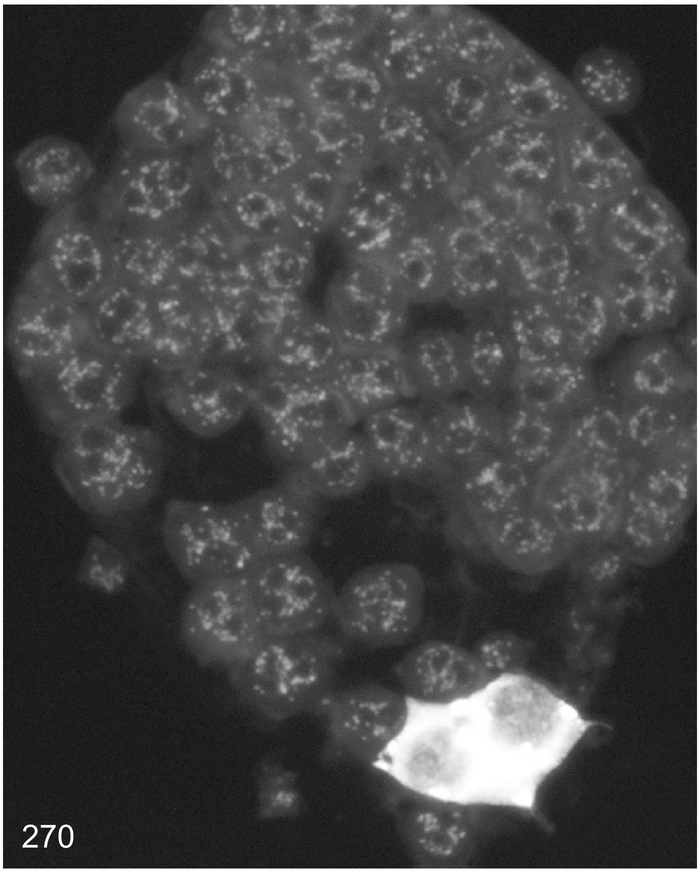
270

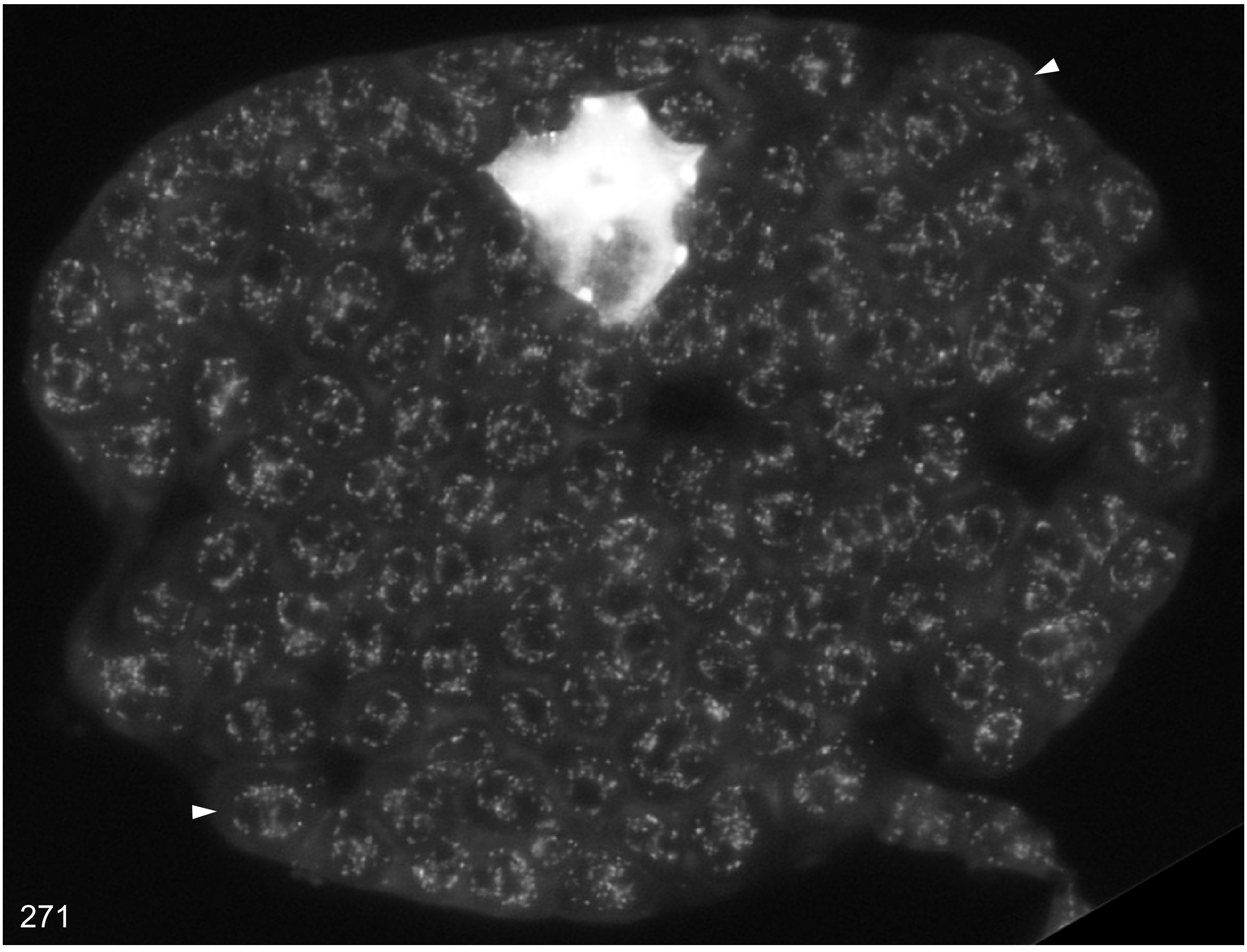
271

Supplemental Dataset 3

### Tobacco (*Nicotiana tabacum*)

Examples of DAPI stained cells of apex areas to primordia leaflets up to 2 mm

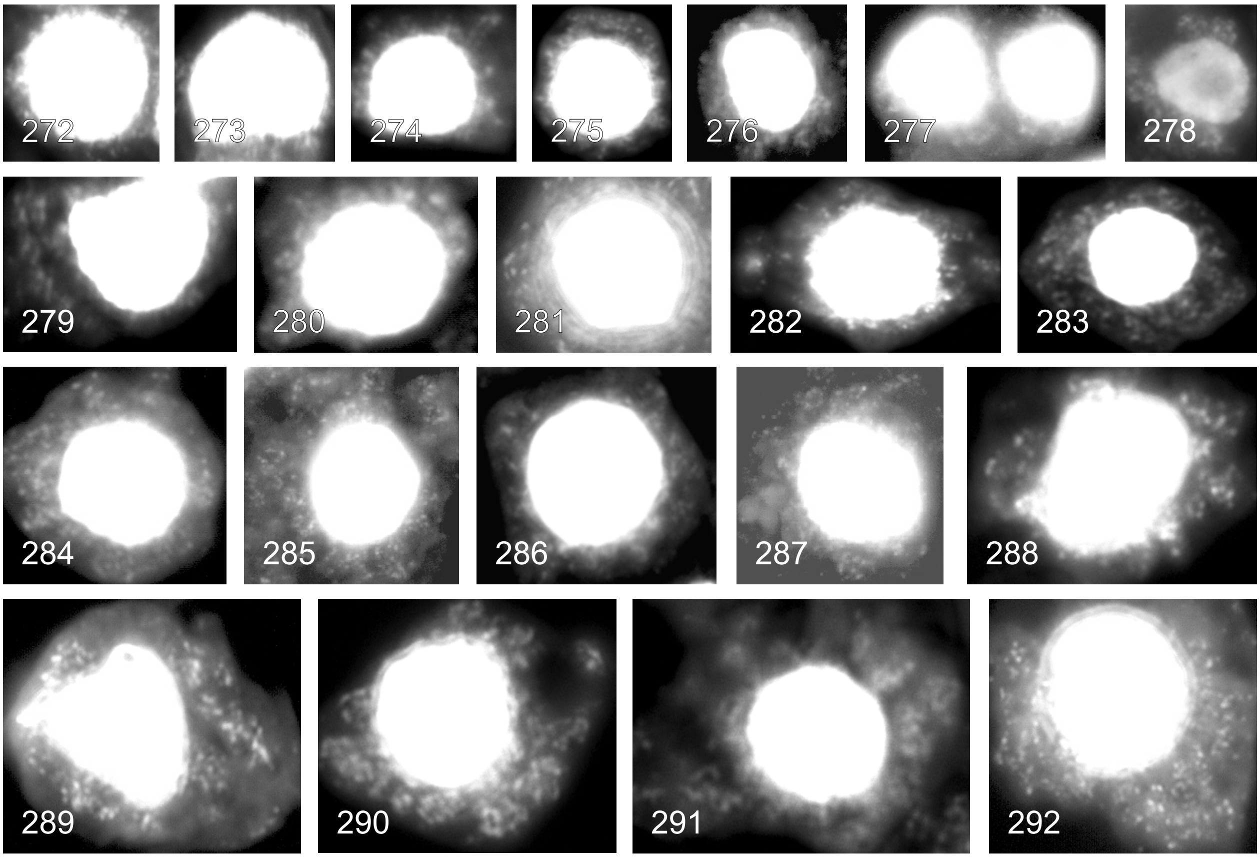

Examples of DAPI stained cells of leaflets between 2 and 10 mm

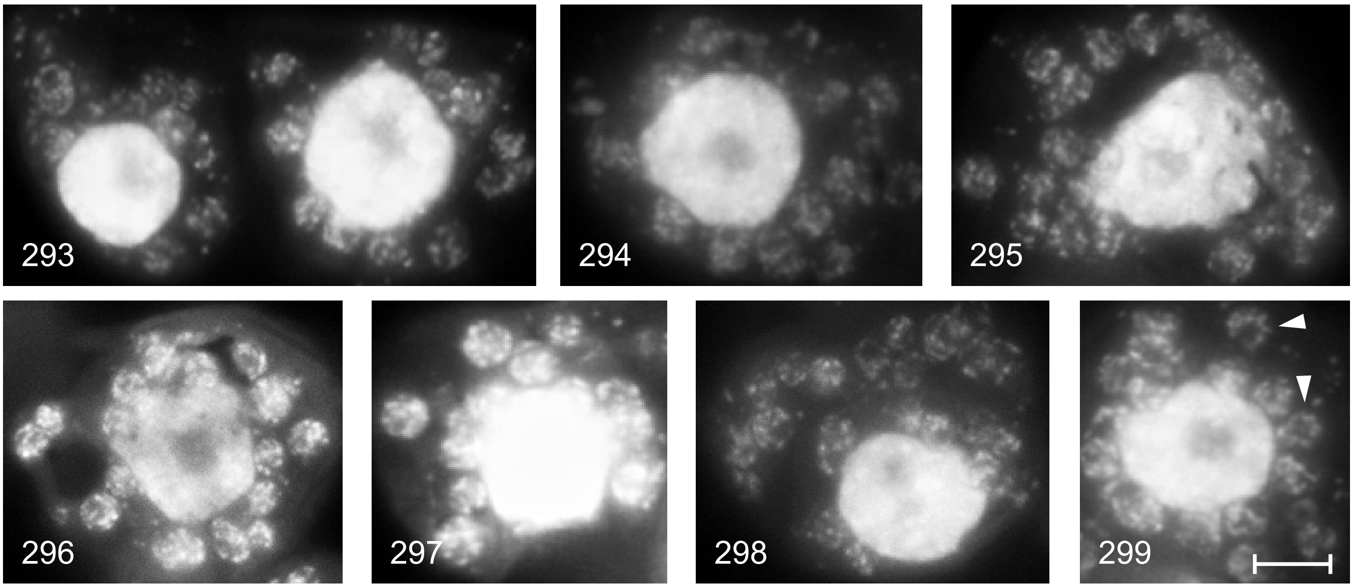

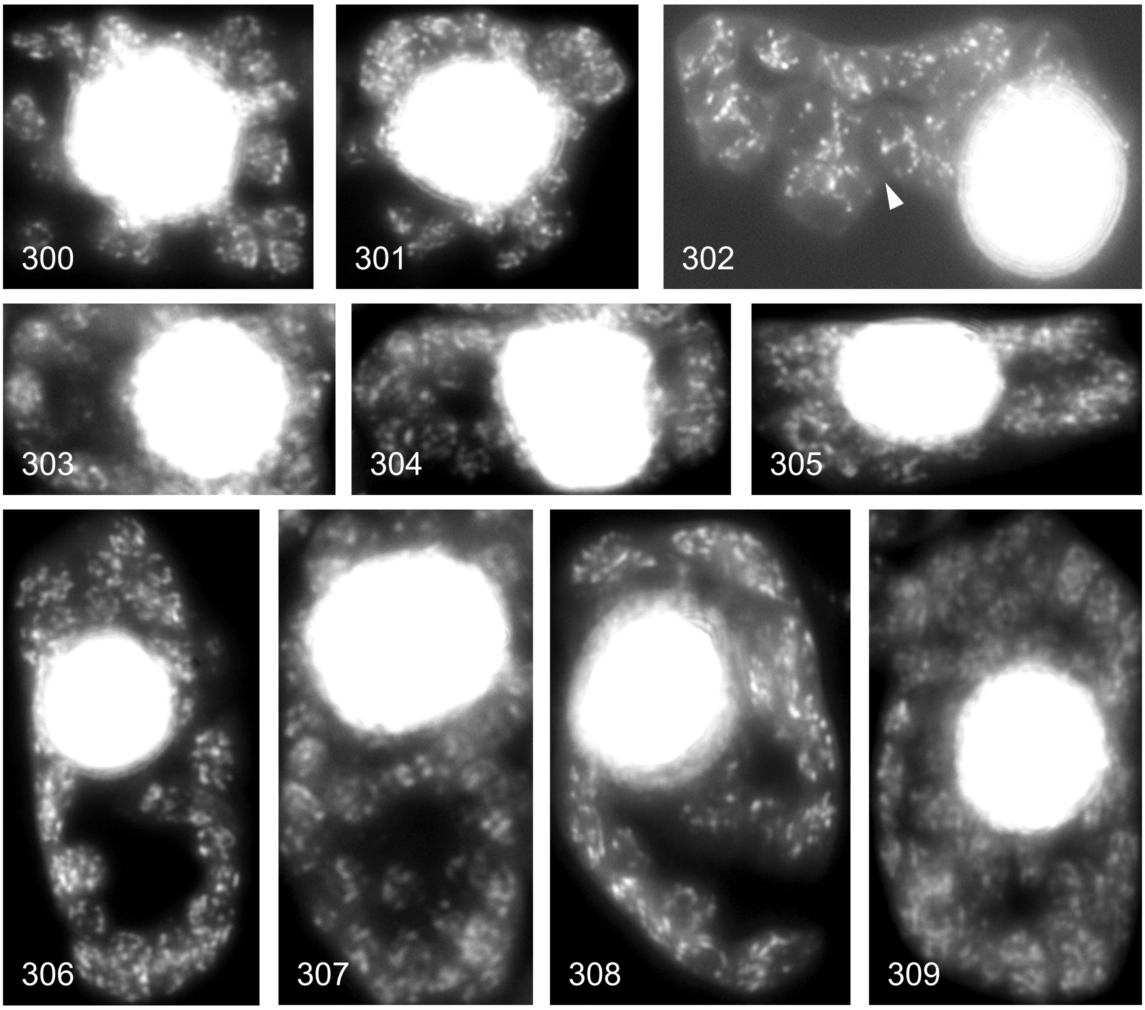

Examples of DAPI stained cells of leaflets between 1 and 2.3 cm

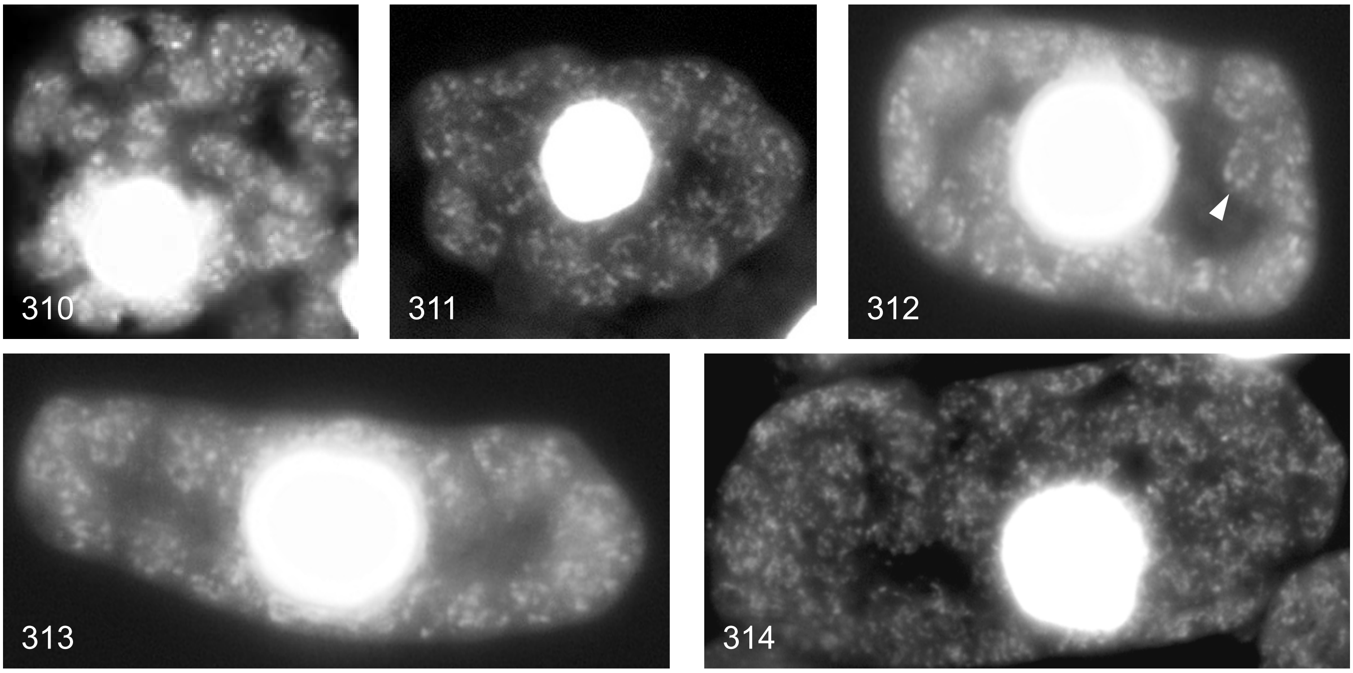

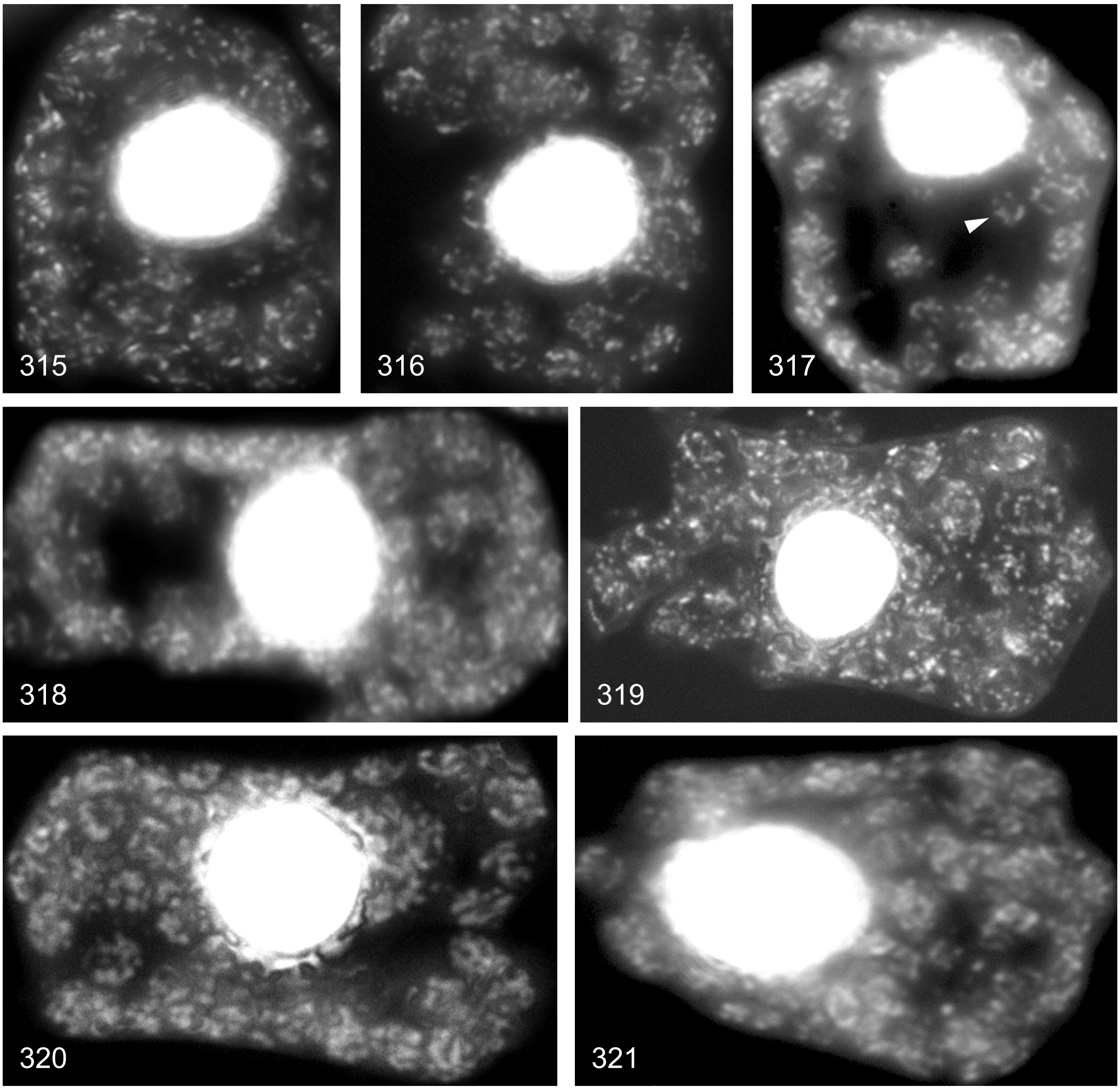

Examples of DAPI stained cells of leaflets between 2 and 4 cm

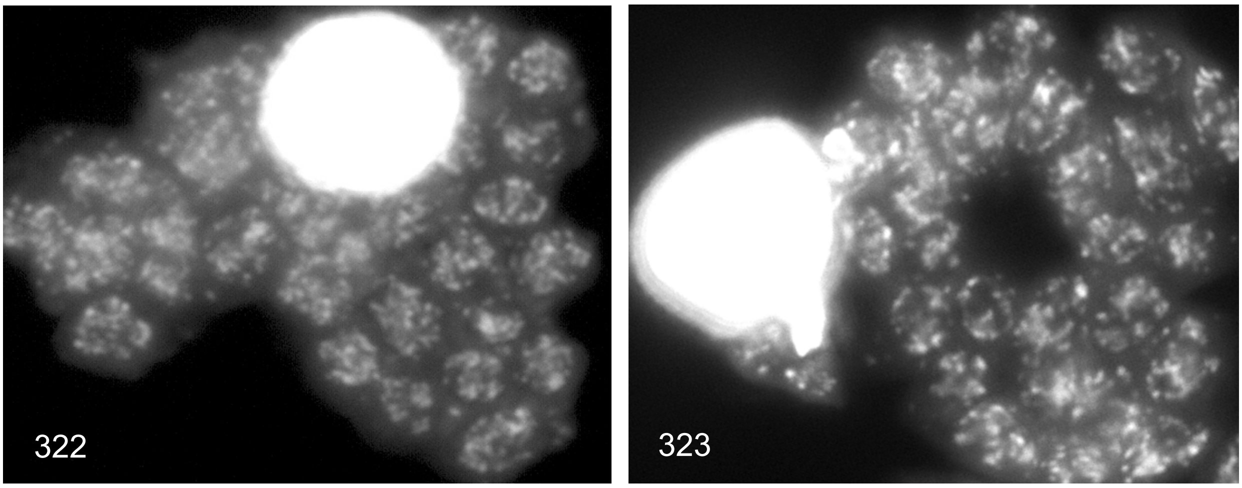

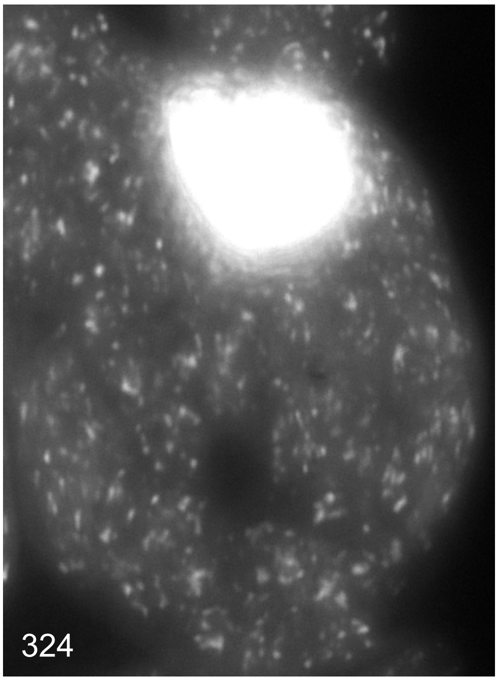
324
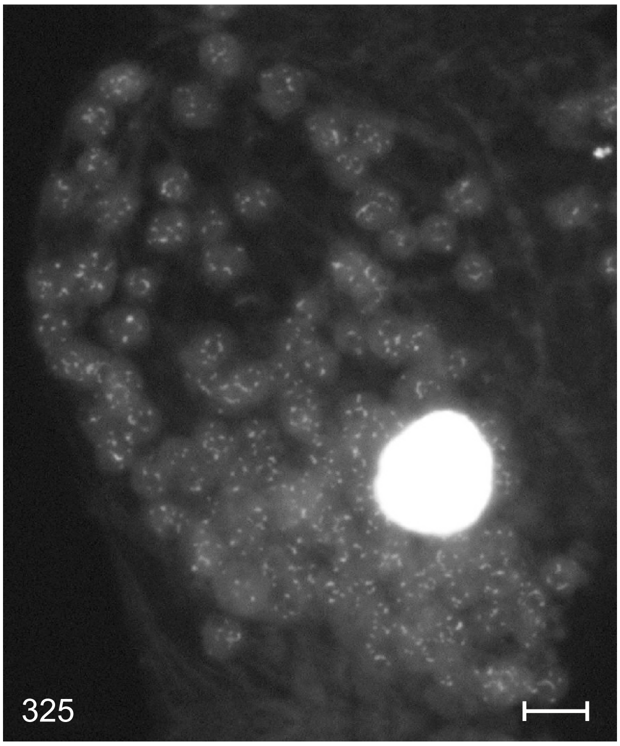
325
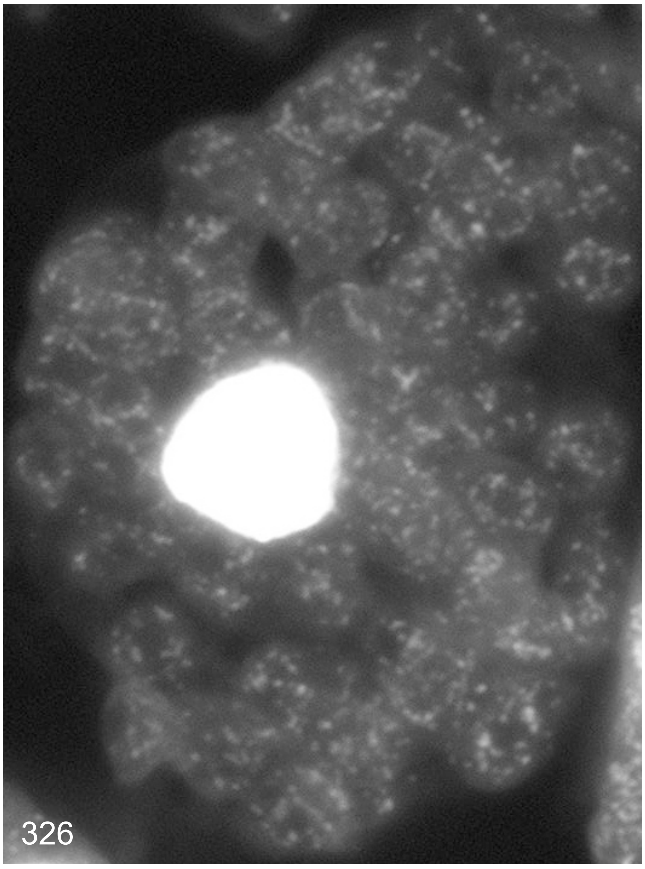
326
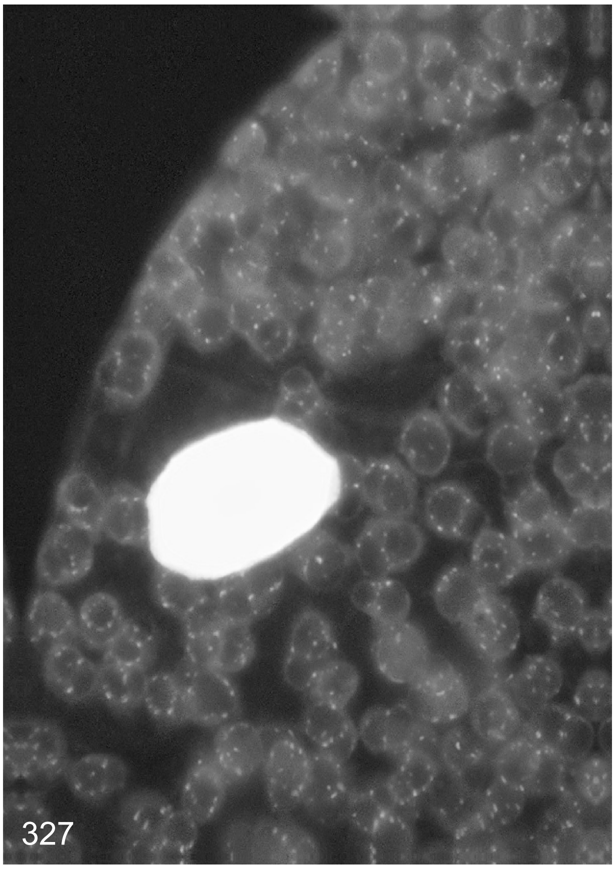
327

Examples of DAPI stained cells of leaflets between 4 and 9 cm

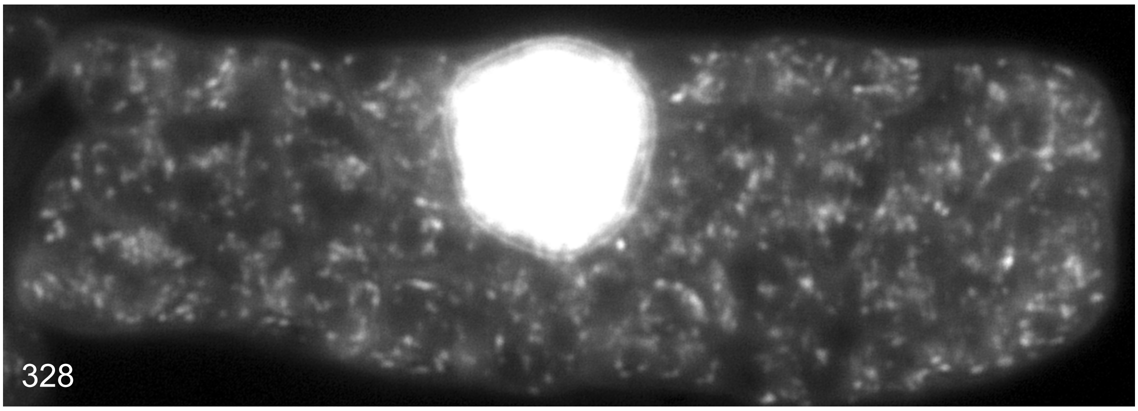

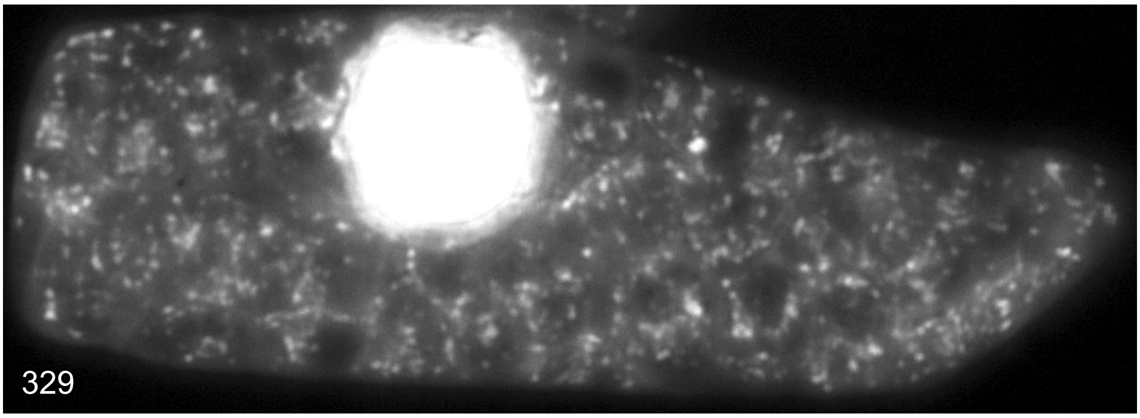

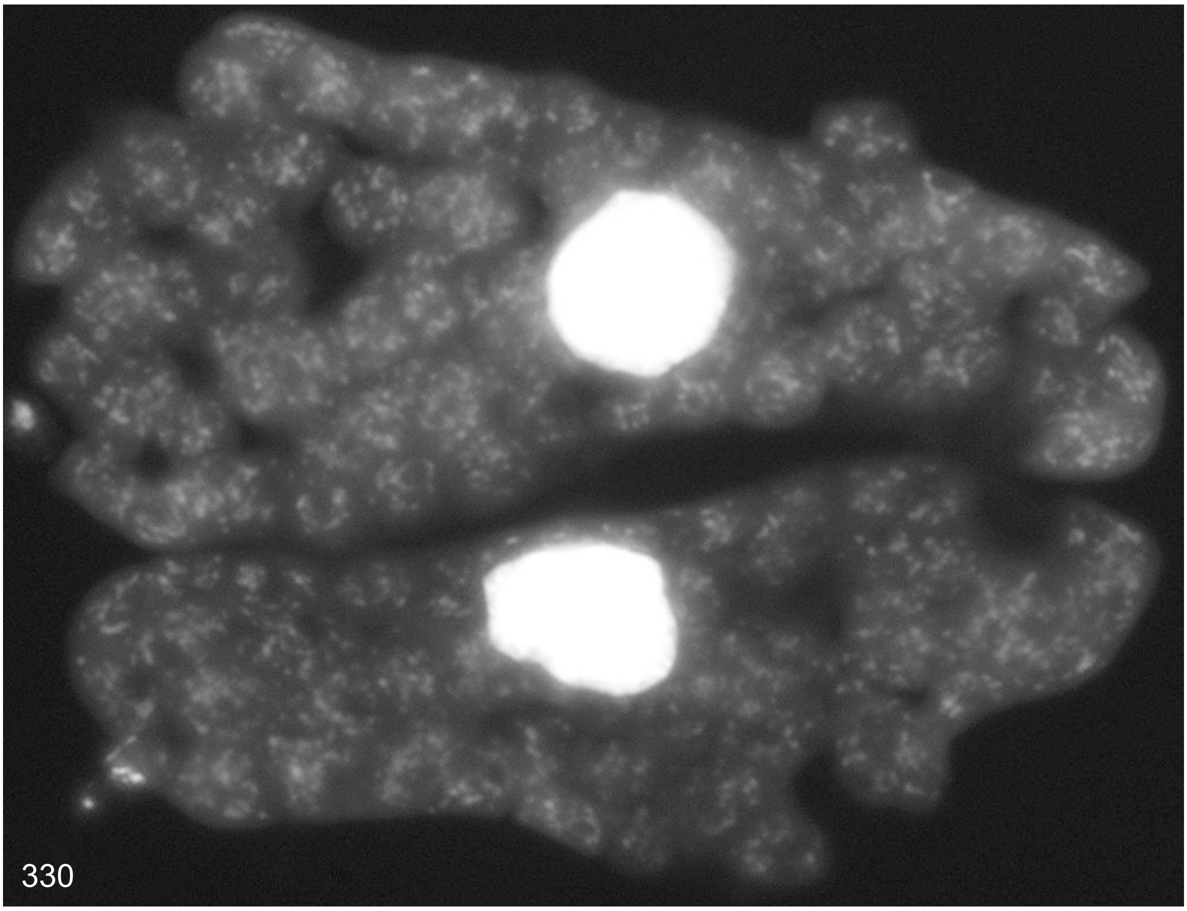

Supplemental Dataset 4

### Maize (*Zea mays*)

Examples of DAPI stained cells around the yellowish meristematic vegetation zone

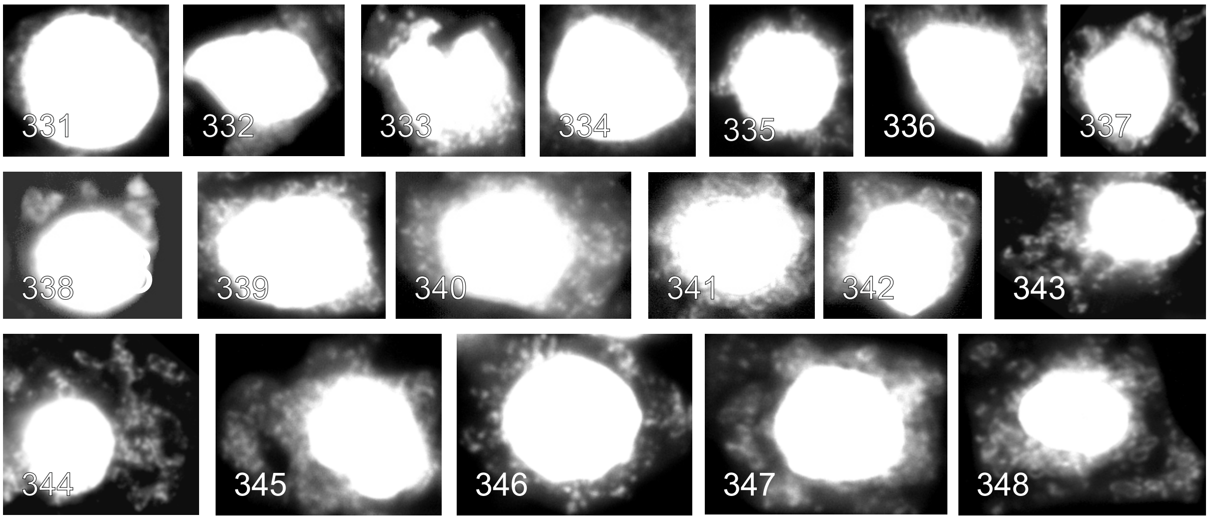

Examples of DAPI stained cells of the greenish region close to the meristematic zone

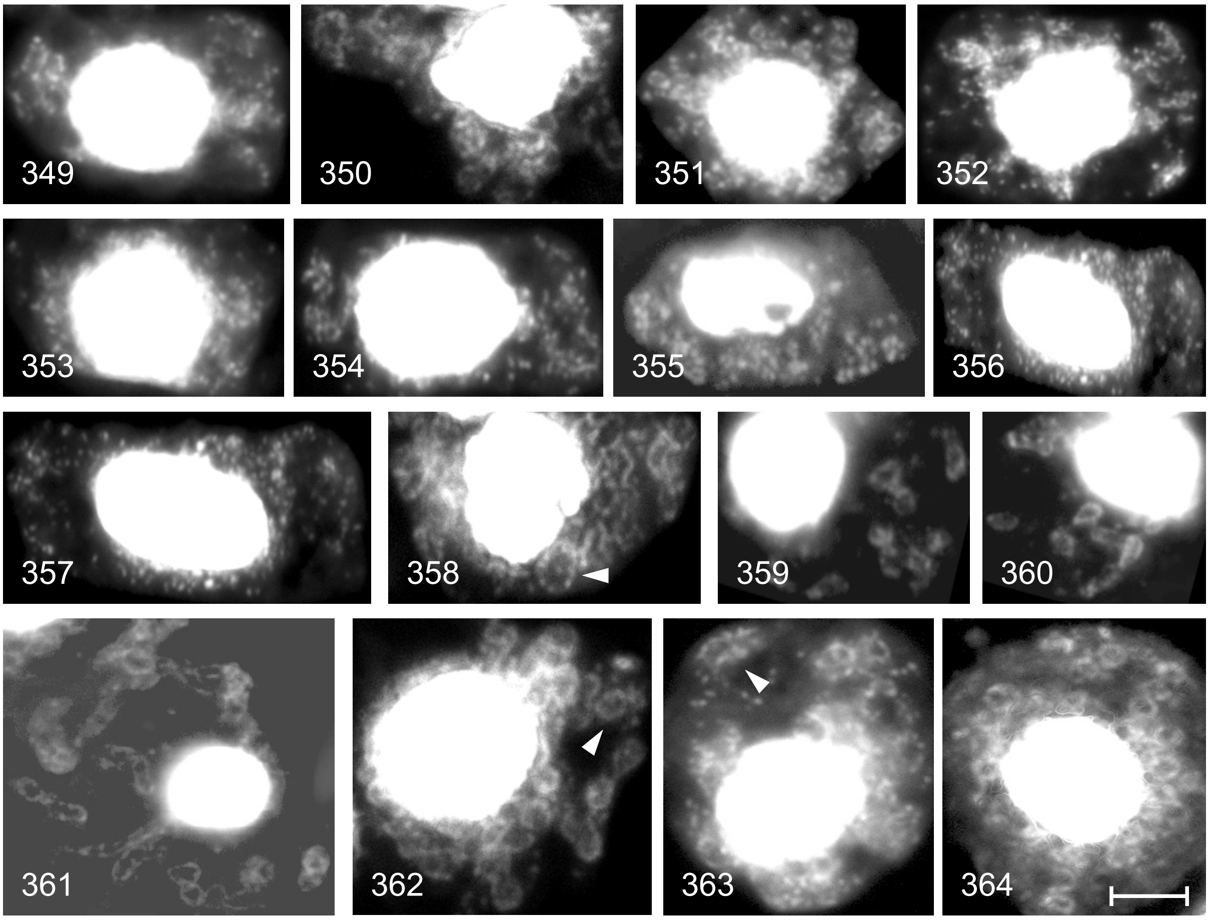

Examples of DAPI stained cell of juvenile leaves

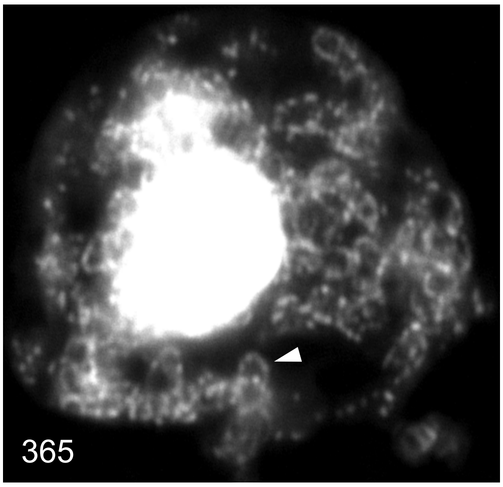
365

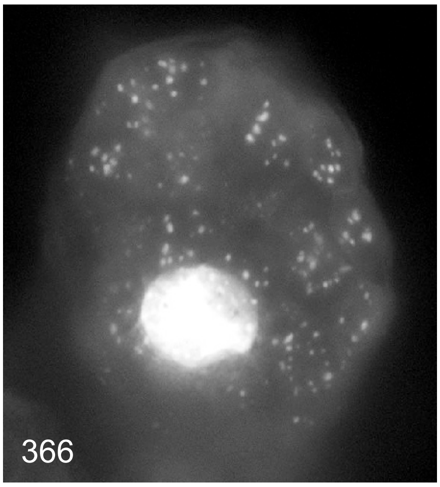
366

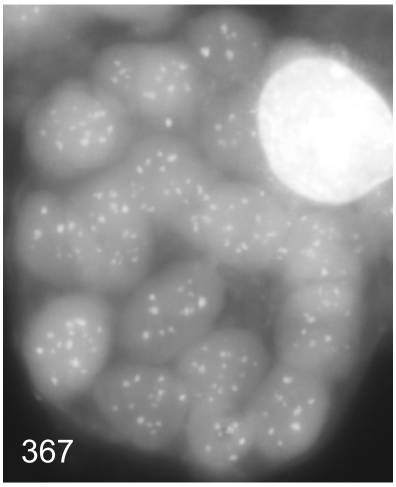
367

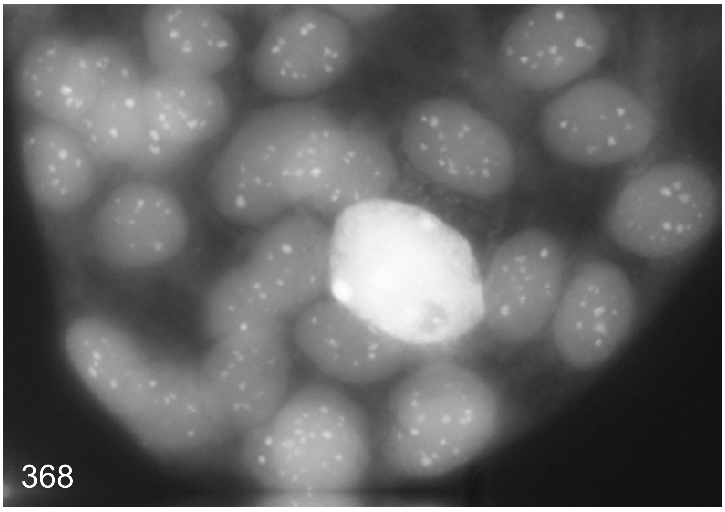
368

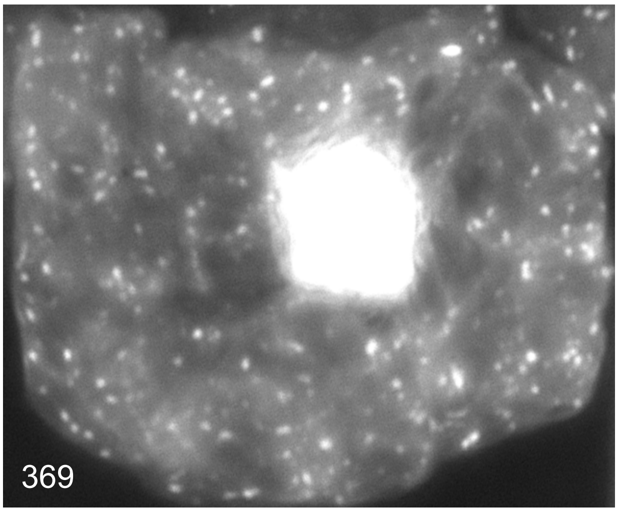
369

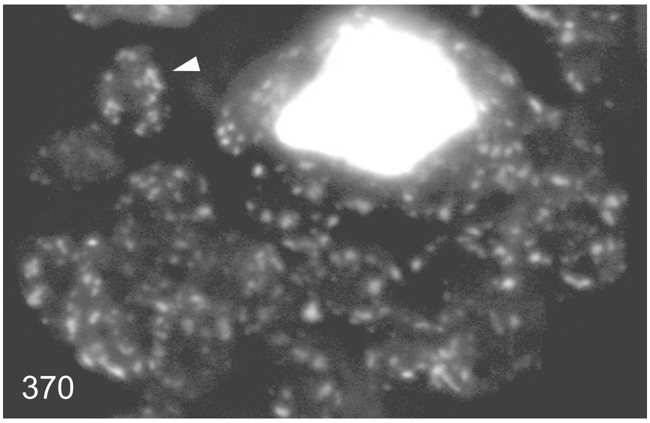
370

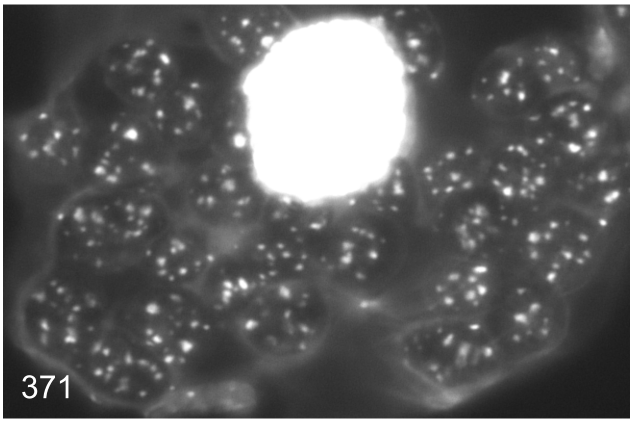
371

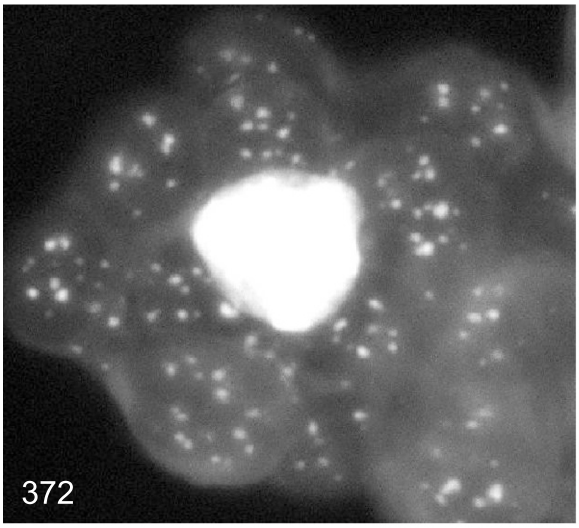
372

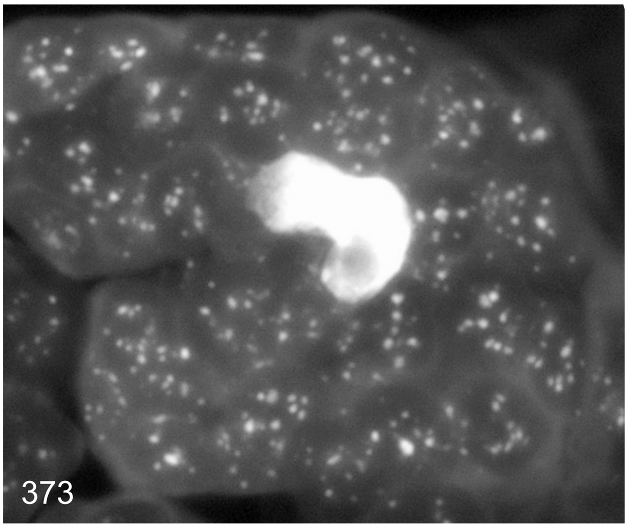
373

Examples of DAPI stained cell of green leaf tissue

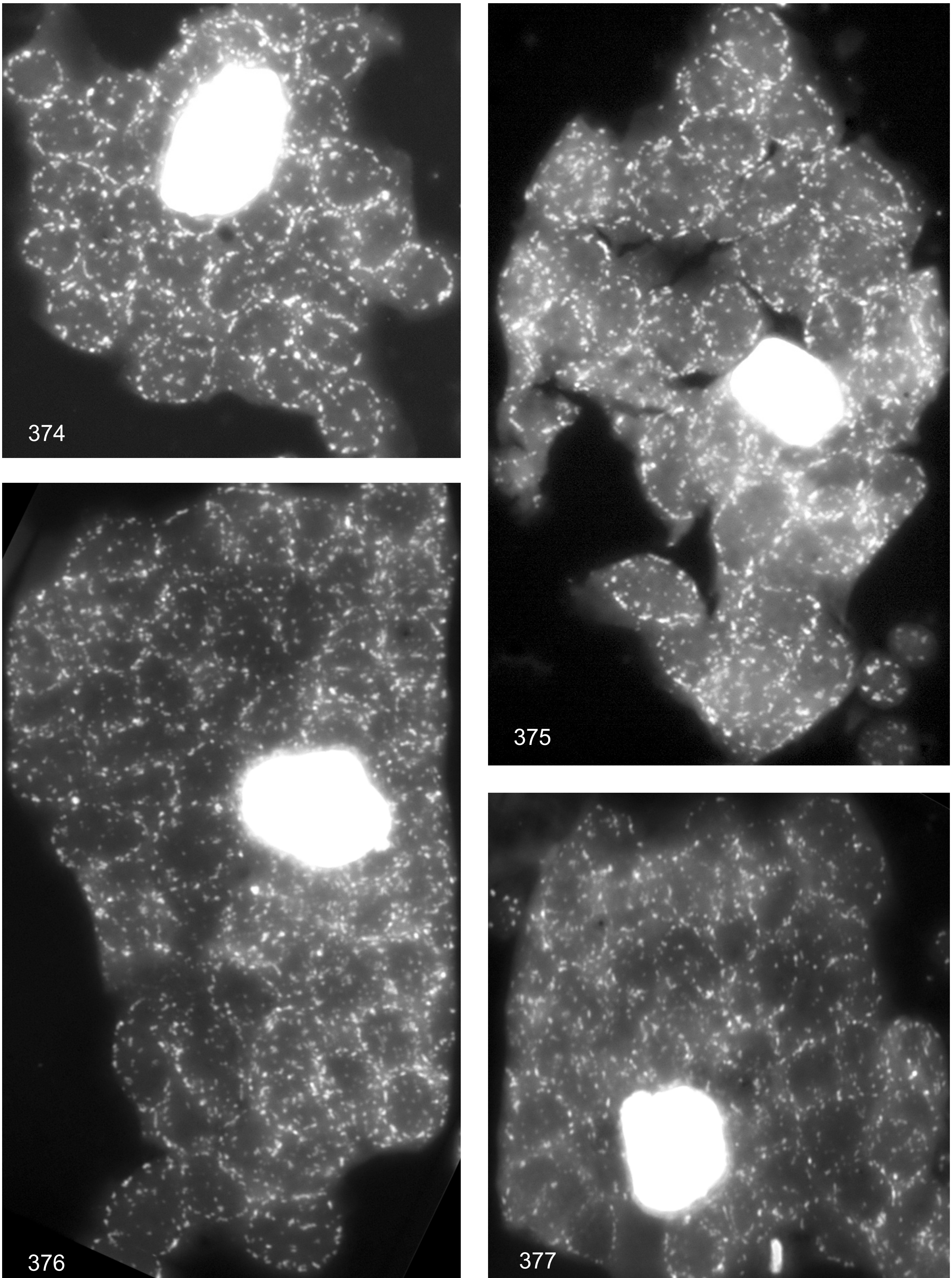

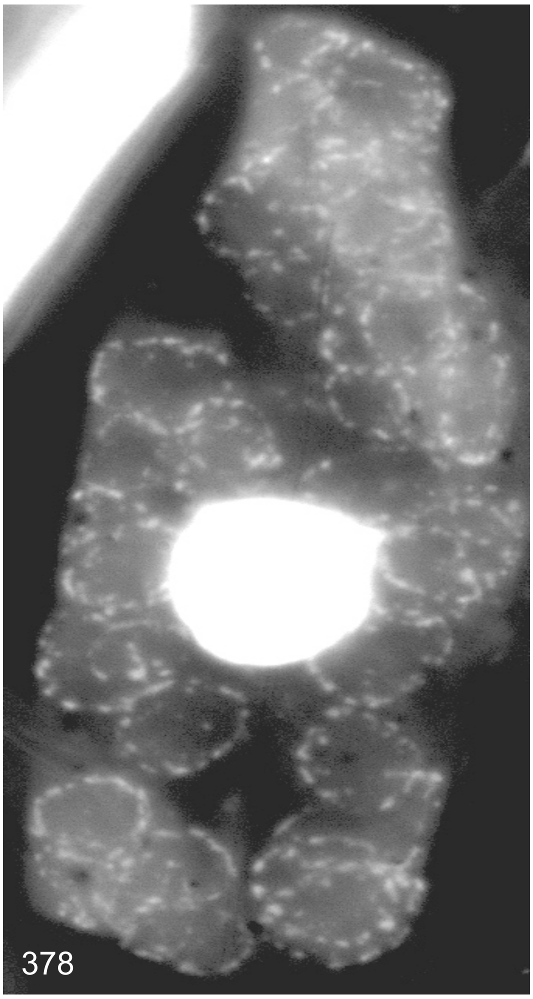
378

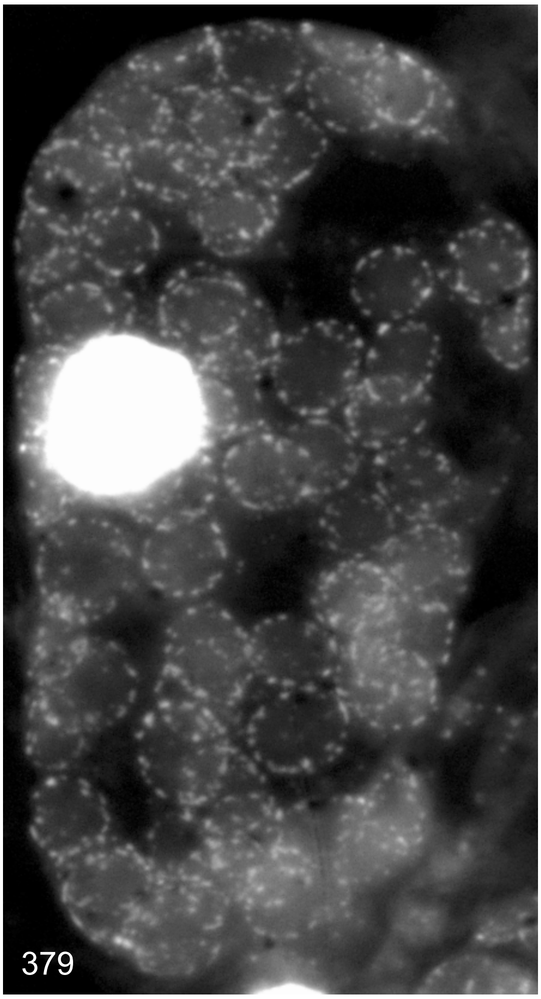
379

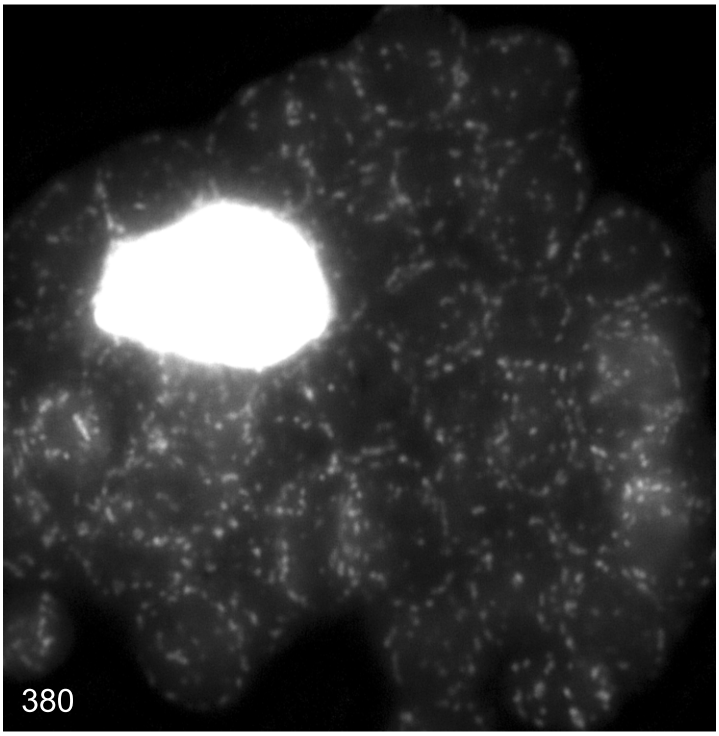
380

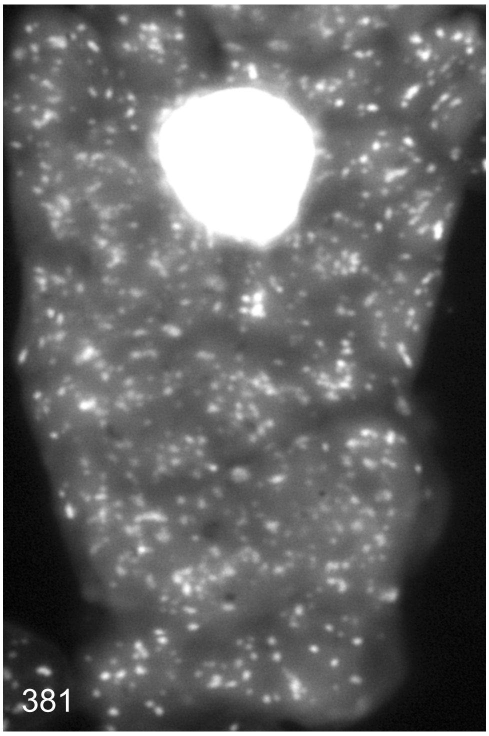
381

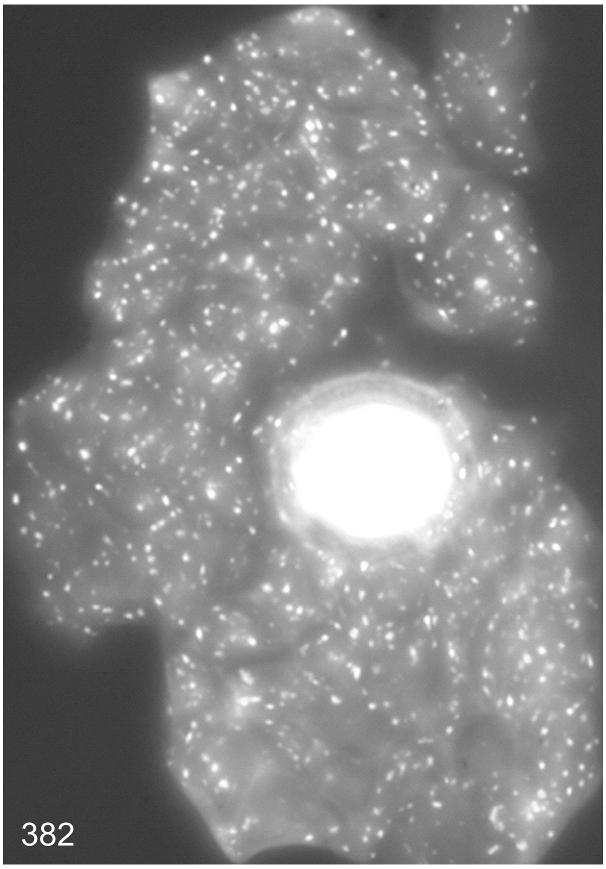
382

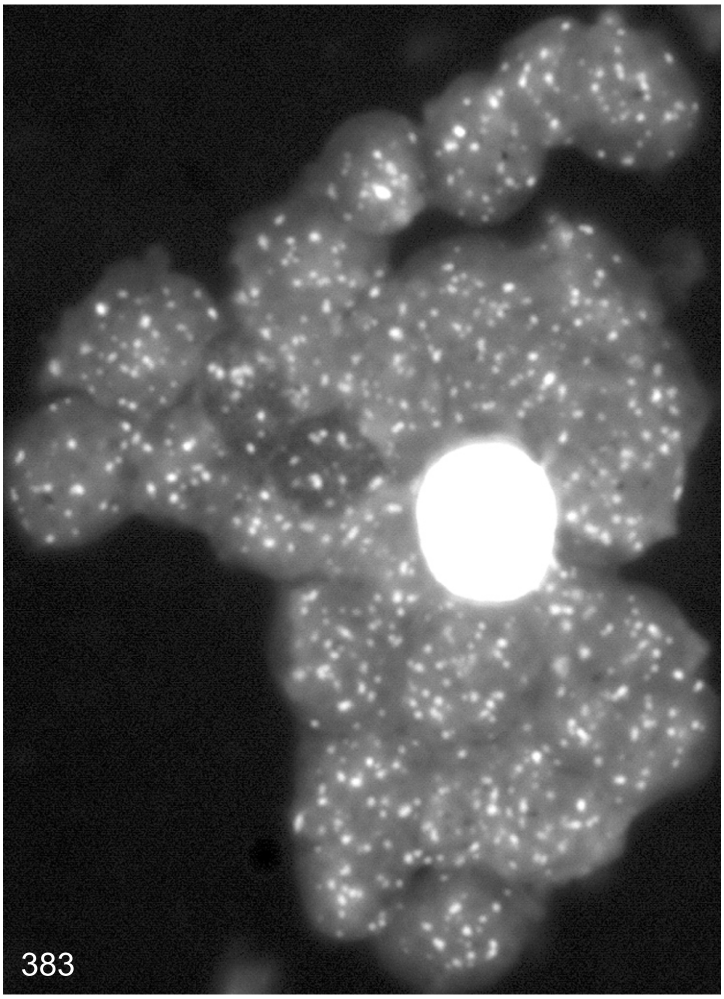
383

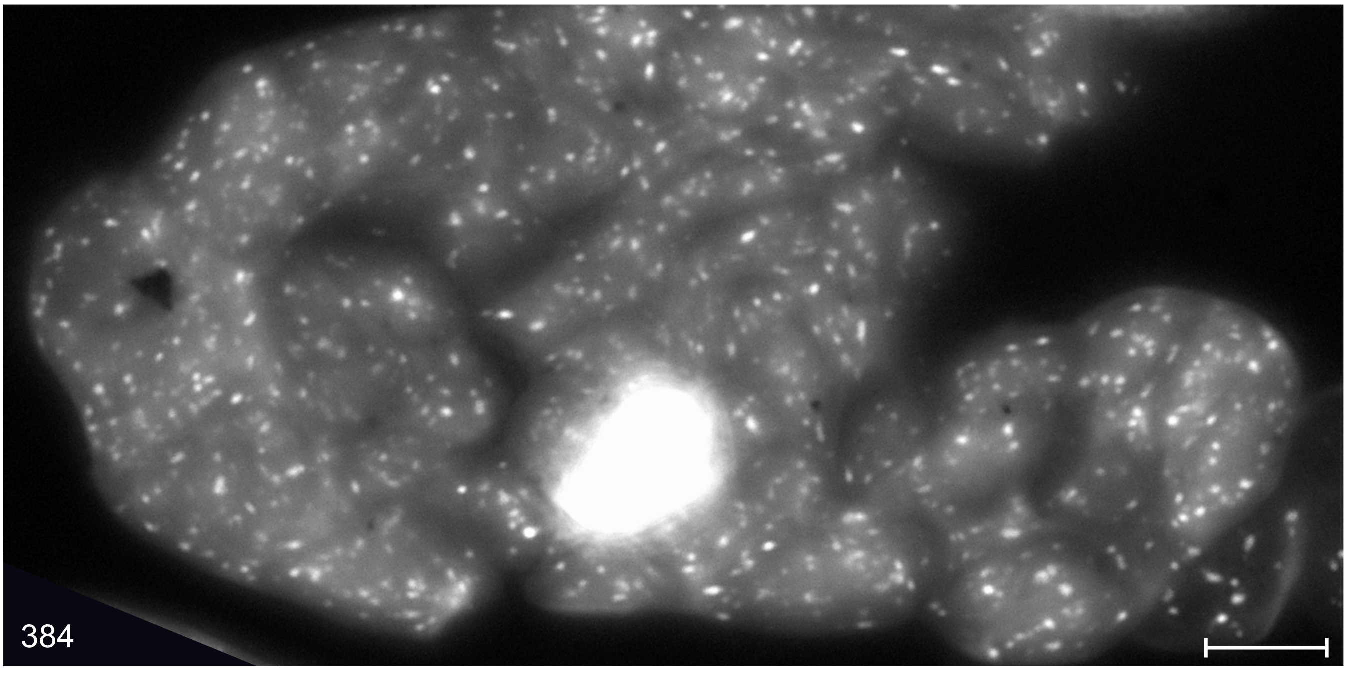
384

**Examples of DAPI stained giant cells**

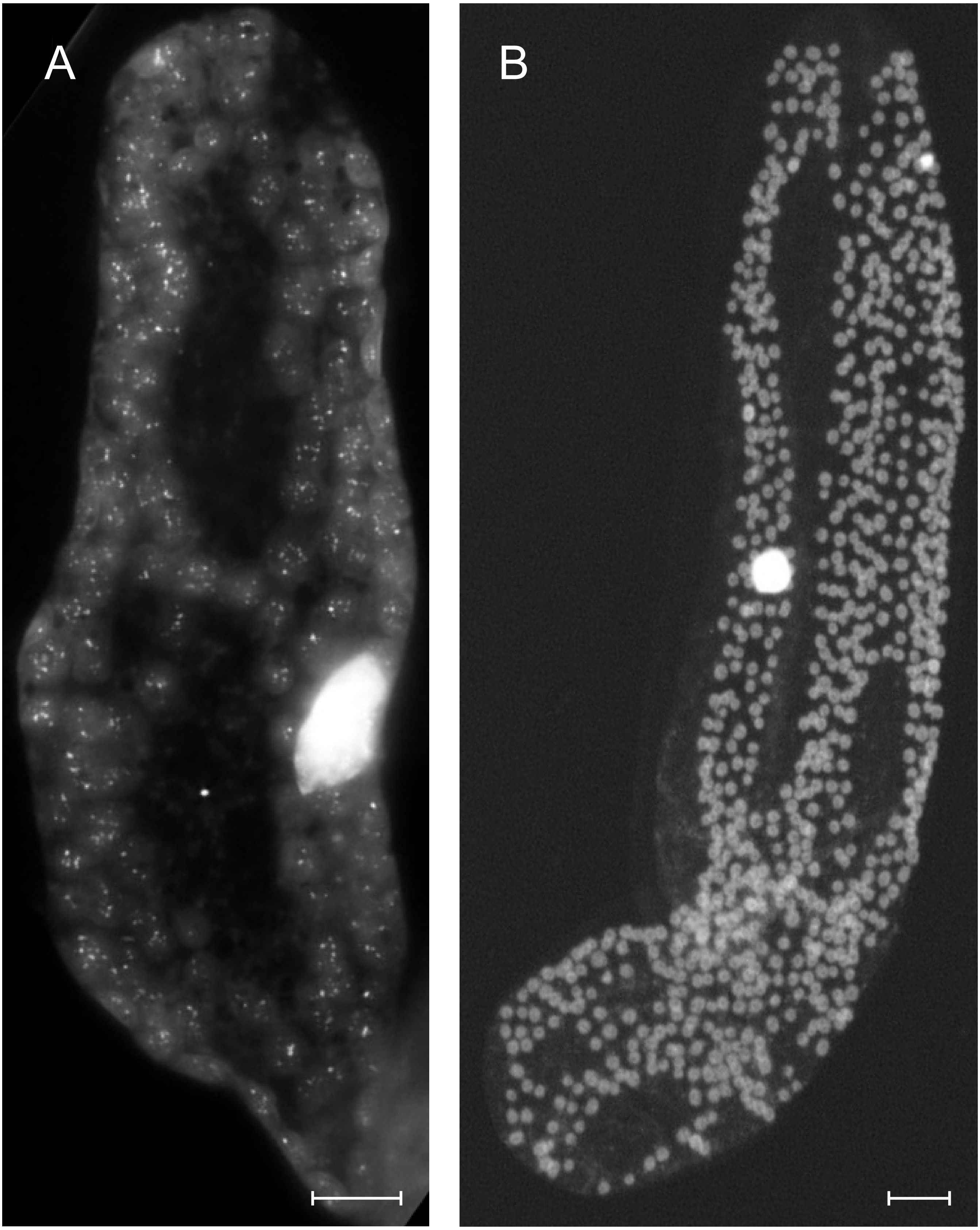

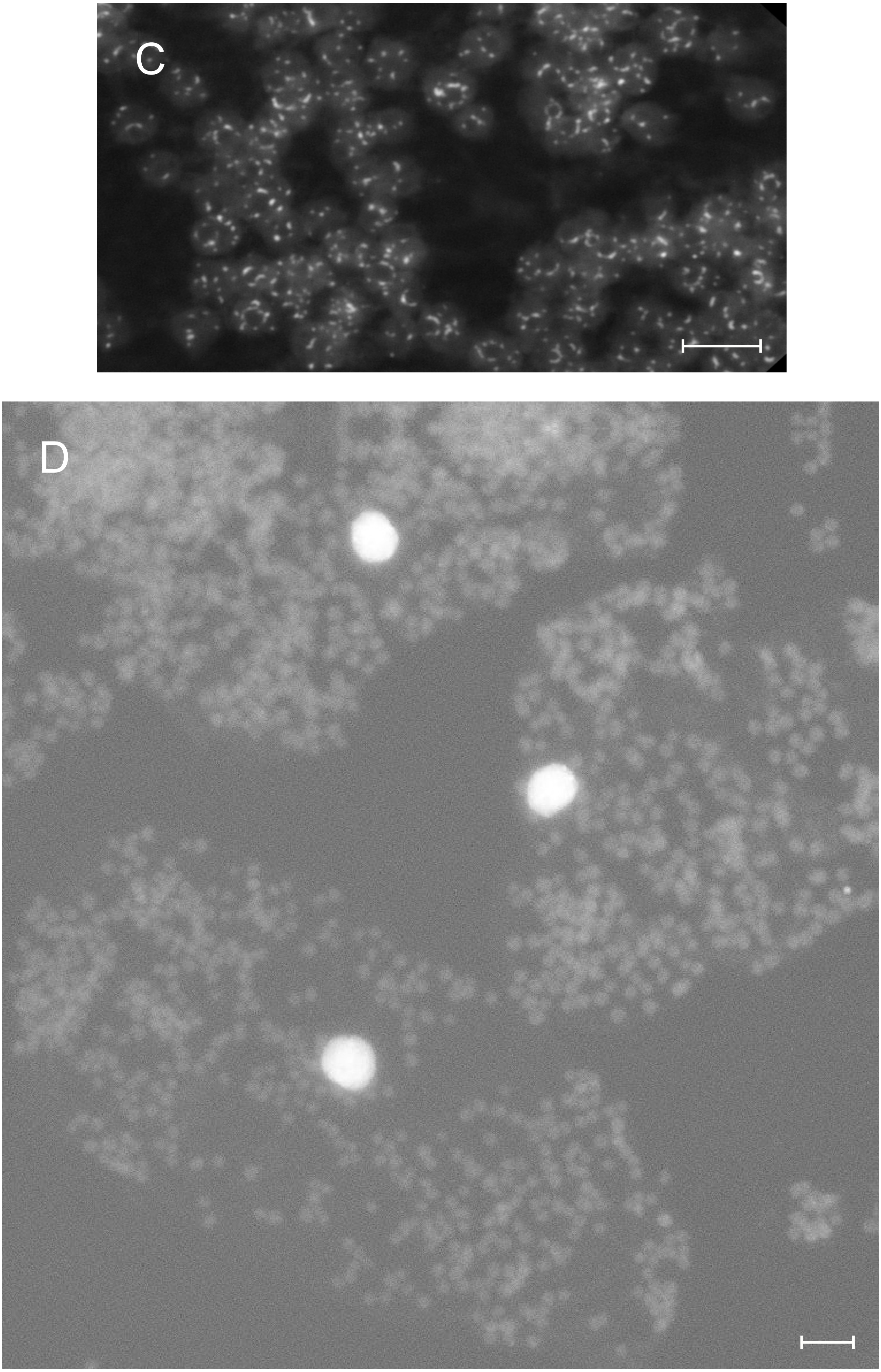
C
D

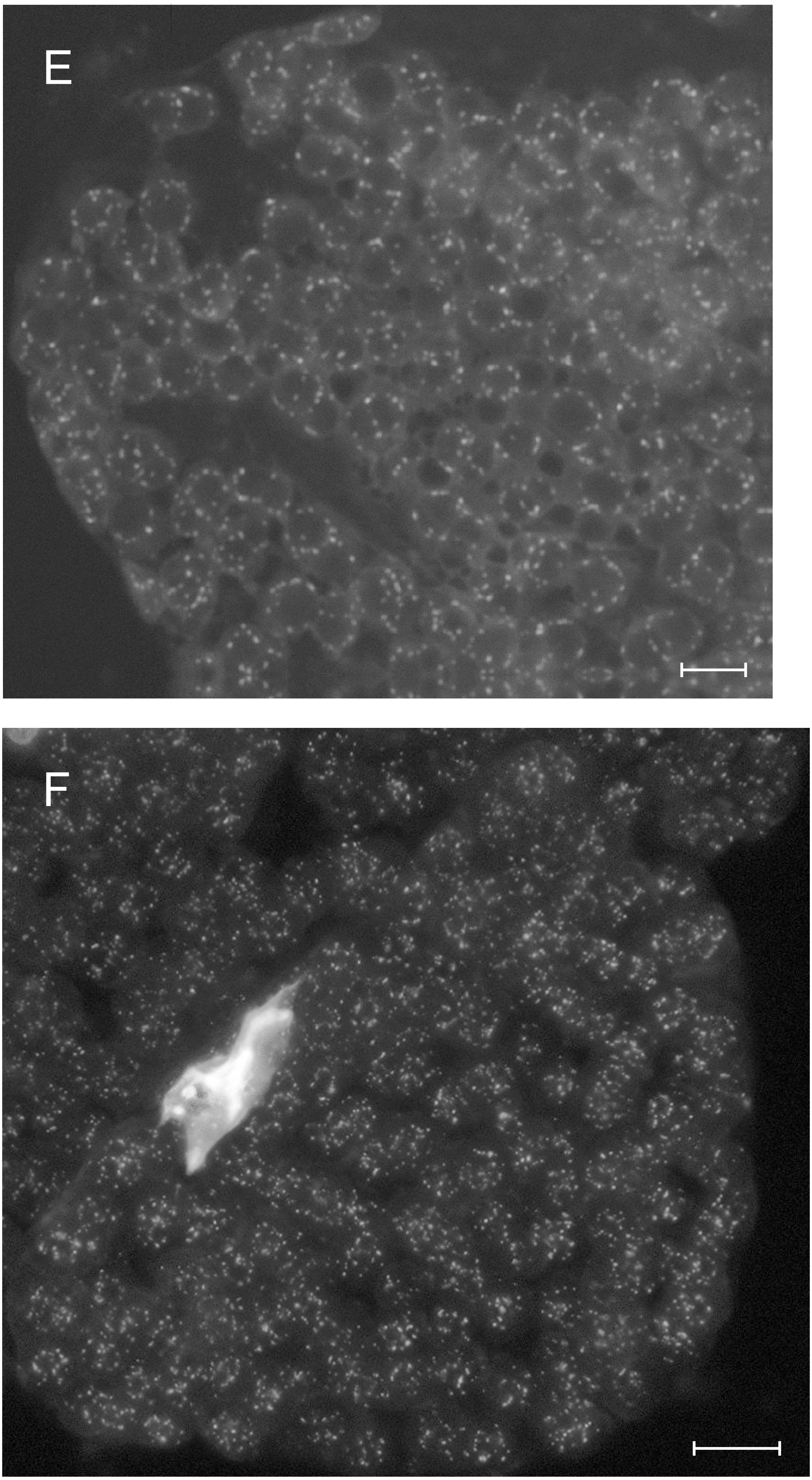
E
F

### Sugar beet (*Beta vulgaris*)

#### Nucleoid densitometry on individual chloroplasts

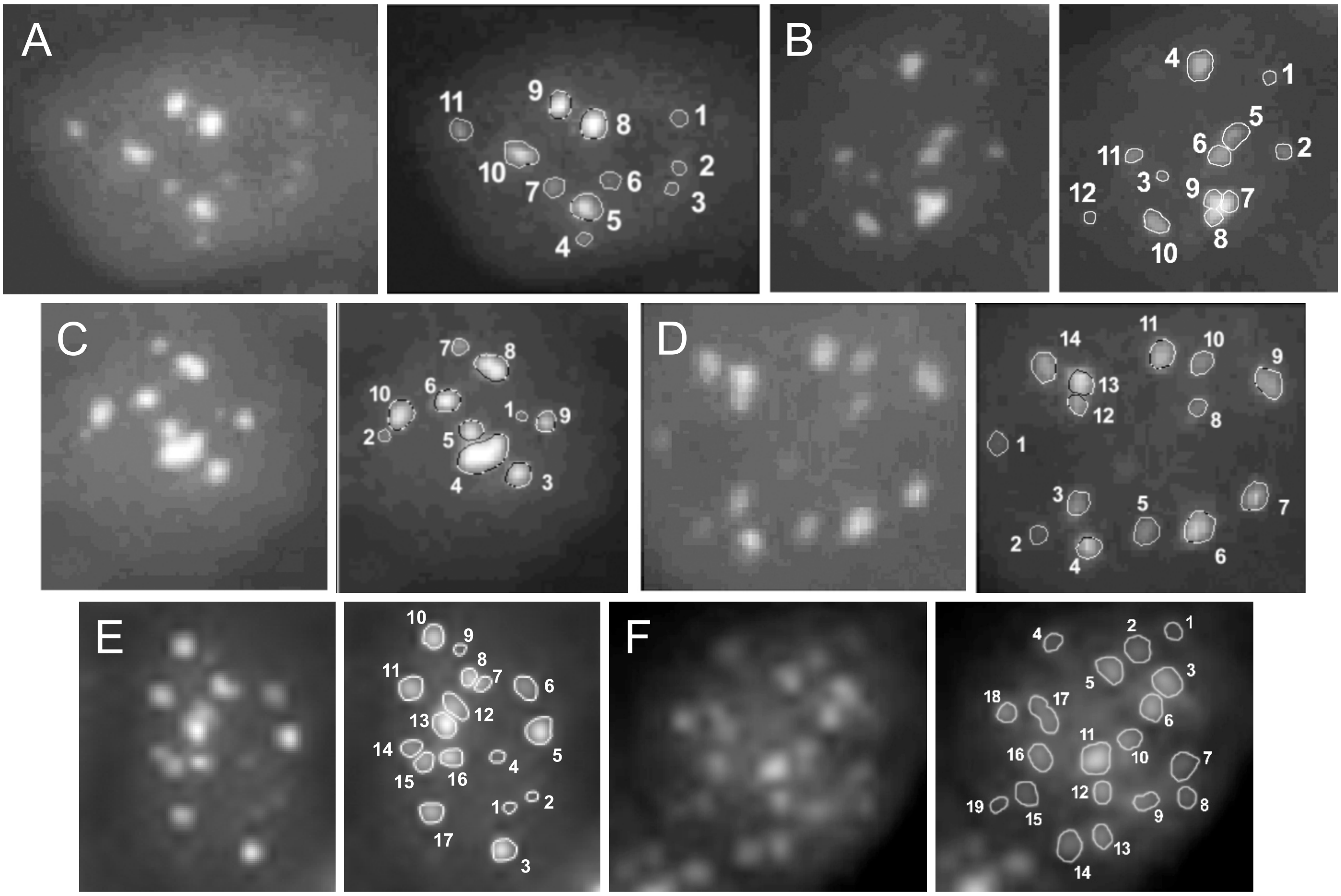

A-C illustrate nucleoid pattern of plastids from juvenile material; D-F those of chloroplasts from premature mesophyll.

T4 normalized measured and visually estimated nucleoid ploidies in individual sugar beet chloroplasts shown in A-F

| Nucleoid no. | Plastid | | | | | |
|---|---|---|---|---|---|---|
| | A | B | C | D | E | F |
| 1 | 1.3 | 0.9 | 0.8 | 1.1 | 1.0 | 1.0 |
| 2 | 1.0 | 1.5 | 1.2 | 0.9 | 0.8 | 2.8 |
| 3 | 0.8 | 0.6 | 4.3 | 2.3 | 6.2 | 5.2 |
| 4 | 1.0 | 5.6 | 20.7 | 3.1 | 1.3 | 2.2 |
| 5 | 6.4 | 3.8 | 4.4 | 2.3 | 7.6 | 4.2 |
| 6 | 1.7 | 3.9 | 5.0 | 4.5 | 4.3 | 4.1 |
| 7 | 2.2 | 5.3 | 2.2 | 3.1 | 2.2 | 2.8 |
| 8 | 8.6 | 2.8 | 9.9 | 1.2 | 3.3 | 1.8 |
| 9 | 5.3 | 2.9 | 3.5 | 3.9 | 0.9 | 2.3 |
| 10 | 6.0 | 3.7 | 8.1 | 2.3 | 5.5 | 2.9 |
| 11 | 2.4 | 1.5 | - | 3.9 | 5.6 | 6.3 |
| 12 | - | 0.7 | - | 1.9 | 5.1 | 3.1 |
| 13 | - | - | - | 4.8 | 7.4 | 2.7 |
| 14 | - | - | - | 3.3 | 3.2 | 3.8 |
| 15 | - | - | - | - | 3.8 | 1.8 |
| 16 | - | - | - | - | 4.6 | 2.9 |
| 17 | - | - | - | - | 4.9 | 3.7 |
| 18 | - | - | - | - | - | 2.3 |
| 19 | - | - | - | - | - | 0.9 |
| Sum measured | 36.7 | 33.2 | 60.1 | 38.6 | 67.7 | 56.8 |
| Sum visually estimated | 35 | 29-33 | - | 34.5-36 | 62-71 | - |
| Average ploidy measured | 3.3 | 3.0 | 6.0 | 2.8 | 4.0 | 3.0 |
| Plastid diameter [µm] | 3.0 | 3.0 | 2.5 | 5.5 | 4.5 | 6.0 |

### Tobacco (*Nicotiana tabacum*)

#### Nucleoid densitometry on individual chloroplasts

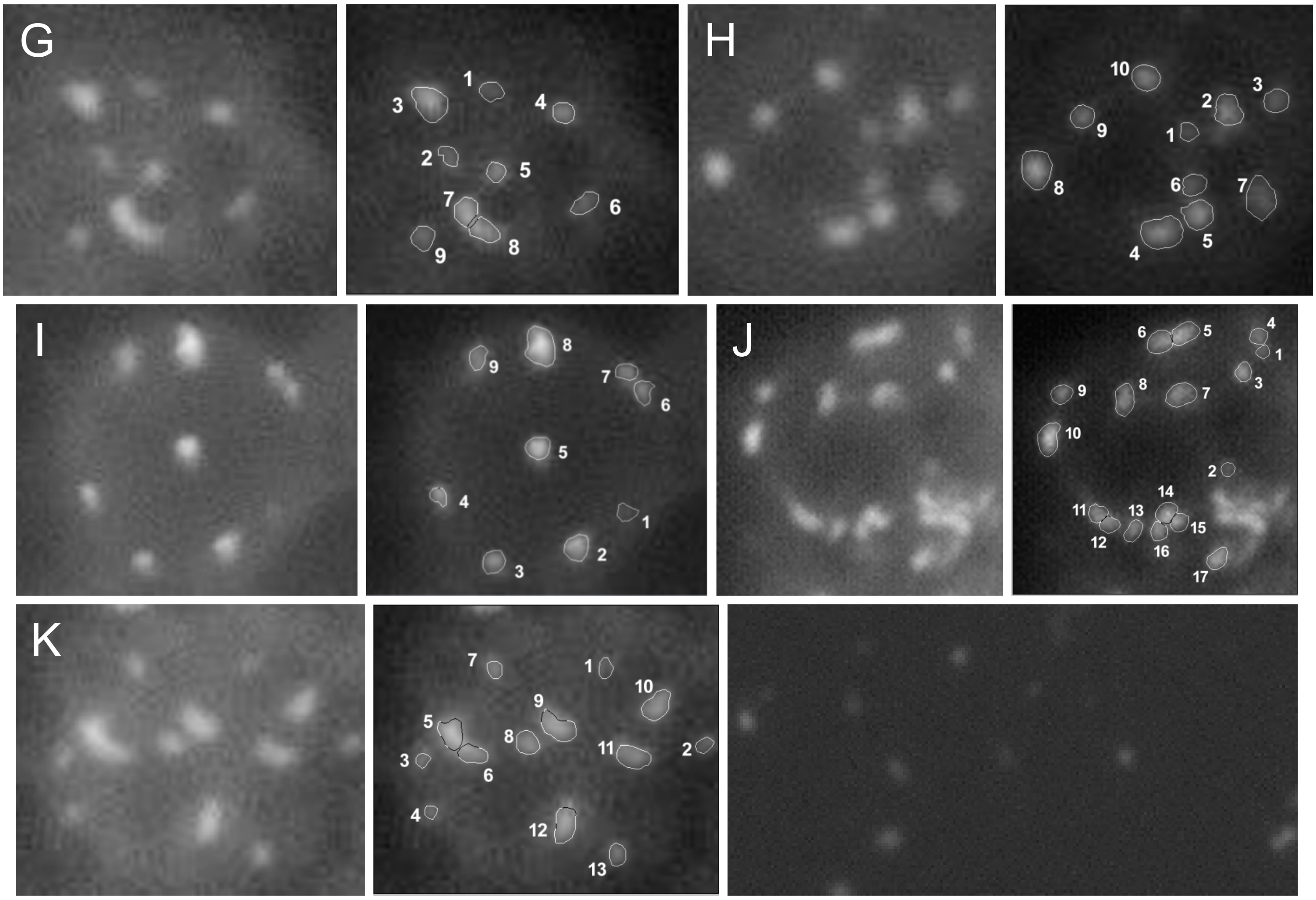

G, H and K are plastids from juvenile mesophyll cells, I and J are examples for chloroplasts of premature mesophyll. T4 standard (third row, right)

T4 normalized measured and visually estimated nucleoid ploidies in individual tobacco chloroplasts shown in G-K

| Nucleoid no. | Plastid | | | | |
|---|---|---|---|---|---|
| | G | H | I | J | K |
| 1 | 0.9 | 1.0 | 0.9 | 1.0 | 1.2 |
| 2 | 1.1 | 3.5 | 4.2 | 1.0 | 1.1 |
| 3 | 5.5 | 1.9 | 3.7 | 2.3 | 0.9 |
| 4 | 2.9 | 5.4 | 2.5 | 1.9 | 0.8 |
| 5 | 2.7 | 3.7 | 4.5 | 4.6 | 5.7 |
| 6 | 1.8 | 1.7 | 2.1 | 4.0 | 4.5 |
| 7 | 4.9 | 3.8 | 2.2 | 4.7 | 1.4 |
| 8 | 4.5 | 4.9 | 8.9 | 3.9 | 2.9 |
| 9 | 1.5 | 2.8 | 2.4 | 2.3 | 5.6 |
| 10 | - | 3.2 | - | 5.3 | 3.7 |
| 11 | - | - | - | 2.2 | 3.6 |
| 12 | - | - | - | 2.3 | 5.1 |
| 13 | - | - | - | 2.2 | 1.7 |
| 14 | - | - | - | 3.3 | - |
| 15 | - | - | - | 2.4 | - |
| 16 | - | - | - | 2.3 | - |
| 17 | - | - | - | 3.3 | - |
| Sum measured | 25.8 | 31.9 | 31.4 | 49.0 | 38.2 |
| Sum visually estimated | - | 29-34 | 29-30 | - | 39-42 |
| Average ploidy measured | 2.9 | 3.2 | 3.5 | 2.9 | 2.9 |
| Plastid diameter [µm] | 2.5 | 3.0 | 4.5 | 5.5 | 3.5 |

### *Arabidopsis thaliana*

Nucleoid densitometry on individual chloroplasts

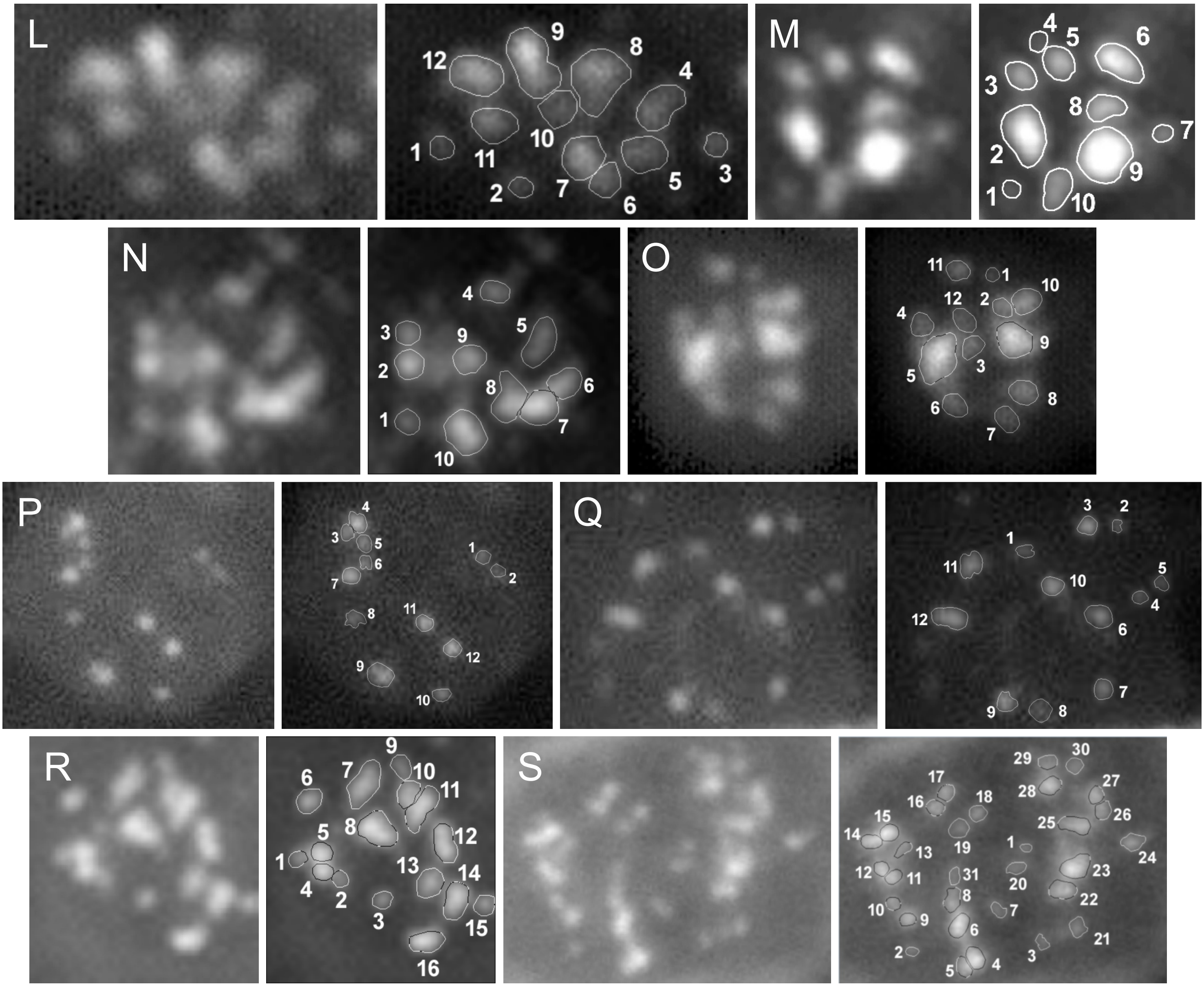

L-O illustrate nucleoid pattern of plastids from juvenile mesophyll, P-R those of chloroplasts from premature and S from mature mesophyll.

### *Arabidopsis thaliana*

#### Nucleoid densitometry on individual chloroplasts

T4 normalized measured and visually estimated nucleoid ploidies in individual *Arabidopsis* chloroplasts shown in L-S

| Nucleoid no. | Plastid | | | | | | | |
|---|---|---|---|---|---|---|---|---|
| | L | M | N | O | P | Q | R | S |
| 1 | 1.0 | 1.0 | 1.0 | 1.0 | 1.1 | 0.9 | 1.0 | 0.8 |
| 2 | 0.9 | 10.8 | 3.8 | 3.4 | 0.9 | 1.1 | 0.9 | 1.0 |
| 3 | 1.3 | 3.5 | 1.6 | 3.5 | 1.4 | 2.7 | 1.2 | 1.3 |
| 4 | 4.6 | 0.9 | 1.3 | 3.7 | 3.4 | 1.5 | 2.7 | 5.5 |
| 5 | 3.8 | 4.0 | 2.4 | 24.9 | 1.7 | 2.0 | 3.4 | 3.4 |
| 6 | 2.3 | 8.9 | 2.6 | 3.7 | 1.4 | 3.0 | 2.3 | 6.3 |
| 7 | 5.5 | 2.3 | 5.9 | 3.3 | 3.9 | 3.1 | 4.9 | 1.6 |
| 8 | 9.7 | 4.8 | 3.6 | 5.8 | 1.5 | 2.3 | 8.5 | 4.2 |
| 9 | 11.2 | 12.4 | 3.6 | 18.0 | 4.2 | 3.9 | 1.4 | 3.5 |
| 10 | 2.4 | 3.8 | 6.9 | 7.3 | 1.7 | 3.9 | 2.5 | 2.0 |
| 11 | 3.8 | - | - | 3.2 | 3.7 | 4.2 | 5.0 | 2.8 |
| 12 | 7.2 | - | - | 2.7 | 3.9 | 5.9 | 4.5 | 2.8 |
| 13 | - | - | - | - | - | - | 3.2 | 2.0 |
| 14 | - | - | - | - | - | - | 4.6 | 3.7 |
| 15 | - | - | - | - | - | - | 1.6 | 4.8 |
| 16 | - | - | - | - | - | - | 4.1 | 3.0 |
| 17 | - | - | - | - | - | - | - | 2.8 |
| 18 | - | - | - | - | - | - | - | 2.1 |
| 19 | - | - | - | - | - | - | - | 1.9 |
| 20 | - | - | - | - | - | - | - | 2.0 |
| 21 | - | - | - | - | - | - | - | 2.3 |
| 22 | - | - | - | - | - | - | - | 6.0 |
| 23 | - | - | - | - | - | - | - | 9.2 |
| 24 | - | - | - | - | - | - | - | 3.3 |
| 25 | - | - | - | - | - | - | - | 6.1 |
| 26 | - | - | - | - | - | - | - | 2.8 |
| 27 | - | - | - | - | - | - | - | 3.0 |
| 28 | - | - | - | - | - | - | - | 4.5 |
| 29 | - | - | - | - | - | - | - | 2.6 |
| 30 | - | - | - | - | - | - | - | 2.3 |
| 31 | - | - | - | - | - | - | - | 1.7 |
| Sum measured | 53.7 | 52.4 | 32.7 | 80.5 | 28.8 | 34.5 | 51.8 | 101.3 |
| Sum visually estimated | - | 58.5-60 | - | 64-71 | 26-29 | 36.5-37 | - | - |
| Average ploidy measured | 4.5 | 5.2 | 3.3 | 6.7 | 2.4 | 2.9 | 3.2 | 3.3 |
| Plastid diameter [µm] | 4.0 | 2.5 | 3.5 | 2.5 | 4.0 | 5.0 | 4.0 | 6.5 |

### Maize (*Zea mays*)

#### Nucleoid densitometry on individual chloroplasts

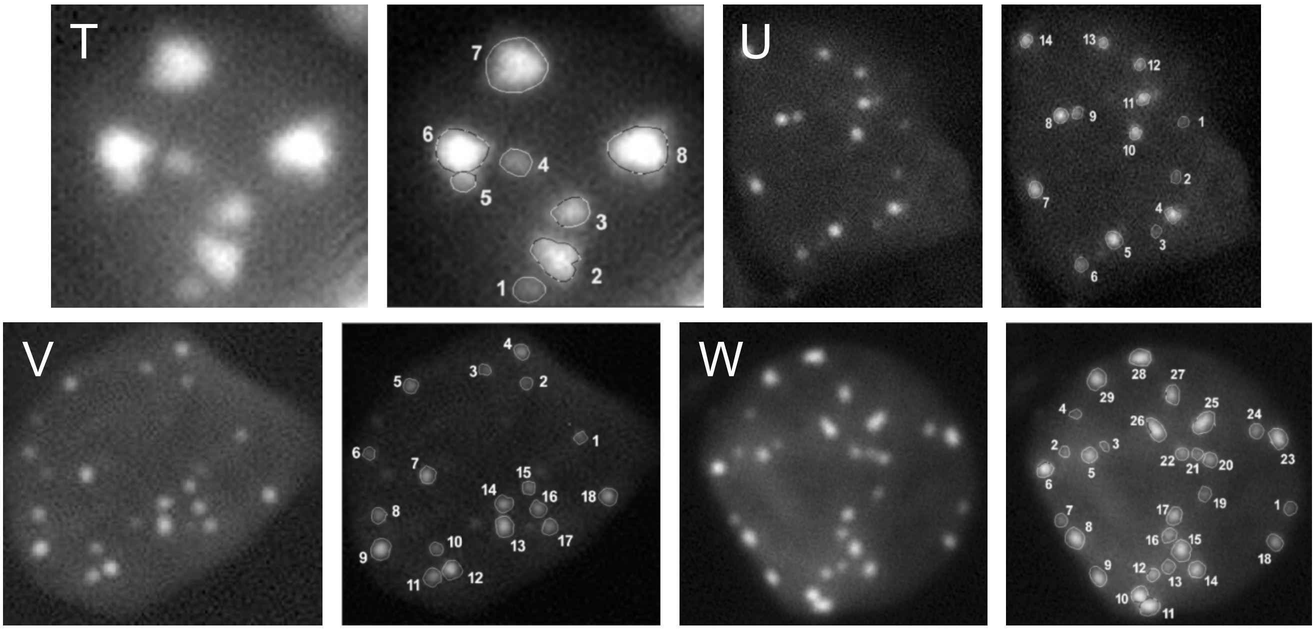

T illustrates the nucleoid  pattern of a plastid of a post-meristematic leaflet cell, U and V are from juvenile material, W is from mature leaf tissue

T4 normalized measured and visually estimated nucleoid ploidies in individual maize chloroplasts shown in T-W

| Nucleoid no. | Plastid | | | |
|---|---|---|---|---|
| | T | U | V | W |
| 1 | 1.0 | 0.9 | 1.0 | 1.2 |
| 2 | 4.8 | 1.1 | 1.0 | 1.1 |
| 3 | 2.1 | 1.0 | 0.9 | 0.8 |
| 4 | 1.1 | 3.0 | 2.5 | 0.9 |
| 5 | 0.8 | 3.7 | 2.0 | 3.1 |
| 6 | 6.7 | 1.7 | 1.0 | 4.0 |
| 7 | 5.7 | 2.9 | 2.5 | 1.5 |
| 8 | 7.6 | 3.0 | 1.8 | 5.0 |
| 9 | - | 1.3 | 3.8 | 3.8 |
| 10 | - | 2.6 | 1.3 | 4.3 |
| 11 | - | 2.9 | 2.5 | 5.2 |
| 12 | - | 1.5 | 3.7 | 2.1 |
| 13 | - | 1.6 | 4.2 | 2.0 |
| 14 | - | 2.5 | 2.7 | 3.6 |
| 15 | - | - | 1.4 | 5.5 |
| 16 | - | - | 2.1 | 2.5 |
| 17 | - | - | 2.0 | 3.3 |
| 18 | - | - | 2.7 | 2.3 |
| 19 | - | - | - | 1.6 |
| 20 | - | - | - | 2.3 |
| 21 | - | - | - | 1.4 |
| 22 | - | - | - | 1.9 |
| 23 | - | - | - | 4.4 |
| 24 | - | - | - | 2.1 |
| 25 | - | - | - | 6.2 |
| 26 | - | - | - | 5.4 |
| 27 | - | - | - | 3.3 |
| 28 | - | - | - | 4.9 |
| 29 | - | - | - | 4.4 |
| Sum measured | 29.8 | 29.7 | 39.1 | 90.1 |
| Sum visually estimated | 32-33 | 30 | 37-38 | - |
| Average ploidy measured | 3.7 | 2.1 | 2.2 | 3.1 |
| Plastid diameter [µm] | 2.5 | 3.5 | 4.5 | 6.0 |

Supplemental Dataset 7

#### Examples of DAPI stained nucleoids in mesophyll chloroplasts

##### Sugar beet (*Beta vulgaris*)

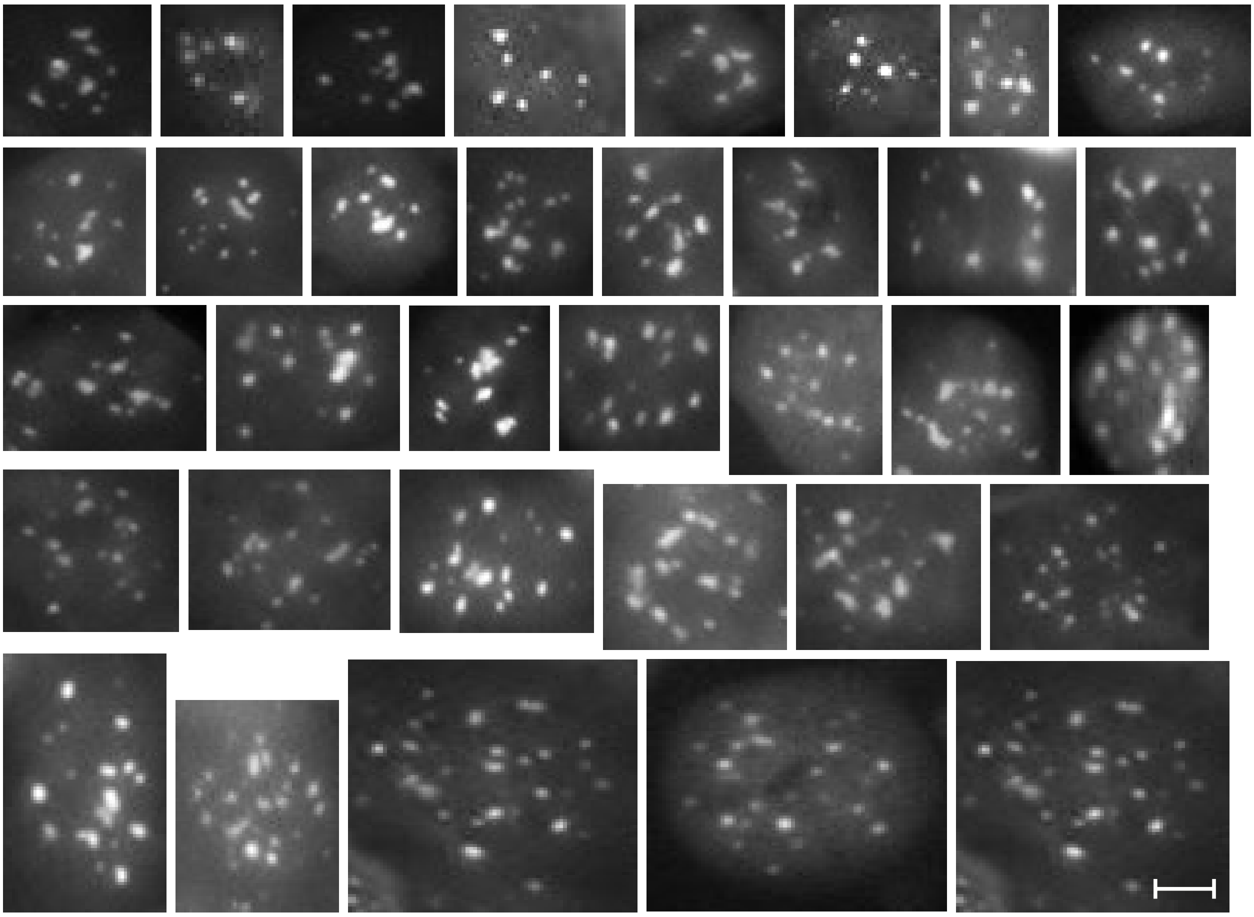

##### *Arabidopsis thaliana*

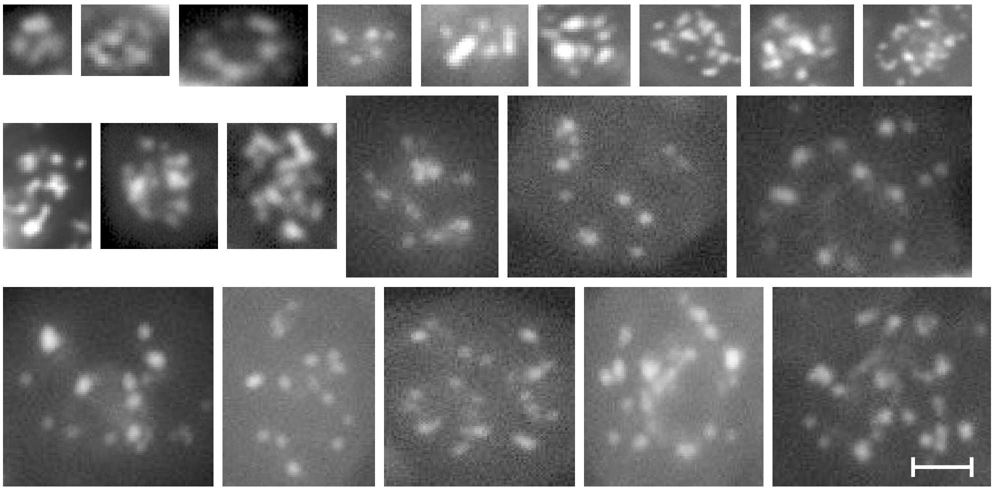

Tabacco (*Nicotiana tabacum*)

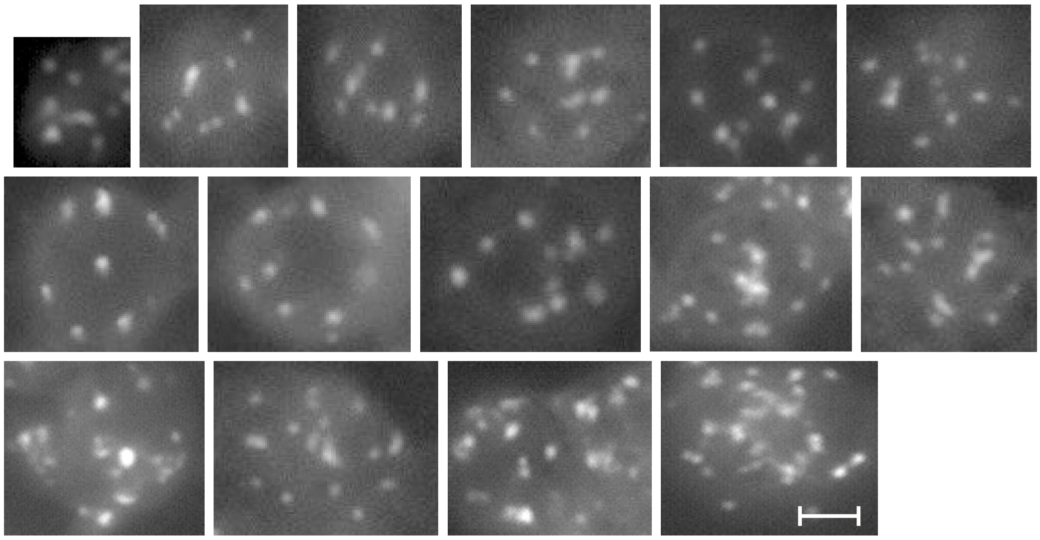

Maize (*Zea mays*)

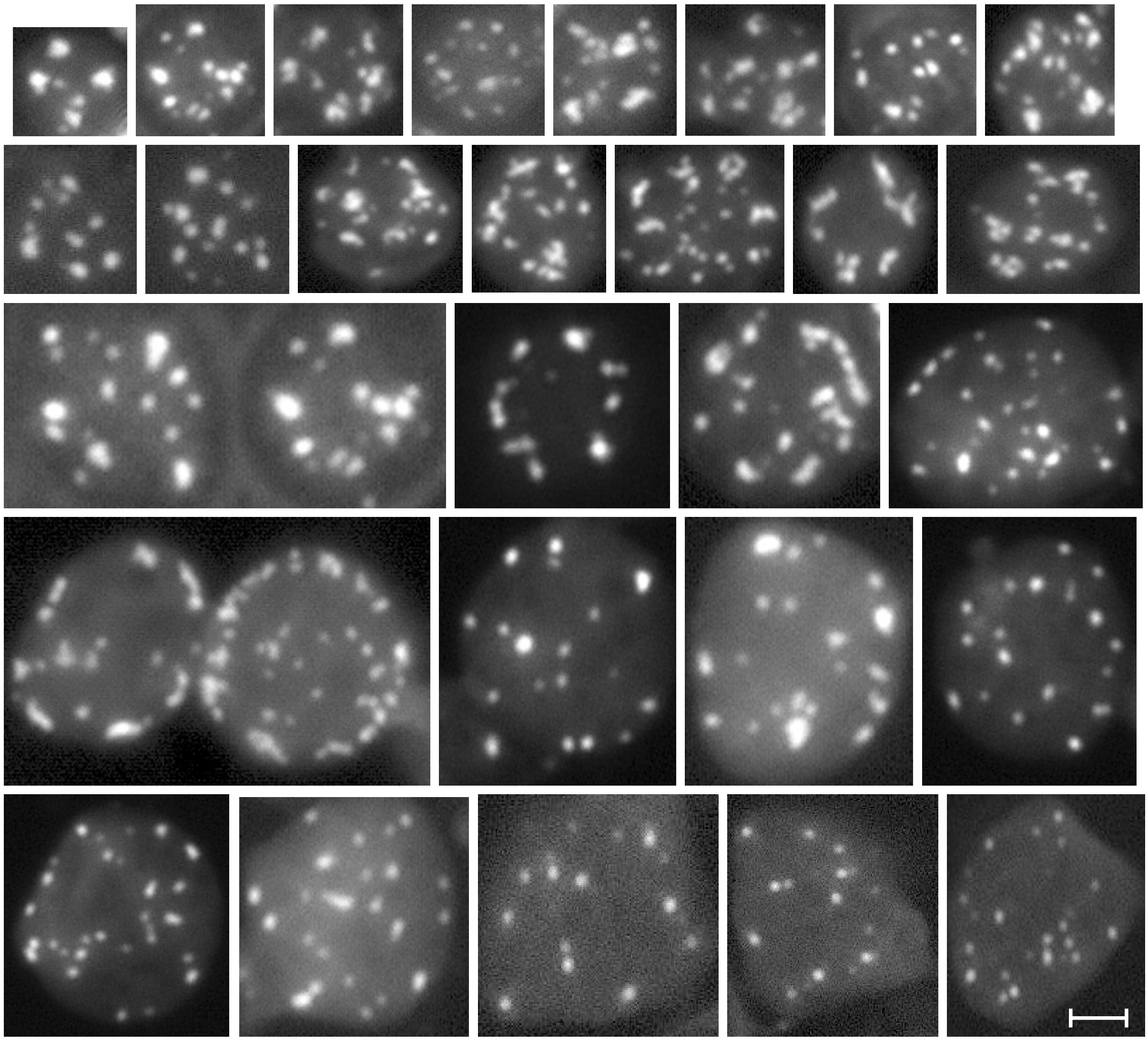

Supplemental Dataset 8

**Examples of protoplast preparations**

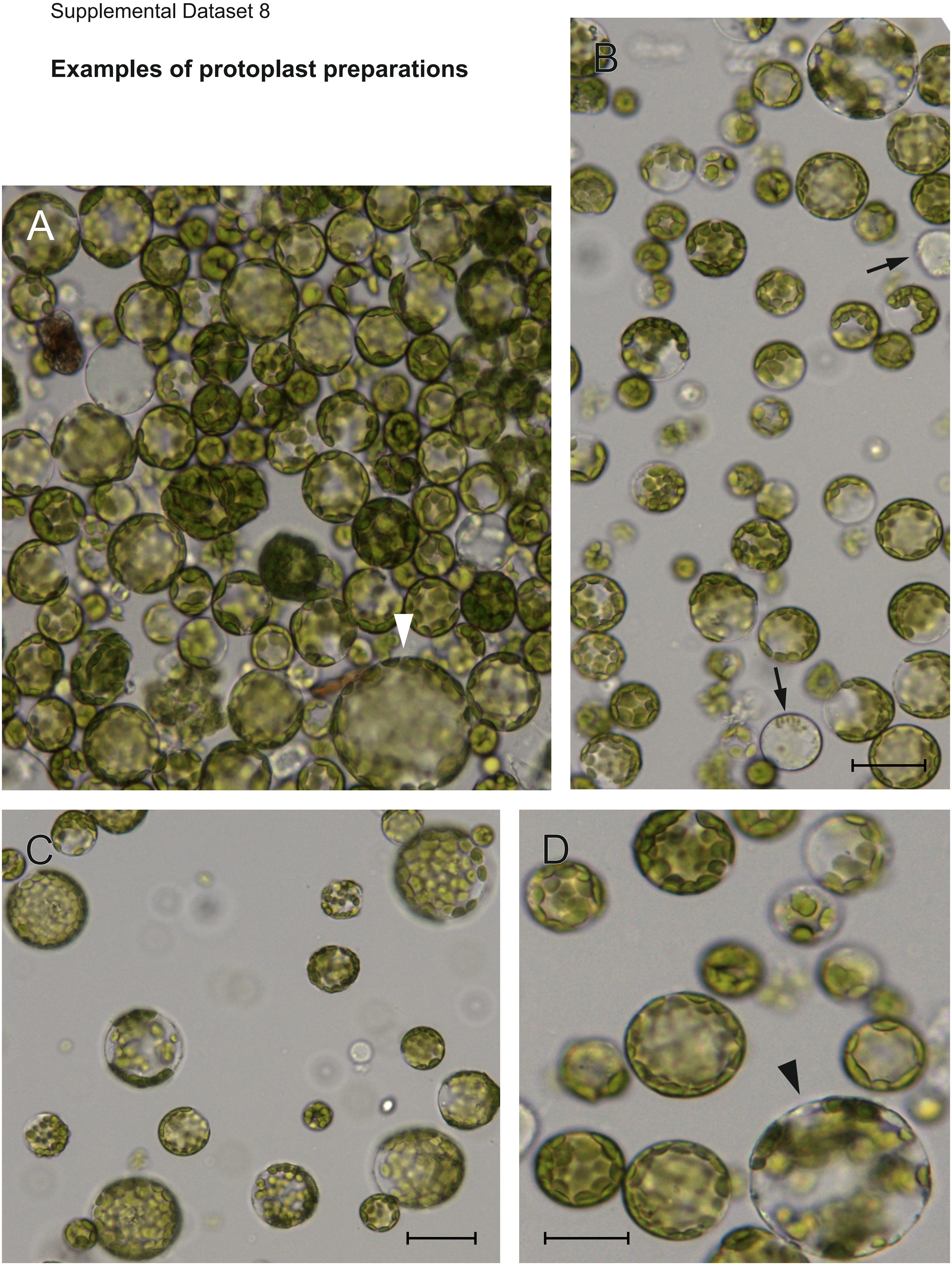

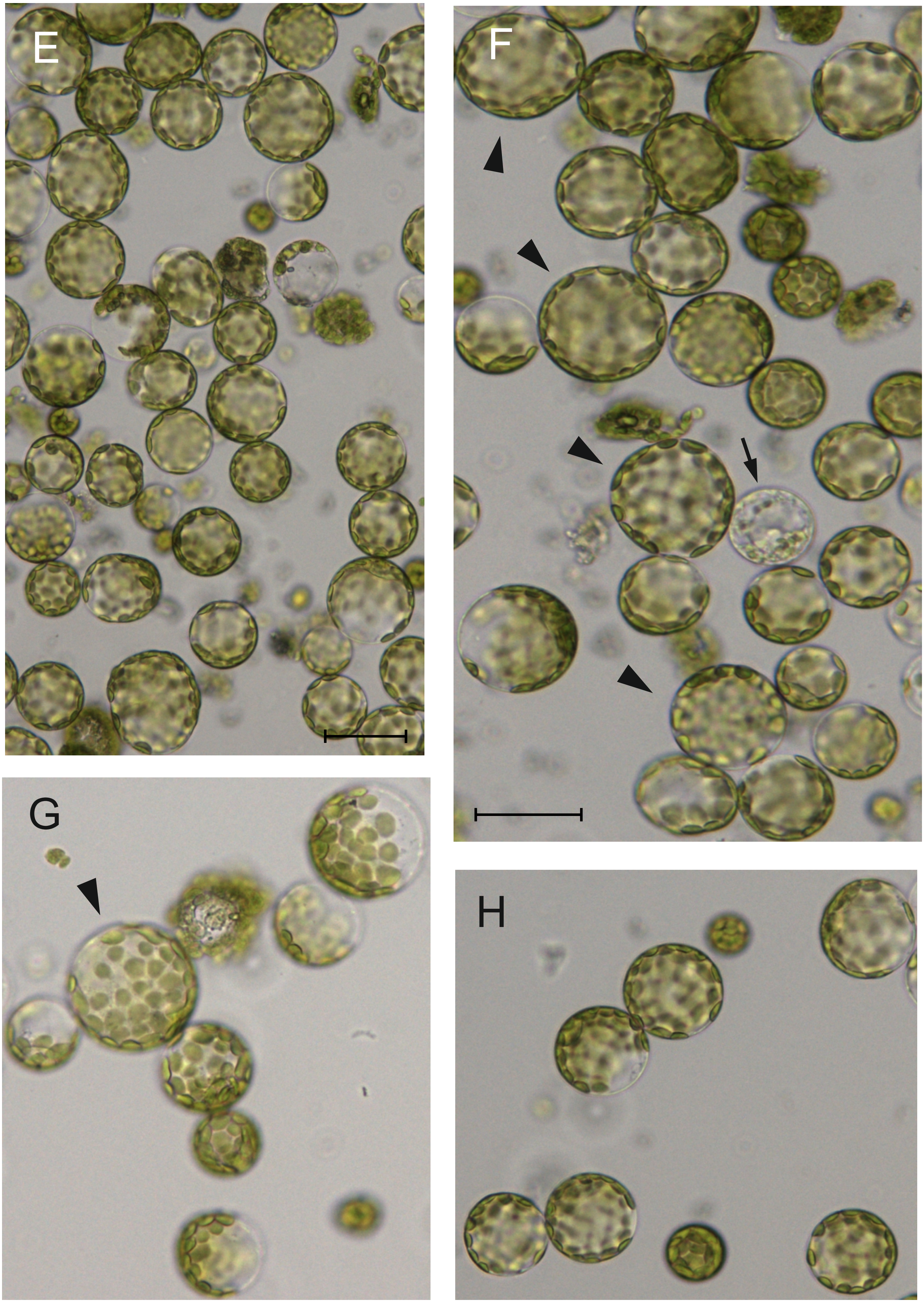
E
F
G
H

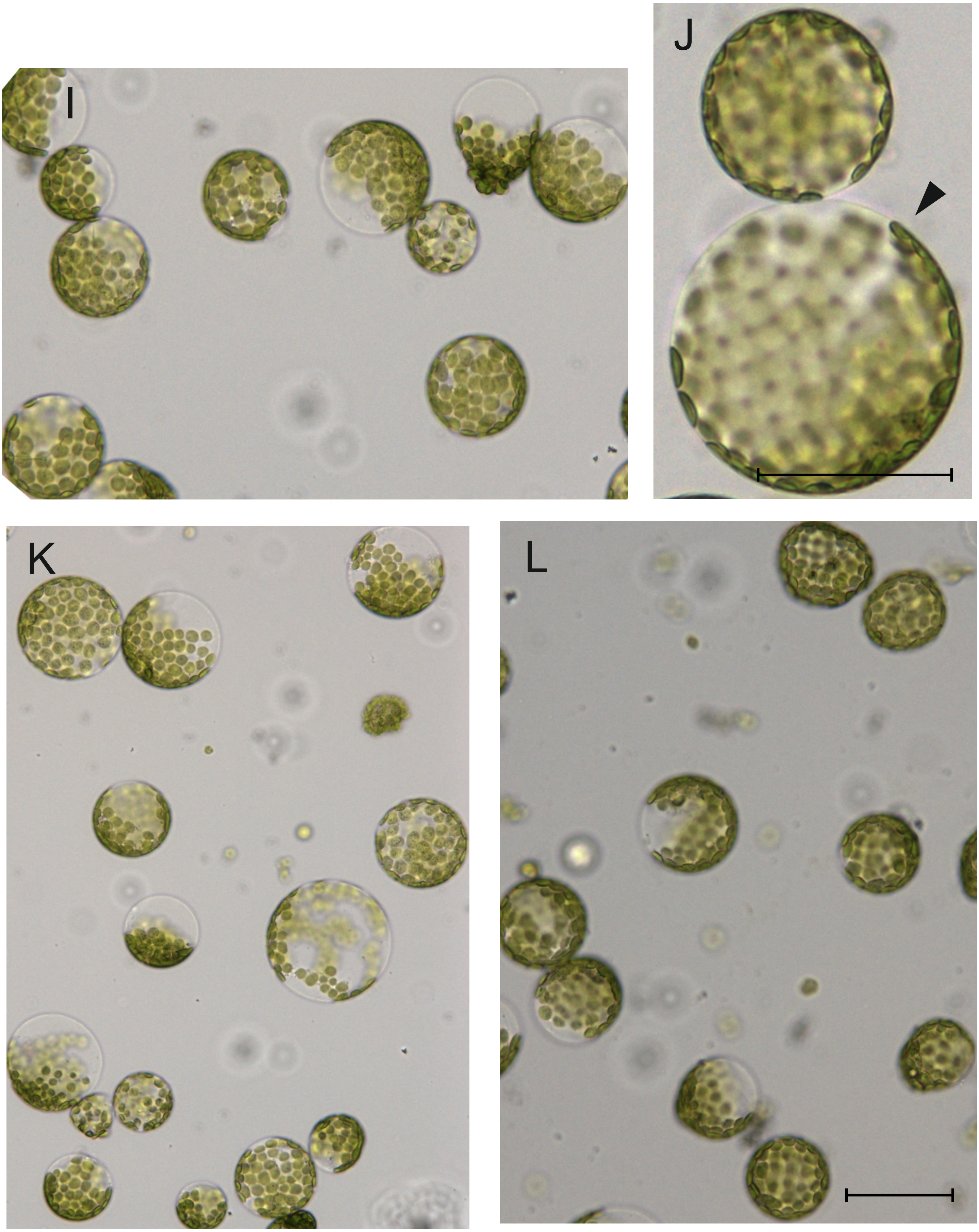
I
J
K
L
